# Genetic, cellular and structural characterization of the membrane potential-dependent cell-penetrating peptide translocation pore

**DOI:** 10.1101/2020.02.25.963017

**Authors:** Evgeniya Trofimenko, Gianvito Grasso, Mathieu Heulot, Nadja Chevalier, Marco A. Deriu, Gilles Dubuis, Yoan Arribat, Marc Serulla, Sébastien Michel, Gil Vantomme, Florine Ory, Linh Chi Dam, Julien Puyal, Francesca Amati, Anita Lüthi, Andrea Danani, Christian Widmann

## Abstract

Cell-penetrating peptides (CPPs) allow intracellular delivery of cargo molecules. They provide efficient methodology to transfer bioactive molecules in cells, in particular in conditions when transcription or translation of cargo-encoding sequences is not desirable or achievable. The mechanisms allowing CPPs to enter cells are ill-defined. Using a CRISPR/Cas9-based screening, we discovered that KCNQ5, KCNN4, and KCNK5 potassium channels positively modulate cationic CPP direct translocation into cells by decreasing the transmembrane potential (V_m_). These findings provide the first unbiased genetic validation of the role of Vm in CPP translocation in cells. *In silico* modeling and live cell experiments indicate that CPPs, by bringing positive charges on the outer surface of the plasma membrane, decrease the V_m_ to very low values (−150 mV or less), a situation we have coined megapolarization that then triggers formation of water pores used by CPPs to enter cells. Megapolarization lowers the free energy barrier associated with CPP membrane translocation. Using dyes of varying sizes, we assessed the diameter of the water pores in living cells and found that they readily accommodated the passage of 2 nm-wide molecules, in accordance with the structural characteristics of the pores predicted by *in silico* modeling. Pharmacological manipulation to lower transmembrane potential boosted CPPs cellular internalization in zebrafish and mouse models. Besides identifying the first proteins that regulate CPP translocation, this work characterized key mechanistic steps used by CPPs to cross cellular membrane. This opens the ground for strategies aimed at improving the ability of cells to capture CPP-linked cargos *in vitro* and *in vivo*.

## Introduction

Cell penetrating peptides (CPPs) are short non-toxic sequences of 5-30 amino acids present in proteins able to cross membranes such as homeoproteins and some viral components. CPPs can also be used to deliver bioactive cargo (siRNAs, DNA, polypeptides, liposomes, nanoparticles and others) in cells for therapeutic or experimental purposes^1–11^. Even though they differ in their origin^12–15^ and physico-chemical properties, the majority of CPPs carry positive charges in their sequence^1, 3, 6, 7^. Poly-arginine (e.g. R9), HIV-1 TAT_47-57_, Penetratin (Antennapedia_43-58_), and Transportan are among the most used and studied CPPs.

CPPs enter cells through a combination of two non-mutually exclusive mechanisms^9, 10^: endocytosis and direct translocation^1–8, 11^. The nature of these entry mechanisms is debated and not fully understood at the molecular level. The vesicular internalization of CPPs may occur through clathrin-dependent endocytosis, macropinocytosis, and caveolin-1-mediated endocytosis^1, 3–8^. In this case, access to the cytoplasm requires that the CPPs break out of endosomes through a poorly understood process called endosomal escape.

Direct translocation allows the CPPs to access the cytosol through their ability to cross the plasma membrane. There is currently no unifying model to explain mechanistically how direct translocation proceeds and no genes have yet been identified to modulate the manner by which CPPs cross cellular membranes. There is a general consensus though that an adequate plasma membrane potential (V_m_) is required for direct translocation to occur. Electrophysiological and pharmacological V_m_ modulations have revealed that depolarization blocks CPP internalization^16, 17^ and hyperpolarization improves the internalization of cationic CPPs^16, 18–21^. By itself, a sufficiently low V_m_ (i.e. hyperpolarization) appears to trigger CPP direct translocation in live cells^16–18^. *In silico* modeling has provided evidence that membrane hyperpolarization leads to the formation of transient water pores, allowing CPP translocation into cells^22–26^ but the free energy landscape governing CPP translocation has not been determined. Moreover, the nature and the structural characteristics of the pores used by CPPs to cross the plasma membrane have not been investigated in live cells.

Here, we provide the first genetic evidence that validates the importance of V_m_ for CPP direct translocation and we characterize the size of the water pores used by CPPs to enter live cells. We also determined the role of the V_m_ in modulating the free energy barrier associated with membrane translocation and the impact of the V_m_ on CPP translocation kinetics.

## Results

### TAT-RasGAP_317-326_ enters cells via endocytosis and direct translocation, but only direct translocation mediates its biological activity

In the present work, we have used TAT-RasGAP_317-326_ as a model compound to investigate the molecular basis of CPP cellular internalization. This peptide is made up of the TAT_48-57_ CPP and a 10 amino-acid sequence derived from the SH3 domain of p120 RasGAP^27^. TAT-RasGAP_317-326_ sensitizes cancer cells to chemo-, radio- and photodynamic therapies^28–31^ and prevents cell migration and invasion^32^. This peptide also exhibits antimicrobial activity^33^. Some cancer cell lines, such as Raji (Burkitt’s lymphoma), SKW6.4 (transformed B-lymphocytes) and HeLa (cervix carcinoma), are directly killed by this peptide in a necrotic-like manner^34^. TAT-RasGAP_317-326_ kills cells by physical disruption of the plasma membrane once it has reached the cytosol by direct translocation and interacted with specific phospholipids found on the cytosolic side of the plasma membrane^35^. To characterize the mode of entry of TAT-RasGAP_317-326_, Raji, SKW6.4 and HeLa cells were incubated with a FITC-labelled version of the peptide and visualized over time. Peptide-cell interaction and internalization occurred through the following pattern: binding to the plasma membrane (Supplementary Movie 1), internalization leading to vesicular staining and/or diffuse cytosolic staining, and eventually cell death characterized by membrane disruption (Supplementary Fig. 1a-d and Supplementary Movie 1). Only in a minority of Raji and SKW6.4 cells, was the peptide solely found in vesicular structures (i.e. in the absence of cytosolic staining) (Supplementary Fig. 1b). Most of the cells acquired the peptide rapidly in their cytoplasm without any detectable presence of labelled vesicles (Supplementary Fig. 1a-b and Supplementary Movies 1-2). Even though labelled vesicles might be masked by the intense cytosolic staining (when present), these results nevertheless demonstrate that at the concentrations used here the main mode of entry of TAT-RasGAP_317-326_ in Raji and SKW6.4 cell lines occurs via a non-vesicular direct plasma membrane translocation mechanism. This entry can be visualized in living cells prior to the saturation of the cytosolic signal (Supplementary Movies 1-2). In the case of TAT-RasGAP_317-326_ uptake in HeLa cells, direct translocation and the vesicular mode of entry appeared to occur equally frequently (Supplementary Fig. 1a-b and Supplementary Movie 3). Direct translocation across the plasma membrane often seemed to originate from specific areas of the cells (Supplementary Fig. 1e-f), suggesting discrete structures on the plasma membrane involved in CPP entry^18, 36–39^. Laser illumination-induced damage to cell integrity^36–39^ was ruled out as a mechanism of fluorescent dye-labelled CPP cellular internalization (Supplementary Fig. 1g). Cells were able to concentrate the CPP in their cytoplasm (Supplementary Fig. 1h). TAT-RasGAP_317-326_ internalization was found to be temperature-dependent (Supplementary Fig. 1i). The contribution of the peptide surface binding to the overall cell-associated signal only accounted to about 20% (Supplementary Fig. 1j-k); hence, the majority of the cell-associated peptide signal comes from within cells. Despite intensity differences, the kinetics of appearance of the two populations with vesicular and diffuse cytosolic staining was similar (Supplementary Fig. 2a-b). There was an almost perfect correlation between i) the percentage of cells with low intensity peptide fluorescence and the percentage of cells with predominant vesicular staining (Supplementary Fig. 2c), ii) the percentage of cells with high intensity peptide fluorescence and the percentage of cells with marked cytosolic staining (Supplementary Fig. 2d). Cell death induced by TAT-RasGAP_317-326_ was mostly recorded in cells from the high-intensity fluorescent population (Supplementary Fig. 2e), which can only be achieved through direct translocation of the peptide through the cell membrane as cell death was not seen in cells where only vesicular CPP entry had occurred. Hence, direct translocation of the peptide is what allows TAT-RasGAP_317-326_ to kill HeLa cells. In Raji and SKW6.4 cells incubated with TAT-RasGAP_317-326_, only one fluorescence intensity peak was observed at a given time point that progressively shifted towards higher fluorescence intensities (Supplementary Fig. 2f). As little vesicular staining was detected in Raji and SKW6.4 cells incubated with TAT-RasGAP_317-326_ (Supplementary Fig. 1a-b), the peak seen in Supplementary Fig. 2f mostly corresponds to cells that have acquired the peptide through direct translocation. As in HeLa cells therefore, it is the direct translocation of the peptide in Raji and SKW6.4 cells that eventually induces their death. Together these results show that even though two modes of entry can be used by TAT-RasGAP_317-326_ to penetrate cells, it is direct translocation that is necessary for its biological activity.

### Identification of potassium channels as mediators of CPP direct translocation into cells

The mode of CPP cellular entry is still debated and no proteins have been identified that regulate this process. The CPP entry process starts after the initial electrostatic interactions between the positively charged CPP and the negatively charged components of the cell membrane^1–8, 11^. Interaction with acid sphingomyelinase^40^, local membrane deformation^41^, as well as calcium fluxes^42^ have been suggested to play a role in CPP direct translocation into cells. To screen for genes that control CPP internalization, we performed a CRISPR/Cas9 screen using TAT-RasGAP_317-326_ as a selective agent (Supplementary Fig. 3a). In Raji and SKW6.4 cell lines, among the most highly significantly differentially modulated genes were specific potassium channels or genes coding for proteins known to regulate such channels indirectly (e.g. PIP5K1A^43^) (Fig. 1a and Supplementary Fig. 3b). KCNQ5, identified in Raji cells, is a voltage-dependent potassium channel. KCNN4 and KCNK5, identified in SKW6.4 cells, are calcium-activated channels and belong to the two-pore (voltage-independent) potassium channel family^44^, respectively.

**Fig. 1:**
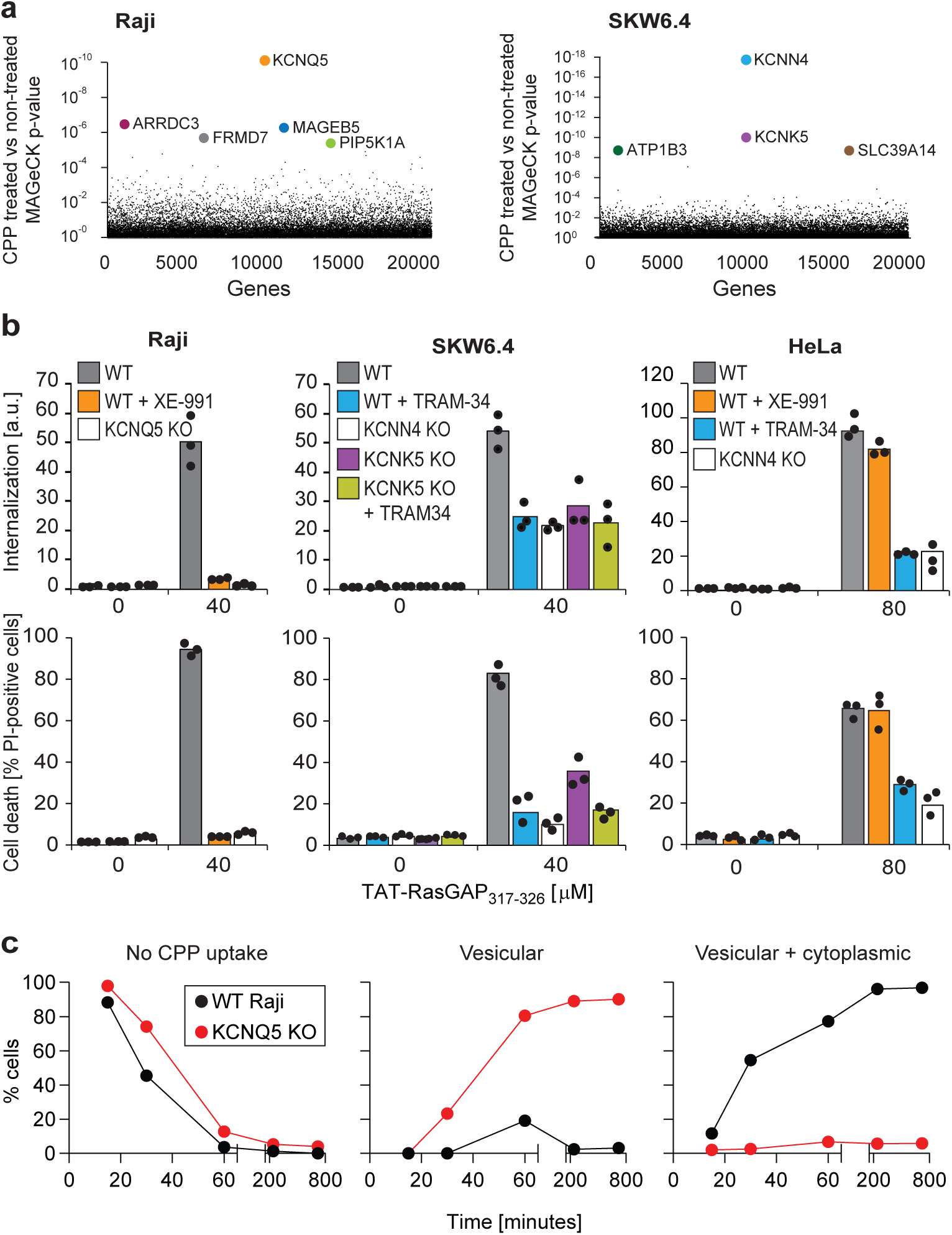
Identification of potassium channels as mediators of direct translocation of CPPs into cells. **a**, Identification of genes implicated in TAT-RasGAP_317-326_ internalization in Raji and SKW6.4 cells. The graphs depict the p-value (calculated using the MAGeCK procedure; see Materials and Methods) for the difference in sgRNA expression between peptide-treated and control cells for the ∼20’000 genes targeted by the CRISPR/Cas9 library. **b**, Quantitation of TAT-RasGAP_317-326_ entry (top) and induced death (bottom) in wild-type (WT) and knock-out (KO) cells. The WT and the corresponding potassium channel KO versions of the indicated cell lines were pretreated or not for 30 minutes with 10 μM XE-991 or with TRAM-34 and then incubated (still in the presence of the inhibitors when initially added) with, or without 40 μM (Raji and SKW6.4 cells) or 80 μM (HeLa cells) TAT-RasGAP_317-326_. Internalization was recorded after one hour and cell death after 16 hours (Raji and SKW6.4) or 24 hours (HeLa). Results correspond to the average of three independent experiments. TAT-RasGAP_317-326_ concentrations and time of incubation used were adjusted so that the CPP induced similar cell death (between 60% and 90%) in the wild-type versions of the different cell lines. **c**, Quantitation of the modalities of TAT-RasGAP_317-326_ entry in wild-type and KCNQ5 knock-out Raji cells. Cells were incubated with FITC-TAT-RasGAP_317-326_ for various periods of time and peptide staining was visually quantitated on confocal images (n>150 cells for each time-point). The high percentage of cells with vesicular staining in the knock-out (KO) cells results from the absence of strong diffuse staining masking endosomes. The results correspond to the average of three experiments.

These potassium channels were pharmacologically or genetically inactivated (Fig. 1b and Supplementary Fig. 4a-c) to validate their involvement in TAT-RasGAP_317-326_ cellular internalization and the resulting death induction. The KCNQ family inhibitor, XE-991^45^, fully blocked peptide internalization in Raji cells and protected them from the killing activity of the peptide (Fig. 1b). Knocking out KCNQ5 (Supplementary Fig. 4a) in these cells produced the same effect as XE-991 (Fig. 1b and Supplementary Fig. 4d). SKW6.4 cells individually lacking KCNN4 or KCNK5 (Supplementary Fig. 4b), or SKW6.4 cells treated with TRAM-34, a KCNN4 inhibitor^46, 47^ were impaired in their ability to take up the peptide and were partially protected against its cytotoxic activity (Fig. 1b and Supplementary Fig. 4d). Inhibition of KCNN4 activity with TRAM-34 in KCNK5 knock-out cells did not further protect the cells against TAT-RasGAP_317-326_-induced death. In HeLa cells, TRAM-34, but not XE-991, inhibited TAT-RasGAP_317-326_ internalization and subsequent death (Fig. 1b). Thus, in HeLa cells, KCNN4 channels participate in the cellular internalization of the peptide. This was confirmed by knocking out KCNN4 in these cells (Fig. 1b and Supplementary Fig. 4c). Resistance to TAT-RasGAP_317-326_-induced death in KCNQ5 knock-out Raji cells and KCNN4 knock-out SKW6.4 or HeLa cells was restored through ectopic expression of the corresponding FLAG- or V5-tagged channels (Supplementary Fig. 4e-f and Supplementary Fig. 5), ruling out off-target effects.

To determine if the identified potassium channels affected the biological activity of TAT-RasGAP_317-326_ once inside cells, incubation with the peptide was performed in the presence of pyrene butyrate, a counter anion that, through electrostatic interaction with arginine residues (Supplementary Fig. 6a), promotes quick and efficient translocation of arginine-bearing peptides into cells^48^. Pyrene butyrate, to promote peptide uptake, has to be used in the absence of serum^48^. But since serum removal sensitized cells to TAT-RasGAP_317-326_-induced death (Supplementary Fig. 6b), we had to adapt the CPP concentrations to perform the experiments that employed pyrene butyrate (Supplementary Fig. 6c). Pyrene butyrate was not toxic at the concentrations used (Supplementary Fig. 6d). Potassium channel knock-out Raji and HeLa cells incubated with both pyrene butyrate and TAT-RasGAP_317-326_ were no longer resistant against peptide-induced death (Supplementary Fig. 6e). These results demonstrate that specific potassium channels play crucial role in the cellular entry of TAT-RasGAP_317-326_ but that they are not involved in the killing activity of the peptide if it is already in cells.

We next determined whether vesicular internalization or direct translocation were affected in cells with impaired potassium channel activities. Compared to their respective wild-type controls, the percentage of cells with diffuse cytosolic location of FITC-TAT-RasGAP_317-326_ was drastically diminished in KCNQ5 knock-out Raji cells (Fig. 1c) and in KCNN4 and KCNK5 knock-out SKW6.4 cells (Supplementary Fig. 7a). The effect was similar in the case of KCNN4 knock-out HeLa cells, albeit to a lesser extent (Supplementary Fig. 7b). This was mirrored by an increase in the percentage of knock-out cells with vesicular staining (Fig. 1c and Supplementary Fig. 7a-b). To assess whether cytosolic peptide accumulation is due to direct translocation and not endosomal escape, FITC-TAT-RasGAP_317-326_ internalization was quantitated over time in a condition where the peptide was allowed to enter cells by endocytosis but then removed from the medium. Supplementary Fig. 7c (left) shows that diffuse cytosolic internalization of the peptide occurred when the peptide was present in the medium (see also left panel of Supplementary Movie 4). However, no significant increase in the cytosolic signal was observed in washed out cells containing peptide-labelled endosomes (Supplementary Fig. 7c, middle and Supplementary Movie 4, right panel). We could nevertheless detect an increase in cytosolic fluorescence when endosomal escape was induced by LLOME^49^. This confirms our earlier observation (Supplementary Fig. 1 and Supplementary Movie 1-2) that in live cell conditions the diffuse cytosolic CPP signal originates from direct translocation and not endosomal escape. The invalidation of potassium channels did not affect transferrin internalization into cells (Supplementary Fig. 7d) or the infectivity of vesicular stomatitis virus^50^, substantiating the non-involvement of these channels in endocytosis pathways.

One possibility to explain the above-mentioned results is that the absence of potassium channels prevents the peptide binding to cells. At a 20 μM concentration, TAT-RasGAP_317-326_ is readily taken up by wild-type Raji cells but not by KCNQ5 knock-out cells. At this concentration, peptide binding was slightly lower in knock-out than in wild-type cells (Supplementary Fig. 7e). However, augmenting the peptide concentrations in the extracellular medium of KCNQ5 knock-out cells to reach surface binding signals equivalent or higher than what was obtained in wild-type cells incubated with a 20 μM peptide concentration still did not result in peptide cellular internalization (Supplementary Fig. 7e). These data show that differences in peptide binding do not explain the inability of the knock-out cells to take up the peptide.

We assessed whether the role of potassium channels in cellular internalization also applied to TAT cargos other than RasGAP_317-326_. TAT-PNA is an oligonucleotide covalently bound to TAT, which can correct a splicing mutation within the luciferase-coding sequence^51, 52^. This can only occur if TAT-PNA reaches the cytosol. The luciferase activity triggered by TAT-PNA was diminished in the presence of potassium channel inhibitors and in potassium channel knock-out cell lines (Supplementary Fig. 8a). Cytosolic access of TAT-Cre, which can recombine a loxP-RFP-STOP-loxP-GFP^53, 54^ gene construct, was then assessed. Switch from red to green fluorescence occurs only when TAT-Cre reaches the nucleus. This took place in wild-type Raji cells but not in the KCNQ5 knock-out cells (Supplementary Fig. 8b). We finally tested a clinical phase III therapeutic D-JNKI1 compound^1, 2^ used in the context of hearing loss and intraocular inflammation. The internalization of this peptide was completely blocked in Raji cells lacking KCNQ5 (Supplementary Fig. 8c, left). D-JNKI1 internalization was also diminished in SKW6.4 cells lacking KCNN4 and KCNK5 channels, as well as in HeLa cells lacking KCNN4 potassium channel (Supplementary Fig. 8c, middle and right panels). These data demonstrate that the absence of specific potassium channels diminishes or even blocks the entry of various TAT-bound cargos.

### Potassium channels maintain plasma membrane polarization that is required for CPP entry into cells

Potassium is the main ion involved in setting the plasma membrane potential (V_m_). The potassium channels identified in the CRISPR/Cas9 screen may therefore participate in the establishment of the cell V_m_. Figure 2a shows that genetic disruption or pharmacological inhibition of KCNQ5 in Raji cells led to an increase in their V_m_ (from -26 mV to -15 mV, validated with electrophysiological recordings; see Supplementary Fig. 9a-b). Surprisingly, such minimal increase in V_m_ l between the wild-type and the respective knock-out cells practically abolished CPP internalization (Fig. 2b), indicating that above a certain threshold, the V_m_ l is no longer permissive for CPP direct translocation. In SKW6.4 and HeLa cells, V_m_ measurement was much more variable than in Raji cells. Nevertheless, a trend of increased V_m_ was observed when KCNN4 or KCNK5 were invalidated genetically or pharmacologically (Fig. 2a). As the CRISPR/Cas9 screens performed in various cell lines identified a variety of potassium channels required for efficient CPP internalization, we conclude that it is the V_m_ maintenance activity of these channels that is important for CPP direct translocation and not some specific features of the channels.

**Fig. 2:**
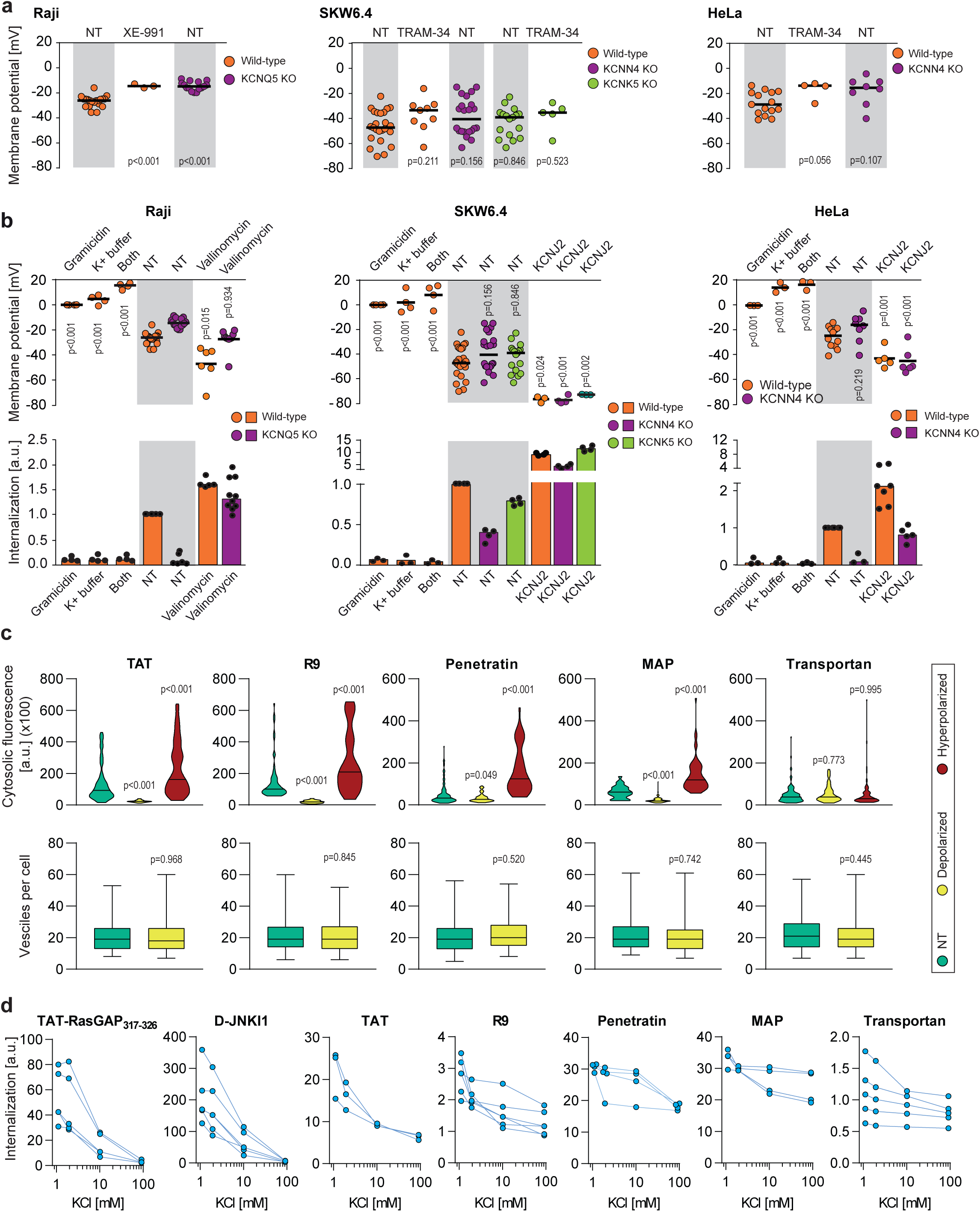
Potassium channels maintain plasma membrane polarization that is required for CPP entry into cells. **a**, Assessment of the resting plasma membrane potential in the indicated wild-type cell lines and the corresponding potassium channel knock-out (KO) clones in the presence or in the absence 10 μM XE-991 or TRAM-34. The grey and white zones correspond to non-treated cells and inhibitor-treated cells, respectively. NT, not treated. The p-values correspond to the assessment of the significance of the differences with the control wild-type condition using ANOVA multiple comparison analysis with Dunnett’s correction. Each dot in a given condition represents an independent experiment. **b**, Effect of cellular depolarization (left of the grey zone) and hyperpolarization (right of the grey zone) on peptide internalization in the absence of serum. The indicated cell lines and the corresponding channel knock-out (KO) clones were pretreated or not with depolarization agents (2 μg/ml gramicidin for 5 minutes or high extracellular potassium buffer for 30 minutes) or with hyperpolarization inducer (10 μM valinomycin), followed by the addition of TAT-RasGAP_317-326_ for one hour. Alternatively, hyperpolarization was achieved by ectopic expression of the KCNJ2 potassium channel. Membrane potential and peptide internalization were then determined. Membrane potential was measured in the presence of DiBac4(3) by flow cytometry. Peptide internalization was measured by flow cytometry in the presence of 0.2% trypan blue. The p-values correspond to the assessment of the significance of the differences with the control wild-type condition using ANOVA multiple comparison analysis with Dunnett’s correction. Each dot in a given condition represents an independent experiment. **c**, Quantitation of cytosolic CPP signal (top) and the number of endocytic vesicles per cell (bottom) in wild-type HeLa cells (n>150 cells) incubated for one hour with 10 μM FITC-CPP in depolarizing (2 μg/ml gramicidin) or hyperpolarizing (10 μM valinomycin) conditions in the absence of serum based on confocal microscopy images (see Supplementary Fig. 9g). Comparison between different conditions to non-treated control was done using ANOVA test with Dunnett’s correction for multiple comparison. The number of endocytic vesicles per cell was quantitated based on confocal images. Statistical comparison was done using t-tests. Quantitation of vesicles was not performed in hyperpolarizing conditions due to masking from strong cytosolic signal. The confocal images were acquired in the middle of the cells based on Hoechst fluorescence. **d**, Internalization of various CPPs in the presence of different concentrations of potassium chloride in the media. Data for a given experiment are linked with thin blue lines.

If the reason why invalidation of the KCNQ5, KCNN4, and KCNK5 potassium channels inhibits TAT-RasGAP_317-326_ cellular entry is cell depolarization, a similar response should be obtained by chemically inducing membrane depolarization. Figure 2b shows that depolarizing cells with gramicidin^55^ (making non-specific 0.4 nm pores^56^ within cell membranes) or by increasing the extracellular concentration of potassium (dissipating the potassium gradient) totally blocked peptide entry into the three studied cell lines. Hence, cellular depolarization in itself inhibits TAT-RasGAP_317-326_ cellular internalization (Fig. 2b and Supplementary Fig. 9c). Next, we determined whether hyperpolarization could reverse the inability of potassium channel knock-out cells to take up TAT-RasGAP_317-326_. Cells were either incubated in the presence of valinomycin^57^, which leads to formation of potassium-like channels, or transfected with KCNJ2 channel that also provokes potassium efflux and membrane hyperpolarization^58^. Figure 2b shows that in Raji cells, treatment with valinomycin was able to hyperpolarize KCNQ5 knock-out cells to a V_m_ value similar to the one found in wild-type cells. This fully restored peptide translocation in the KCNQ5 knock-out cells. Hyperpolarization in SKW6.4 and HeLa cells was achieved through KCNJ2 ectopic expression. This reversed the poor capacity of the KCNN4 and KCNK5 knock-out SKW6.4 cells, as well as KCNN4 knock-out HeLa cells, to take up TAT-RasGAP_317-326_ (Fig. 2b). Moreover, hyperpolarization increased peptide internalization in wild-type cells (Fig. 2b). Additionally, cells such as primary rat cortical neurons that naturally have a low V_m_ (−48 mV) take up the CPP in their cytosol more efficiently than cells with higher V_m_ such as HeLa cells (−25 mV) (Supplementary Fig. 9d). Altogether, these results demonstrate that the V_m_ modulates internalization of TAT-RasGAP_317-326_ in various cell lines. This internalization can be manipulated through cellular depolarization to block it and through hyperpolarization to increase it, confirming earlier results obtained for the R8 CPP in Jurkat cells^16^.

Next, we assessed whether the entry of different classes of CPPs was also regulated by the membrane potential. The most commonly used CPPs in biology and medicine, TAT, nanomeric arginine (R9), Penetratin, MAP, and Transportan were tested in depolarized and hyperpolarized conditions. The cytosolic signal of these CPPs in HeLa cells appeared to originate from both vesicular and diffuse cytosolic staining (Supplementary Fig. 9e-g). With the notable exception of Transportan, depolarization led to decreased cytosolic fluorescence of all CPPs, while hyperpolarization favored CPP translocation in the cytosol (Fig. 2c, Supplementary Fig. 9g and 10a). Transportan, unlike the other tested CPPs, enters cells predominantly through endocytosis (Supplementary Fig. 9e), which could explain the difference in response to V_m_ modulation. The number of CPP-positive vesicles per cell was similar in normal and depolarized conditions, further confirming that CPP endocytosis is not affected by V_m_ (Fig. 2c, bottom panels). Next, we assessed if modulating V_m_ through changes in potassium concentrations in the extracellular medium (Supplementary Fig. 10b) impacted CPP internalization. Although the extent of the response varied from one CPP to another, the global observation was that the more depolarized the cells are, the less able they are to take up CPPs (Fig. 2d). CPP membrane binding was only minimally affected by depolarization (Supplementary Fig. 10c). The reason why depolarized cells do no take up CPPs is therefore not a consequence of reduced CPP binding to cells. The V_m_ also controlled peptide internalization in non-transformed rat primary cortical neurons (Supplementary Fig. 10d). Together these results show that CPP internalization can be modulated in vitro by varying the V_m_ of cells.

Extracellular pH might regulate CPP internalization^59^. However, in our experimental system, appearance of TAT-RasGAP_317-326_ in the cytosol through direct translocation was unaffected by varying the extracellular pH, whether or not V_m_ modulators were used (Supplementary Fig. 10e-f).

### CPP direct translocation modeling

To further study the mechanism of CPP cellular entry through direct translocation, we took advantage of coarse-grained molecular dynamics technique and MARTINI force field^60, 61^. In our simulations we have used TAT-RasGAP_317-326_, TAT, R9, Penetratin, MAP, and Transportan in presence of a natural cell membrane-like composition (for both inner and outer leaflets) while earlier studies have employed simpler membrane composition^17, 20, 22– 24, 26^. Membrane hyperpolarization was achieved by setting an ion imbalance^24, 62–64^ through a net charge difference of 30 positive ions (corresponding to a V_m_ of ∼2V) between the intracellular and extracellular space. This protocol allowed us to observe CPP translocation across membranes within a few tens of nanoseconds (Fig. 3a-b, Supplementary Fig. 11a and Supplementary Movie 5).

**Fig. 3:**
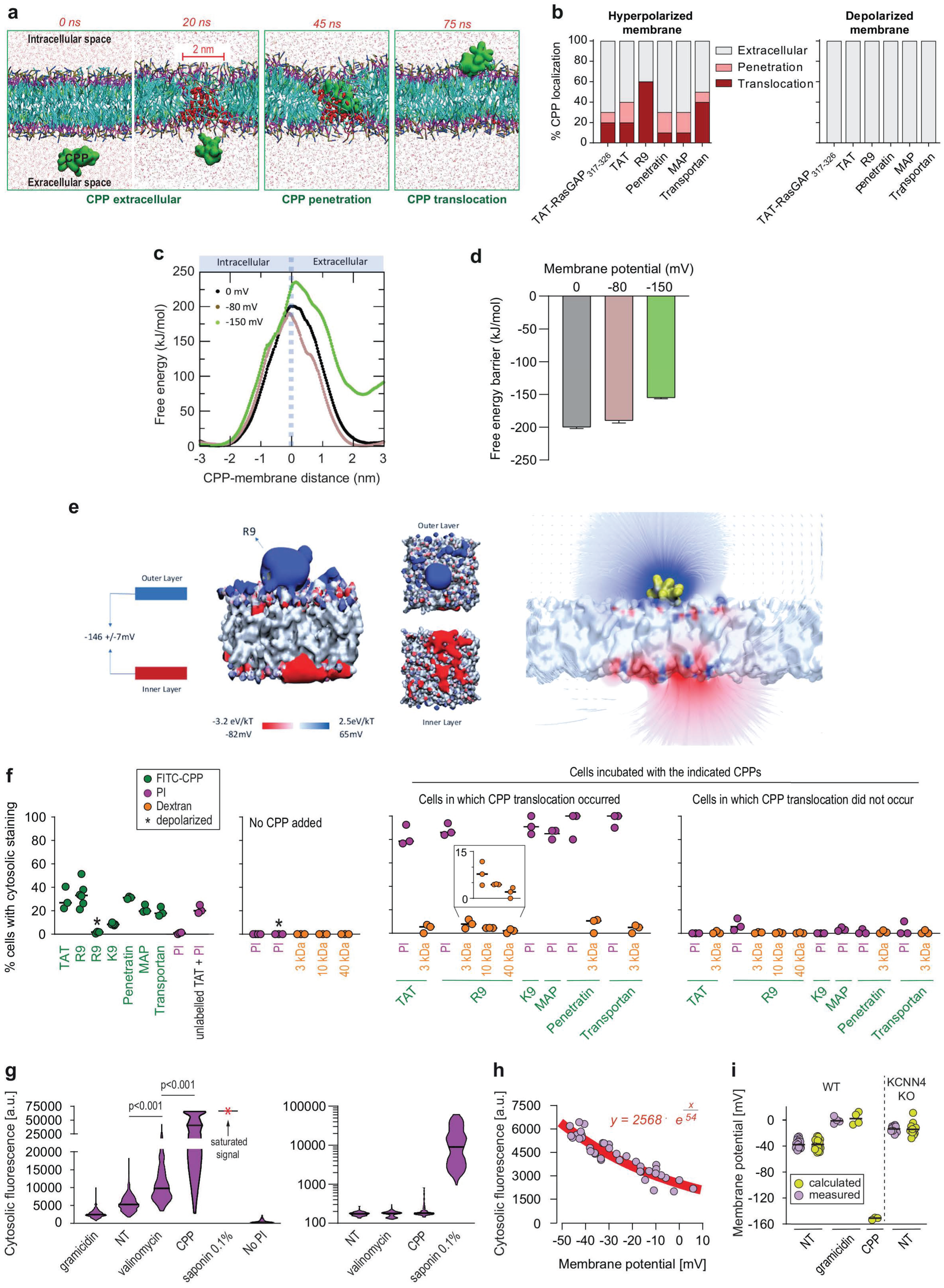
Hyperpolarization favors the formation of ∼2 nm-wide water pores used by CPPs to translocate into cells. **a**, Visualization of *in silico* modelled, time-dependent, TAT-RasGAP_317-326_ penetration and subsequent translocation across cellular membrane through a water pore. Water molecules within membranes are depicted by red spheres (and by red dots outside the membrane). **b**, Quantitation of CPP localization in hyperpolarized or depolarized conditions based on coarse-grained molecular dynamics simulations. Membrane hyperpolarization was achieved through a net charge difference of 30 positive ions between intracellular and extracellular space in a double bilayer system^24, 71–73^ obtaining a transmembrane potential of -2.2 V. Such low membrane potential was required to visualize translocation within the time frame of the simulations (100 nanoseconds). **c,** Free energy landscape of R9 translocation reported as a function of CPP-membrane distance. The metadynamics simulations have been performed at transmembrane potential values of 0 mV, -80 mV, and -150 mV (black, brown, and green curve). **d**, Free energy barrier for CPP translocation at different transmembrane potential values. **e**, Electrostatic potential maps of molecular systems that contain or not one R9 peptide in contact with the cell membrane, without any applied external electrostatic field. **f**, Quantitation of the percentage of cells with cytosolic staining after the indicated treatment. The indicated compounds (32 μg/ml PI, 200 μg/ml dextran, 40 μM CPP, except MAP, which was used at 20 μM) were incubated for 30 minutes on HeLa cells. Depolarization, indicated by an asterisk, was induced with 2 μg/ml gramicidin. The percentage of cells displaying cytosolic internalization of the indicated molecules was then determined on confocal images (n>200 cells; see Supplementary Fig. 12a-b). Inset corresponds to an enlargement of the percentage of cells positive for dextran in the presence of R9. The results correspond to at least three independent experiments. **g**, Left graph: quantitation of PI cytosolic internalization in wild type HeLa cells after 30 minutes of incubation in normal, depolarizing (2 μg/ml gramicidin) or hyperpolarizing (10 μM valinomycin) conditions in the presence or in the absence of 40 μM FITC-R9. Cytosolic internalization was quantitated from confocal images using ImageJ (n>300 cells; see the methods and Supplementary Fig. 12b). Right graph: as in left graph, but using lower laser power to avoid saturation of the signal obtained in saponin-permeabilized cells. The p-values correspond to the assessment of the significance of the differences with the non-treated (NT) control condition using ANOVA multiple comparison analysis with Dunnett’s correction. The results correspond to three independent experiments. **h**, Relation between cytosolic PI cytosolic internalization and membrane potential measured with the DiBac4(3) sensor in HeLa cells. Each dot represents an independent experiment. **i**, The relation between PI cytosolic internalization and membrane potential from panel e was used to calculate membrane potential based on PI fluorescence in HeLa cells and its corresponding KCNN4 KO. Measured membrane potential was acquired in the presence of DiBac4(3) based on its cytosolic fluorescence. Each dot in a given condition represents an independent experiment.

In presence of the ∼2V V_m_, the CPPs approached the membrane on the extracellular side and this led to the formation of a water column within the membrane that the CPP then used to move to the intracellular space (Supplementary Movie 5). The movement of the positive charges carried by the CPPs, as well as extracellular cations, to the intracellular compartments via the water pore induced membrane depolarization. This depolarization provoked the collapse of the water pore and membrane resealing. Even though CPPs play an active role in their internalization, the mere presence of the CPP in the absence of a sufficiently low V_m_ was not sufficient to trigger water pore formation (Fig. 3b and Supplementary Movie 6). These data confirm earlier work describing the role of the V_m_ in CPP penetration into or through bilipidic membranes^17, 20, 22–26^. Differences in the physico-chemical properties of CPPs not only influenced the tendency of each peptide to translocate into the intracellular compartment (Fig. 3b), but also modified the time needed for water pore formation at a given V_m_ (Supporting Figure 11b). The more positively charged a CPP (e.g. R9 peptide), the higher probability to translocate across cell membranes (Fig. 3c-d) and faster kinetics of water pore formation at a given V_m_. As usually done in the computational field, we used very high V_m_ values in order to capture ns occurring events. To estimate the time needed for CPPs to translocate through membranes at -150 mV, we established the relationship between the time of CPP translocation and the V_m_ in the Volt range used during the simulation runs (Supplementary Fig. 11b) and extrapolated from this relationship the time needed for CPP translocation at a V_m_ of -150 mV (Supplementary Fig. 11b). Even though this extrapolation is likely to lack accuracy because of the well-known limitation of the MARTINI forcefield in describing the absolute kinetics of the molecular events, the values obtained are consistent with the kinetics of CPP direct translocation observed in living cells (Figure 1c and Supplementary Fig. 1b and 9e). With the exception of Transportan, the estimated CPP translocation occurred within minutes. This is consistent with our observation that Transportan enters cells predominantly through endocytosis and its internalization is therefore not affected by changes in V_m_ (Fig 2c-d and Supplemental Fig. 9e).

We also applied a metadynamics protocol to estimate the impact of the V_m_ on the free energy landscape of R9 translocation. The free energy barriers recorded in depolarized membranes (V_m_ = 0) and polarized membranes (V_m_ = -80 mV) were similar (Fig. 3c-d). The obtained value of about 200 kJ/mol is in in line with recent estimation of the free energy barrier associated with CPP tranlocation at a Vm=0^24^. Only at much lower V_m_ values (−150 mV) was a marked decreased in free energy barrier recorded. This indicates that hyperpolarization values found in resting cells (down to about -80 mV in neurons and higher in many other cells types^65^) are not more favorable than fully depolarized membranes to establish conditions favorable for the formation of water pores. It appears therefore that cells need to decrease their V_m_ to much lower values (e.g. -150 mV or lower) to reach conditions compatible with water pore formation.

This observation is in apparent contradiction with our results showing direct translocation in cells at –25 mV (Figure 2), as well as to the experiment demonstrating that CPP cytosolic internalization was more efficient in cortical neurons in comparison to less negatively charged HeLa cells (Supplementary Fig. 9d). We therefore postulate that the presence of CPPs on the cell surface induces locally a substantial voltage drop from the resting Vm. To test this assumption, we analyzed the electrostatic potential map in a molecular system composed of the R9 peptide in contact with the plasma membrane in the absence of an external electrostatic field (Fig. 3e). The results indicate that the presence of CPPs at the cell surface is sufficient to decrease locally the transmembrane potential to about -150 mV (Fig. 3e). This was not observed in the absence of the CPP.

In conclusion, our data support a model where CPPs further decrease the Vm of resting cells to very low values (equal or less than -150 mV) that are compatible with spontaneous water pore formation and that we coin megapolarization.

### Structural characterization of the pore allowing CPP entry in live cells

To evaluate the structural characteristics of the water pores triggered by CPPs in live cells, cells were co-incubated with molecules of different sizes and FITC-labelled CPPs at a peptide/lipid ratio of 0.012-0.018 (Supplementary Fig. 11c-d). These ratios correspond to the experimental conditions we are using when cells are exposed to 20-40 μM CPPs. Propidium iodide (PI), with a diameter of 0.8-1.5 nm^66^ or 3 kDa, 10 kDa, and 40 kDa dextrans, 2.3 ±0.38 nm^67^, 4.5 nm and 8.6 nm (size estimation provided by Thermofisher), respectively, were used to estimate the size of the water pores formed in the presence of CPP. While PI and CPPs efficiently co-entered cells, there was only marginal co-entry of the dextrans with the CPPs (Fig. 3f and Supplementary Fig. 12a-b). CPPs did not bind to PI (Supplementary Fig. 12c) and thus PI entry and accumulation within cells was not the result of CPP carry over. The marginal cytosolic co-internalization of dextrans was inversely correlated with their size (inset in the third panel of Fig. 3f). These data indicate that water pores triggered by CPPs allow molecules up to ∼2 nm to efficiently enter cells. These results are in line with the *in silico* prediction of the water pore size obtained by analyzing the structure of the pore at the transition state (i.e. when the CPP is crossing the cell membrane; see Figure 3a). The average diameter of the water pore corresponding to the transition state is 1.6+/-0.26 nm.

Molecules in the 2-5 nm diameter range can still use this entry route to a limited extent (of note, the Cre recombinase with a diameter of 5 nm, estimated from crystal structure (NDB:PD0003), can translocate into cells when hooked to TAT; see Supplementary Fig. 8b) but molecules with larger diameters are mostly prevented to do so.

Despite identical net positive charges (Supplementary Fig. 9f), K9 was less capable of translocating into cells compared to R9 (Fig. 3f and Supplementary Fig. 9e)^68^. This may be due to the deprotonation of K9 once in the plasma membrane (see discussion). However, in the few cases when cells have taken up K9, PI co-internalized as well (Fig. 3f). This indicates that K9 has a reduced capacity compared to R9 to trigger water pore formation but when they do, PI can efficiently translocate through the pores created by K9.

PI staining is commonly used to assess cell membrane integrity, frequently associated with cell death (see for example Supplementary Fig. 2e). This dye poorly fluoresces in solution (Supplementary Fig. 12d). However, the PI cytosolic intensity values in dead permeabilized cells are several orders of magnitude higher than those recorded after cell hyperpolarization (compare the left and right panels of Fig. 3g). Cell viability was not affected by transient treatment with the indicated CPPs, V_m_ modulating agents and PI (Supplementary Fig. 12e). These results indicate that water pore formation does not compromise cell survival.

Modelling experiments indicate that water pores are created in membranes subjected to sufficiently high (absolute values) V_m_. We therefore tested whether the mere hyperpolarization of cells (i.e. in the absence of CPPs) could trigger the translocation of PI into cells, indicative of water pore formation. Figure 3g (left) shows that the hyperpolarizing drug valinomycin significantly increased PI cell permeability. In contrast, depolarization, mediated by gramicidin, reduced PI internalization (Fig. 3g, left). Cells incubated with CPPs took up PI in their cytosol to an even greater extent than when cells were treated with valinomycin (Fig. 3g, left).

Figure 3h shows the correlation between cytosolic PI accumulation over time and V_m_. Based on this correlation, we estimated the V_m_ of cells incubated with a CPP to be in the order of -150 mV (Fig. 3i). In accordance with the modelling experiments, these data further support the notion i) that water pore formation in cells is favored by cell hyperpolarization and inhibited by depolarization and ii) that CPPs themselves^18, 69^ further contribute to the establishment of local megapolarization in the plasma membrane.

### Megapolarization improves CPP internalization *in vivo*

We investigated whether it was possible to experimentally manipulate the V_m_ to favor CPP internalization in *in vivo* situations. Systemic exposure of zebrafish embryos to valinomycin led to cell hyperpolarization (Supplementary Fig. 13a) and improved internalization of a TAT-based CPP (Supplementary Fig. 13b). This systemic treatment, while not acutely toxic, halted development (Supplementary Fig. 13c-e). However, local valinomycin injection did not affect long-term viability (Supplementary Fig. 13f) and efficiently increased CPP cellular internalization (Fig. 4a). Subcutaneous injections of valinomycin in mice induced tissue hyperpolarization (Supplementary Fig. 13g) and boosted the CPP delivery in skin cells (Fig. 4b). These results demonstrate that hyperpolarizing drugs can be used to ameliorate CPP internalization in animal tissues.

**Fig. 4:**
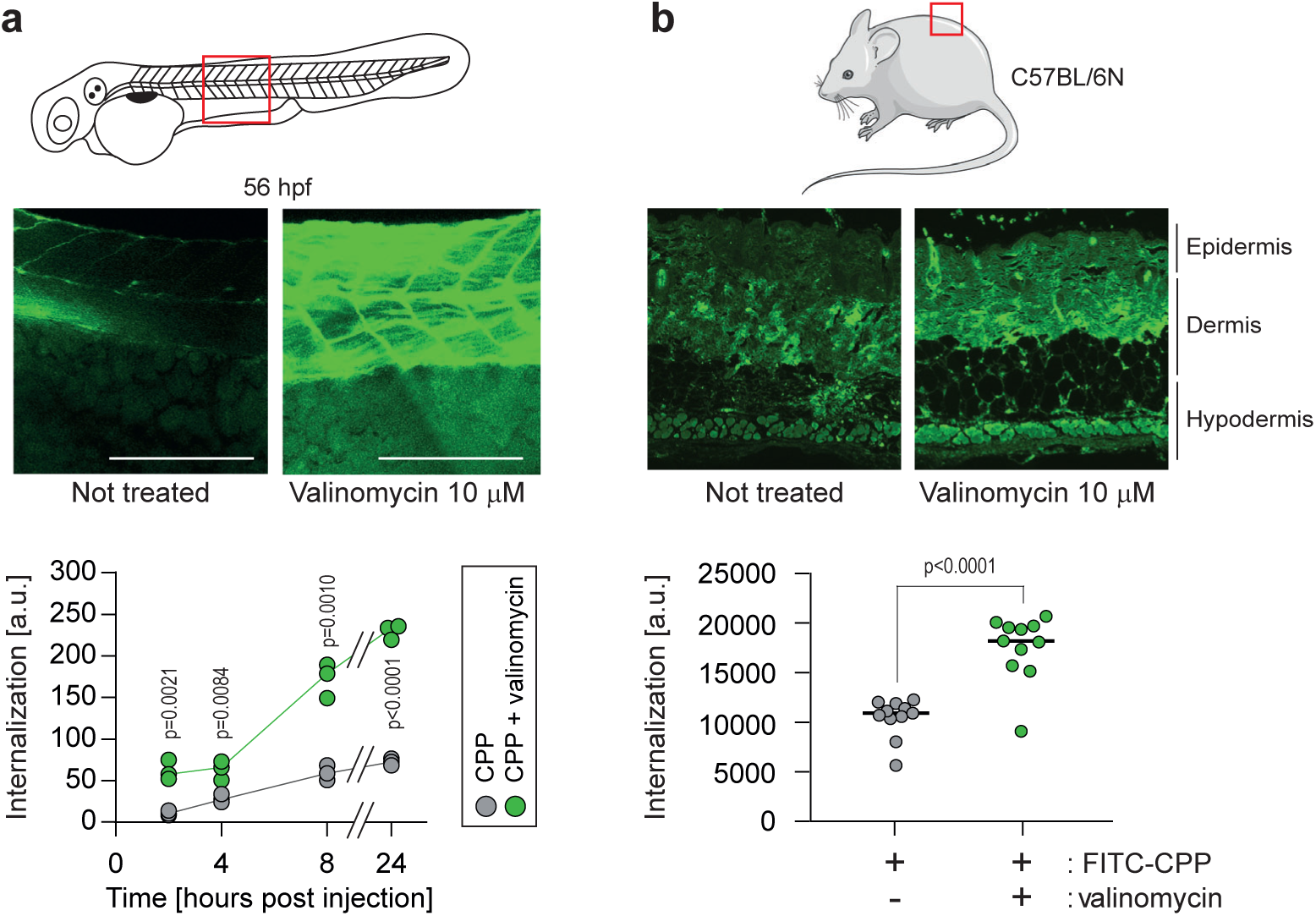
Hyperpolarization improves CPP internalization *in vivo.* **a**, CPP internalization in zebrafish embryos in normal and hyperpolarized conditions. Forty-eight-hour post fertilization, zebrafish embryos were injected with 3.12 μM FITC-TAT-RasGAP_317-326_(W317A) with or without 10 μM valinomycin. Scale bar: 200 μm. The results correspond to three independent experiments. **b**, CPP internalization in C57BL/6N mice in normal and hyperpolarized conditions. Mice were injected with 5 μM FITC-TAT-RasGAP_317-326_(W317A) with or without 10 μM valinomycin (n=11 injections per condition). In both panels, the p values associated with the comparisons of the “CPP” and “CPP + valinomycin” conditions were calculated using two-tailed paired t-tests.

## Discussion

Multiple models, mostly inferred from artificial experimental paradigms, have been proposed to explain CPP direct translocation. These include the formation of pores made of the CPPs themselves that they use for their own entry, the formation of inverted micelles in the plasma membrane that translocate the CPPs, or diffusion of the CPPs across the plasma membrane^1, 3–5, 8^. Our simulation and cellular data, while providing no evidence for such models, demonstrate that CPP cellular internalization is potassium channel-and V_m_ - dependent *in vitro* and *in vivo*. Potassium channels are required to establish a basal low V_m_, subsequently permissive for CPP direct translocation. Hyperpolarizing drugs, such as valinomycin, enhance permissiveness. When CPPs come into contact with the plasma membrane, they decrease even more the V_m_, resulting in a locally megapolarized membrane. This increases the likelihood of water pore formation that the CPPs then use to penetrate cells according to their electrochemical gradient (Supplementary Fig. 14). Water pores are created by a combination of lipid head group reorientation coupled to intrusion of a column of water in the membrane bilayer. Water movement plays therefore an active role in the formation of the pore and is not merely occurring once the pores are formed. The movement of the positive charges carried by the CPPs into the cell, as well as the transport of extracellular cations (e.g. Na^+^), dissipates the V_m_, resulting in the collapse of the water pores and sealing of the plasma membrane. CPP-mediated formation of water pores is therefore transient and does not affect cell viability. Multiple rounds of CPP-driven water pore formation and CPP translocation into cells can lead to intracellular accumulation of the CPP to concentrations higher than found outside cells (Supplementary Fig. 1h).

It has not been possible to measure directly the precise values of the V_m_ that allow the formation of water pores used by CPPs to enter cells. Using an indirect calculation mode based on the uptake of PI alongside CPPs, we have estimated that a V_m_ in the order of -150 mV is required for water pores to be formed (Figure 3h). This might be an underestimation however as modeling data indicate that, at -150 mV, the free energy barrier, while being markedly diminished compared to those calculated at -80 mV or 0 mV, is not fully abrogated (Figure 3d). Possibly therefore, the local V_m_ where CPPs interact with the plasma membranes is much lower than -150 mV.

Our model posits that the number of positively charged amino acids influence the ability of CPPs to hyperpolarize cells and hence to form water pores that they take to translocate into cells. CPP hydropathy strongly correlates with penetration of water molecules in the lipid bilayer, thus supporting the hypothesis that the amount of water each CPP can route inside the membrane is modulated by the hydrophobic and hydrophilic character of the peptide^70^. The nature of cationic amino acids in peptides determines their translocation abilities. It is known for example that peptides made of 9 lysines (K9) poorly reaches the cytosol (Fig. 3f and Supplementary Fig. 9e) and that replacing arginine by lysine in Penetratin significantly diminishes its internalization^68, 71^. According to our model, K9 should induce megapolarization and formation of water pores that should then allow their translocation into cells. However, it has been determined that, once embedded into membranes, lysine residues tend to lose protons^72, 73^. This will thus dissipate the strong membrane potential required for the formation of water pores and leave the lysine-containing CPPs stuck within the phospholipids of the membrane. In contrast, arginine residues are not deprotonated in membranes and water pores can therefore be maintained allowing the arginine-rich CPPs to be taken up by cells. This phenomenon however cannot be modeled by coarse-grained *in silico* simulations because the protonation state is fixed at the beginning of the simulation runs and is not allowed to evolve. Therefore, the uptake kinetics of lysine-rich peptide, such as MAP, appears artefactually similar as the uptake kinetics of arginine-rich peptides such as R9 (Supplementary Fig. 11b). CPPs can also enter cells through endocytosis, a mode of entry that is not inhibited by cell depolarization (Fig. 2c and Supplementary Fig. 7). The differences between CPPs in terms of how efficiently direct translocation is modulated by the V_m_ (Fig. 2c-d and Supplementary Fig. 10a) could be explained by their relative dependence on direct translocation or endocytosis to penetrate cells. The more positively charged a CPP is, the more it will enter cells through direct translocation and consequently the more sensitive it will be to cell depolarization (Fig. 2c). On the other hand, when endocytosis is the predominant type of entry, CPP cytosolic uptake will be less affected by both hyperpolarization and depolarization, which is what is observed for Transportan internalization in HeLa cells (Fig. 2c and Supplementary Fig. 10a).

We propose, based on the work described here, that hyperpolarization induced by drugs such as valinomycin represents a simple alternative or parallel approach to optimize CPP internalization. However hyperpolarizing drugs may be toxic when systemically applied. For example, valinomycin at the concentrations used to induce hyperpolarization (10 μM) would be lethal if systemically injected in mice (LD50 in the low micromolar^74^). On the other hand, local administration of valinomycin is far less toxic^75, 76^ as confirmed here in zebrafish and mice. Hyperpolarizing agents may therefore be preferentially used for local or topical applications, which is incidentally the case for the clinically approved CPPs^1, 2^.

Strategies to improve CPP delivery are becoming increasingly complex through the use of nanoparticles^77^, double-coated nanoparticles^78^, liposome-polycation-DNA complexes^79^, branched peptides^80^, etc. Our data provides a characterization at molecular level that can be taken advantage of to i) improve or optimize « old » CPPs, ii) design new CPPs, iii) help explain the behaviors of newly discovered CPPs^81–83^, iv) discriminate between target cells and cells that should be left unaffected based on V_m_ and v) distinguish between direct translocation and endosomal escape. The present work indicates that the impact on megapolarization should be evaluated when chemical modifications are performed on cationic CPPs to augment their delivery capacities.

## Author contributions

Conception and design of study: ET, GG, MH, NC, AD and CW

Acquisition of data: ET, GG, MH, NC, GD, YA, MS, SM, FO and LCD

Analysis and/or interpretation of data: ET, GG, MH, NC, MAD, MS, SM, GV, FO, LCD, JP, FA, AL, AD and CW

Funding acquisition: CW, NC and FA Resources: CW and FA

Drafting the manuscript: ET, GG and CW

Revising the manuscript and approval of the submitted version: all authors

## Acknowledgements

The lab of CW is supported by grants from the Swiss National Science Foundation (n° CRSII3_154420, IZCSZ0-174639 and 158116, awarded to CW and NC respectively). The zebrafish work performed in the lab of FA was supported the Swiss National Science Foundation (n° 320030_170062, to FA). We are thankful to Prof. Denise Nardelli Haefliger and her group, especially to Sonia Domingos Pereira and Laurent Derre for helpful discussions. We are thankful to the Cellular Imaging Facility, Mouse Pathology Facility and Genomic Technologies Facility at the University of Lausanne for the resources provided and their technical help. We would like to thank Giacomo Nanni, for his technical help in running the molecular simulations. We are also thankful to the Swiss National Supercomputing Centre (CSCS).

## Materials & Correspondence

Correspondence and requests for materials should be addressed to CW.

## Materials and Methods

### Chemicals

Puromycin 10 mg/ml (Thermo Fisher, ref. no. A11138-02) was aliquoted and stored at -20 °C. Blasticidin (Applichem, ref. no. A3784) was dissolved at 1 mg/mL in water and stored at -20 °C. XE-991 and TRAM-34 (Alomone labs, ref. no. X-100 and T-105 respectively) was dissolved in DMSO at 100 mM and stored at -20 °C. Cells were preincubated with 10 μM of these inhibitors for 30 minutes and then kept throughout the experiments. Pyrene butyrate (Sigma, ref. no. 257354) was dissolved in DMSO at 20 mM and stored at -20 °C. Live Hoechst 33342 (Sigma, ref. no. CDS023389) was aliquoted and stored at -20°C. Trypan Blue 0.4% (Life technologies, ref. no. 15250061) was stored at room temperature. AlexaFluor488-labeled human transferrin was dissolved in PBS 5 at mg/ml and stored at 4°C (Thermo Fisher, ref. no. 13342). TexasRed-labelled neutral 3’000 and 40’000 Da dextran was dissolved in PBS at 10 mg/ml and stored at -20°C (Thermo Fisher, ref. no. D3329 and D1829, respectively). TMR-labelled 10’000 neutral dextran was dissolved in PBS at 10 mg/ml and stored at -20°C (Thermo Fisher, ref. no. D1816).

### Antibodies

The rabbit polyclonal anti-V5 (Bethyl, ref. no. A190-A120), mouse monoclonal anti-FLAG antibody was from Sigma-Aldrich (ref. no. F1804), rabbit monoclonal anti-actin (Cell signaling, ref. no. 4970) and rat monoclonal anti-γ-tubulin (Santa Cruz, ref. no. sc-51715) antibodies were used for Western blotting.

### Cell lines

All cell lines were culture in 5 % CO2 at 37°C. Raji (kind gift from the laboratory of Aimable Nahimana, ATCC: CCL-86), SKW6.4 (kind gift from the laboratory of Pascal Schneider, ATCC: TIB-215) and HeLa (ATCC: CCL-2) cells were cultured in RPMI (Invitrogen, ref. no. 61870) supplemented with 10 % heat-inactivated fetal bovine serum (FBS; Invitrogen, ref. no. 10270-106). HEK293T (ATCC: CRL-3216) were cultured in DMEM supplemented with 10% FBS and were used only for lentiviral production. All cell lines were authenticated via Microsynth cell authentication service. Unless, otherwise indicated, experiments were performed in RPMI with 10% FBS.

### Zebrafish

Zebrafish (Danio rerio) from AB line were bred and maintained in our animal facility under standard conditions93, more specifically at 28.5°C and on a 14:10 hours light:dark cycle at the Zebrafish facility of the School of Biology and Medicine (cantonal veterinary approval VD-H21). Zebrafish of 20 hours post fertilization were collected and treated with 0.2 mM phenylthiourea (PTU, Sigma, St. Louis, MO) to suppress pigmentation. Embryos were raised at 28.5°C in Eggwater (0.3 g sea salt/L reverse osmosis water) up to 4 days post fertilization.

### Mice

C57BL/6NCrl were acquired from Charles River laboratories, which were then housed and bred in our animal facility. All experiments were performed according to the principles of laboratory animal care and Swiss legislation under ethical approval (Swiss Animal Protection Ordinance; permit number VD3374.a).

### Primary cortical neuronal culture

Sprague-Dawley rat pups (from Janvier, France) were euthanized in accordance with the Swiss Laws for the protection of animals, and the procedures were approved by the Vaud Cantonal Veterinary Office. Primary neuronal cultures from cortices of 2-day-old rats were prepared and maintained at 37 °C with a 5% CO2-containing atmosphere in neurobasal medium (Life Technologies, 21103-049) supplemented with 2% B27 (Invitrogen, 17504044), 0.5 mM glutamine (Sigma, G7513) and 100 μg/ml penicillin-streptomycin (Invitrogen, 15140122) as described previously^84^. Neurons were plated at a density of ∼3 × 10^5^ cells on 12-mm glass coverslips coated with 0.01% poly-L-lysine (Sigma, P4832). Half of the medium was changed every 3-4 days and experiments were performed at 12–13 days in vitro.

### Confocal microscopy

Confocal microscopy experiments were done on live 300’000 cells. Cells were seeded for 16 hours onto glass bottom culture dishes (MatTek, corporation ref. no. P35G-1.5-14-C) in 2 mL RPMI, 10%FBS and treated as described in the Figures in 1 mL media, 10%FBS. For nuclear staining, 10 μg/ml live Hoechst 33342 (Molecular probes, ref. no. H21492) was added in the culture medium 5 minutes before washing cells twice with PBS. After washing, cells were examined with a plan Apochromat 63x oil immersion objective mounted on a Zeiss LSM 780 laser scanning fluorescence confocal microscope equipped with gallium arsenide phosphide detectors and three lasers (a 405 nm diode laser, a 458-476-488-514 nm argon laser, and a 561 nm diode-pumped solid-state laser). Time-lapse experiments were done using an incubation chamber set at 37°C, 5% CO2 and visualized with a Zeiss LSM710 Quasar laser scanning fluorescence confocal microscope equipped with either Neofluar 63x, 1.2 numerical aperture (NA) or plan Neofluar 100x, 1.3 NA plan oil immersion objective (and the same lasers as above). Visual segregation of cells based on type of CPP entry, associated either with vesicular or diffuse cytosolic staining, was performed as shown in Supplementary Fig. 1c. Cell images were acquired at a focal plane near the middle of the cell making sure that nuclei were visible. Note: image acquisition was performed using the same settings for the data presented in the same panel or in the related supplementary panels, unless otherwise indicated.

### Flow cytometry

Flow cytometry experiments were performed using a Beckman Coulter FC500 instrument. Cells were centrifuged and resuspended in PBS prior to flow cytometry. Data analysis was done with Kaluza Version 1.3 software (Beckman Coulter).

### Cell death and CPP internalization measurements

With the exception of neurons, cell death was quantitated with 8 μg/ml propidium iodide (Sigma, ref. no. 81845). Unless otherwise indicated, cell death was assessed after 16 hours of continuous incubation in Raji and SKW6.4 cells and 24 hours in HeLa cells. Prior to treatment, 300’000 cells were seeded in 6-well plates for 16 hours in 2 mL media, 10% FBS. Treatment was done in 1 mL media with 10%FBS. Cell death and peptide internalization were analyzed by flow cytometry. Internalization measurements were done after one hour of incubation. Peptide internalization in primary cortical neurons was assessed by confocal microscopy with LSM780. Cell-associated fluorescence was quantitated with ImageJ.

### Lentivirus production

Recombinant lentiviruses were produced as described^92, 93^ with the following modification: the envelope plasmid pMD.G and the packaging vector pCMVΔR8.91 were replaced by pMD2.G and psPAX2, respectively.

### In vitro membrane potential measurements

Two methods were used to assess cellular membrane potential in vitro. With the first method, the membrane potential was determined by incubating 300’000 cells for 40 minutes with 100 nM of the fluorescent probe DiBAC4(3) (Thermofisher, ref. no. B438) in 6-well plates in 1 mL media, 10% FBS. and the median fluorescence intensity was then assessed by flow cytometry. Calculation of the actual membrane potential in mV based on the DiBAC4(3) signals was performed as described earlier^85, 86^. The second method relied on electrophysiology recordings. To perform these, the bath solution composition was (in mM): 103.9 NaCl, 23.9 NaHCO3, 2 CaCl2, 1.2 MgCl2, 5.2 KCl, 1.2 NaH2PO4, 2 glucose and 1.7 ascorbic acid. The pipet solution was composed of (in mM): 140 KMeSO4, 10 HEPES, 10 KCl, 0.1 EGTA, 10 phosphocreatine and 4 MgATP. The patch pipets had a resistance of 2.4-3.6 MΩ. Perforated patch recordings were performed as previously described^87^. Briefly, freshly prepared gramicidin D (Sigma, ref. no. G5002), at 2.8 μM final concentration, was added to prefiltered patch pipet solution and then sonicated for three consecutive times during 10 seconds. Cell-attached configuration was achieved by applying negative pressure on patch pipet until seal resistance of over one Giga Ohm was reached. After gaining cell access through gramicidin created pores, membrane potential measurements were done in current clamp at 0 pA for at least three minutes. Since primary rat neurons are killed following full depolarization induced by gramicidin, the standard curve from membrane potential calculations^105, 106^ were performed using gramicidin-treated Raji cells incubated with increasing concentrations of DiBac4(3) for 40 minutes. Images of the cells were then taken using a LSM780 confocal microscope and the cell-associated fluorescence quantitated with ImageJ.

### Relative membrane potential assessment in vivo

Zebrafish embryos in Egg water (see Zebrafish section) were incubated for 40 minutes in the presence or in the absence of various concentrations of valinomycin together with 950 nM DiBac4(3). The embryos were then fixed and visualized under a confocal LSM710 microscope^88^. DiBac4(3)-associated fluorescence of a region of interest of about 0.0125 mm2 in the tail region was quantitated with ImageJ. The values were normalized to the control condition (i.e. in the absence of valinomycin). Mice were intradermally injected with 10 μl of a 950 nM DiBac4(3) PBS solution containing or not 10 μM valinomycin and sacrificed one hour later. The skin was excised, fixed in 4% formalin, paraffin-embedded and used to prepare serial histological slices. Pictures of the slices were taken with a CYTATION3 apparatus. The DiBac4(3)-associated fluorescence in the whole slice was quantitated with ImageJ. The slice in the series of slices prepared from a given skin sample displaying the highest fluorescence signal was considered as the one nearest to the injection site. The signals from such slices are those reported in the Figures.

### Experimental modulation of the plasma membrane potential

#### Flow cytometry assessment of CPP internalization

Raji, SKW6.4 or HeLa cells: three hundred thousand cells were plated on non-coated plates to avoid cell adherence in 500 μl RPMI, 10% FBS. Cellular depolarization was induced by preincubating the cells at 37°C with 2 μg/ml gramicidin for 5 minutes and/or by placing them in potassium-rich buffer^41^ for 30 minutes (40 mM KCl, 100 mM potassium glutamate, 1 mM MgCl2, 1 mM CaCl2, 5 mM glucose, 20 mM HEPES, pH7.4). Cells were then treated with the selected CPPs at the indicated concentrations for one hour when peptide internalization was recorded or with 100 nM DiBac4(3) for 40 minutes when membrane potential needed to be measured. Hyperpolarization in Raji cells in the presence of TAT-RasGAP317-326 was performed by treating the cells with 10 μM valinomycin for 20 minutes in RPMI without serum. Cells were then treated with 5 µM TAT-RasGAP317-326 for one hour or 100 nM DiBac4(3) for 40 minutes. In the case of SKW6.4 and HeLa cells, hyperpolarization was induced by infection with a viral construct expressing KCNJ2 (see “virus production” section). Cells were then treated with 40 µM of indicated CPP for one hour or 100 nM DiBac4(3) for 40 minutes.

#### CPP cytosolic internalization quantitation based on confocal microscopy

Three hundred thousand wild-type HeLa cells were plated overnight on glass-bottom dishes in 2 mL RPMI with 10% FBS. The next day, serum was removed and cells were preincubated at 37°C with 2 μg/ml gramicidin for 5 minutes, 10 μM valinomycin for 20 minutes or were left untreated in 1 mL media with 10%FBS. The indicated CPPs were then added and cells were incubated for one hour at 37°C. Cells were then washed and visualized in RPMI without serum under a confocal microscope. CPP cytosolic internalization was quantitated within a cytosolic region devoid of endosomes using ImageJ. The number of CPP-positive vesicles was visually determined per cell in a given focal plane.

Neurons (12 days post-isolation) were preincubated 30 minutes with 5 mM TEA (tetraethylammonium, Sigma Aldrich, ref. no.T2265; gramicidin is toxic in these neurons; see section “Membrane potential measurements in vitro”) to induce depolarization or 10 μM valinomycin to induce hyperpolarization in bicarbonate-buffered saline solution (116 mM NaCl, 5.4 mM KCl, 0.8 mM MgSO4, 1 mM NaH2PO4, 26.2 mM NaHCO3, 0.01 mM glycine, 1.8 mM CaCl2, 4.5 mg/mL glucose) in a 37°C, 5% CO2 incubator. The cells were then incubated one hour with 2 μM of FITC-labeled TAT-RasGAP_317-326_. The cells were finally washed thrice with PBS and images were acquired using a LSM780 confocal microscope. Cell-associated peptide fluorescence was quantitated using ImageJ.

### Setting membrane potential by changing potassium concentrations in the media

RPMI-like media made without potassium chloride and without sodium chloride was from Biowest (Table S1). Varying concentrations of potassium chloride were added to this medium containing 10% FBS. Sodium chloride was also added so that the sum of potassium and sodium chloride equaled 119 mM (considering the concentrations of sodium and potassium in FBS). Three hundred thousand cells were preincubated in 1 mL media, 10% FBS containing different concentrations of potassium for 20 minutes, then different CPPs at a 40 μM concentration were added and cells were incubated for one hour at 37°C in 5% CO2. Cells were washed once in PBS and CPP internalization was measured by flow cytometry. The corresponding membrane potential was measured with DiBac4(3).

### In silico CPP translocation free energy though MARTINI coarse-grained simulations

An asymmetric multi-component membrane was constructed and solvated using CHARMM-GUI^97, 98^. Each layer contained 100 lipids (Table S2), in a previously described composition^99^. The membrane was solvated with 2700 water molecules, obtaining a molecular system of 10200 particles. The MARTINI force field^60^ was used to define phospholipids’ topology through a coarse-grained (CG) approach. The polarizable water model has been used to assess the water topology^89^. The R9 peptide model has been obtained by PEPFOLD-3 server^90^, as done in previous studies in the field^69, 91^. For each molecular system, R9 peptide was positioned 3 nm far from the membrane outer leaflet, in the water environment corresponding to the extracellular space. The elastic network ELNEDYN^92^ has been applied to reproduce the structural and dynamic properties of the CPPs. The molecular system has been minimized by a steepest descent protocol, and then equilibrated through five MD simulations of 1ns each under the NPT ensemble. Position restraints were applied during the first three molecular dynamics (MD) simulations and gradually removed, from 200 kJ/mol*nm2 to 10 kJ/mol*nm2. Velocity rescaling^102^ temperature coupling algorithm and time constant of 1.0 ps were applied to keep the temperature at 310.00 K. Berendsen^93^ semi-isotropic pressure coupling algorithm with reference pressure equal to 1 bar and time constant 5.0 ps was employed. Electrostatic interactions were calculated by applying the particle-mesh Ewald (PME)^94^ method and van der Waals interactions were defined within a cut-off of 1.2 nm. Periodic boundary conditions were applied in all directions. Trajectories were collected every 10 ps and the Visual Molecular Dynamics (VMD)^95^ package was employed to visually inspect the simulated systems. Three different transmembrane potential values have been considered: 0 mV, 80 mV, and 150 mV. In the MD simulations, an external electric field Eext was applied parallel to the membrane normal z, i.e., perpendicular to the bilayer surface. This was achieved by including additional forces Fi =q*Eext acting on all charged particles i. In order to determine the effective electric field in simulations, we applied a computational procedure reported in literature^96^. A well-tempered metadynamics protocol^97^ was applied to estimate the free energy landscape of CPP translocation. Two collective variables have been considered: the lipid/water density index, and the CPP-membrane distance. Further details about the lipid/water density index definition are reported in Supplementary Fig. 15. The metadynamics protocol has been carefully validated, as reported in Supplementary Table S5. Gaussian deposition rate of 2.4 kJ/mol every 5 ps was initially applied and gradually decreased on the basis of an adaptive scheme. Gaussian widths of 0.5, and 0.2 nm were applied following a well-established scheme^98–101^. In particular, the Gaussian width value was of the same order of magnitude as the standard deviation of the distance CV, calculated during unbiased simulations. The well-tempered metadynamics simulations were computed using GROMACS 2019.4 package^102^ and the PLUMED 2.5 open-source plug-in^103^. The reconstruction of the free-energy surface was performed by the reweighting algorithm procedure^104^, allowing the estimation of the free energy landscape. Each system was simulated (with a 20 fs time step) until convergence was reached. Further details about the convergence of each metadynamics simulation are reported in Supplementary Fig. 17-19. The electrostatic potential maps were computed by the APBS package^105^ on the molecular system composed of R9 peptide in contact with the cell membrane, without any applied external electrostatic field. In detail, the non-linear Poisson-Boltzmann equation was applied using single Debye-Huckel sphere boundary conditions on a 97×97×127 grid with a spacing of 1Å centered at the COM of the molecular system. The relative dielectric constants of the solute and the solvent were set to 2.5 and 78.4, respectively. The ionic strength was set to 150 mM and the temperature was fixed at 310K^101, 105^. The average and standard deviation values of the local transmembrane potential have been computed considering ten different trajectory snapshots taken from the molecular trajectory.

### In silico cell membrane hyperpolarization modeling through ion-imbalance in Martini coarse-grained simulations

The translocation mechanism of each CPP has been studied by ion-imbalance in a double bilayer system^24, 62–64^. The same asymmetric membrane considered to perform the single-bilayer simulations was used to build up the double-bilayer system. The double-membrane system was solvated with 4300 water molecules, obtaining a molecular system of 20000 particles. The MARTINI force field was used to define phospholipids’ topology through a coarse-grained (CG) approach. The polarizable water model has been used to model the water topology^89^. The elastic network ELNEDYN^106^ has been applied to reproduce the structural and dynamic properties of the CPPs.

For each molecular system, one CPP was positioned in the middle of the double bilayer system, 2nm far from the membrane outer leaflets, in the water environment corresponding to the extracellular space. Then, the system was equilibrated through four MD simulations of 100ps, 200ps, 500 ps, and 100 ns under the NPT ensemble. Position restraints were applied during the first three MD simulations ang gradually removed, from 200 kJ/mol*nm2 to 10 kJ/mol*nm2. Velocity rescaling^107^ temperature coupling algorithm and time constant of 1.0 ps were applied to keep the temperature at 310.00 K. Berendsen^93^ semi-isotropic pressure coupling algorithm with reference pressure equal to 1 bar and time constant 5.0 ps was employed. Then, all systems were simulated for the production run in the NPT ensemble with the time step of 20 fs with Parrinello-Rahman pressure coupling^108^.

Membrane hyperpolarization was achieved through a net charge difference of 30 positive ions between intracellular and extracellular space, considering all charged ions of the system and fulfilling the full system electroneutrality. Ten different replicas of each molecular simulation have been performed until the water pore formation and closure events have been observed. The visual inspection of the simulated molecular systems is reported in Supplementary Figure 11a. To analyze whether the CPPs were able to cross the membrane and reach the intracellular compartment, their trajectories were studied in the last five nanoseconds of each simulation replica. Considering the CPP position with respect the membrane bilayers and the CPP’s solvent accessible surface area (SASA), three different compartments were defined: intracellular space, lipid bilayer (cell membrane) and extracellular space. The radius of the water pores within the membrane was calculated as previously done in literature^66, 121^. We assumed that the central part of the cylindrical water pore contains N water molecules at the same density as outside of the water flux.

### In vitro assessment of water pores

Three hundred thousand wild-type HeLa cells were incubated with 32 ug/ml PI (0.8-1.5 nm diameter 75) or 200 µg/ml dextran of different size in the presence or in the absence of indicated CPP in normal, depolarizing (2 µg/ml gramicidin) or hyperpolarizing (10 µM valinomycin) conditions in 1 mL media, 10% FBS. Time-lapse images were acquired by confocal microscopy every 10 seconds. The percentage of cells where direct CPP translocation has occurred, as well as the percentage of cells positively stained for PI, were manually quantitated using ImageJ based on snap shot images taken after 30 minutes of incubation, as shown in Supplementary Fig. 12b. Quantitation of cell percentage was not selective in terms of fluorescence intensity. Cytosolic PI fluorescence was assessed with ImageJ, by selecting a region within the cell cytoplasm devoid of endosomes. Saponin 0.1% (Sigma, ref. no. 4706, diluted in PBS weight:volume) was used as a permeabilizing agent (30 minutes incubation at 37°C in a 5% CO2 incubator) that leads to cell death to differentiate signal intensity between live cells with water pores and dead cells. As, the signal of PI internalization in saponin treated cells was saturated, lower laser settings were used to look at dead cells than at cells with water pores. Three fitting models were obtained:

- exponential decline: *y* = 2570 · *e*^(−*x*/54)^

- exponential: *y* = 2570 · *e*^−0.02^ ^*x*^

- modified power: *y* = 2570 · 0.98^*x*^

These equations fitted equally well the PI uptake/Vm curve in Fig. 3h. For the calculations used in Fig. 3i, the exponential decline equation was used.

### Zebrafish viability

FITC-TAT-RasGAP_317-326_(W317A) internalization in zebrafish was assessed either by adding the peptide directly in Egg water or by injection. Experiments in which the peptide was added in the water were performed on fish between four and 24 hours post fertilization. Viability assays were done on embryos of four, six and 24 hours post fertilization to determine a maximal nonlethal dose of the peptide that can be used. Different concentrations of the peptide were added to 500 µl water per well in 24-well plate, with between eight and eleven embryo per well. Fish viability was visually assessed at 20 hours post incubation with the peptide. Hyperpolarization associated viability was visually assessed at different time points in presence or in the absence of the peptide and in presence or in the absence of different concentration of valinomycin. Zebrafish were visualized with binocular microscope and CYTATION3 apparatus. Survival was visually assessed under a binocular microscope by taking into consideration the embryo transparency (as dead embryos appear opaque), general development characteristics and motility.

### CPP internalization in vivo

To assess peptide internalization in zebrafish, two methods were used: 1) addition of the peptide directly in 500 µl of Egg water in 24-well plates containing between eight and twelve embryos per well or 2) intramuscular injections. In the case of the first method, after the indicated treatments, zebrafish were washed, fixed in 4% PFA/PBS for one hour at room temperature. Whole embryos were mounted on slides with Fluoromount-G (cBioscience, ref. no. 00-4958-02). Zebrafish were then visualized under a LSM710 confocal microscope. Experiments where the peptide was added directly to the water were performed on zebrafish at 18 hours post fertilization to limit cuticle development that would hinder peptide access to the cells. In the case of the second method, 8 nl injections (containing the various combinations of peptide and valinomycin and 0.05 % (vol:vol) phenol red as an injection site labelling agent) were done on 48 hours post fertilization embryos into the tail muscle around the extremity of yolk extension, after chorion removal and anesthesia with 0.02% (w:vol) tricaine ^122^ buffered in sodium bicarbonate to pH7.3. At this age, zebrafish already have well developed tissues that can be easily visually distinguished. Injections were done with an Eppendorf Microinjections FemtoJet 4i apparatus. After the indicated treatments, embryos were fixed in 4% PFA/PBS and visualized under a confocal microscope. Some embryos were kept alive for viability evaluation post injection until the age of 4 days.

Experiments with mice were performed in 10-14 weeks old C57BL/6NCrl mice anaesthetized with ketasol/xylasol (9.09 mg/ml ketasol and 1.82 mg/ml xylasol in water; injection: 10 μl per g of body weight). The back of the mice was shaved and intradermic injections were performed (a total of 10 μl was injected). Mice were kept under anesthesia for one hour and Artificial tears (Lacryvisc) were used to avoid eye dryness. Mice were then sacrificed by CO2 inhalation, skin was cut at injection sites, fixed in 4% formalin and paraffin embedded for histology analysis. For each sample, 10 to 15 slides were prepared and peptide internalization was visualized with a CYTATION3 apparatus. Fluorescence intensity was quantitated with ImageJ. The slices displaying the highest fluorescence signal were considered as those nearest to the injection site and the fluorescent values from these slides were used in Fig. 4b.

### Temperature dependent internalization

Two hundred thousand cells were incubated with 40 µM (Raji and SKW6.4) or 80 μM (HeLa) of FITC-labelled TAT-RasGAP_317-326_ for one hour in Eppendorf tubes in RPMI media supplemented with 10% serum and 10 mM Hepes on a thermoblock at different temperatures. Peptide internalization was measured by flow cytometry after PBS wash and addition of 0.2% trypan blue (in PBS) to quench extracellular fluorescence.

### Pyrene butyrate treatment

Three hundred thousand cells were seeded in 6-well plates in 2 mL RPMI, 10% FBS for 16 hours. Cells were then washed twice with serum-free medium and treated for 10 minutes with 50 µM pyrene butyrate or 0.25% (vol:vol) DMSO as vehicle control at 37°C in 5% CO2 in 1 mL media. Then cells were treated with TAT-RasGAP_317-326_ for the indicated period of time.

### Assessment of endosomal escape and direct translocation

Three hundred thousand cells were seeded onto glass-bottom dishes in 2 mL RPMI, 10% FBS for 16 hours. Quantitation of cytosolic fluorescence was performed within live HeLa cells pre-incubated with 80 µM TAT-RasGAP_317-326_ for 30 minutes at 37°C in 1 mL media, with 10% FBS and then incubated for the indicated periods of time in the presence (i.e. no wash after the pre-incubation) or in the absence (i.e. following three consecutive washes with RPMI supplemented with 10% FBS) of extracellular labelled peptide. Endosomal escape from lysosomes was induced in presence of 1 mM LLOME (L-Leucyl-L-Leucine methyl ester)^49^, added in the 1 mL media, 10% FBS 30 minutes after CPP was washed out and persisted throughout the experiment). Confocal images were taken every five minutes after the 30-minute pre-incubation. For each cell, the fluorescence intensity of one region of interest (ROI) devoid of labelled endosomes throughout the experiment was quantitated over time using ImageJ Time Series Analyzer V3. The surface of the ROI was identical for all cells. Only cells displaying labelled endosomes after the 30-minute pre-incubation were analyzed. Note that the washing steps, for reasons unclear at this time, induced a slightly higher initial ROI intensity signal.

### Transferrin internalization quantitation

Wild-type HeLa cells were plated in 12-well plates (200’000 cells per well) for 16 hours in 1 ml RPMI (Invitrogen, ref. no. 61870), supplemented with 10 % heat-inactivated fetal bovine serum (FBS; Invitrogen, ref. no. 10270-106). Cells were then incubated in presence of 20 µg/ml AlexaFluor-488 conjugated transferrin for 20 minutes at 37°C in 5% CO2. Cells were washed with PBS and pelleted after trypsinization. To quench membrane bound transferrin fluorescence, cells were resuspended in 0.2% trypan blue diluted in PBS. Transferrin internalization was quantified by flow cytometry using Beckman Coulter FC500 instrument. Data analysis was done with Kaluza Version 1.3 software (Beckman Coulter).

### TAT-PNA-induced luciferase activity

The LeGOiG2-LUC705 lentiviral construct encodes a luciferase gene interrupted by a mutated human beta globin intron 2. This mutation creates a new aberrant splicing site at position 705 that when used produced an mRNA that encodes a truncated non-functional luciferase^50^. In the presence of the TAT-peptide nucleic acid (TAT-PNA) CPP described below, the aberrant splice site is masked allowing the production of a functional luciferase enzyme. Lentiviruses produced using this construct were employed to infect cells. The doses used resulted in >90% cells infected (based on GFP expression from the lentiviral vector). The infected cells (200’000 cells in 12-well plates containing 1 ml of RPMI, 10% FBS) were treated or not with 5 μM TAT-PNA (GRKKRRQRRR-CCTCCTACCTCAGTTACA). TAT-PNA is made of TAT_48-57_ and an an oligonucleotide complimentary to a sequence containing the aberrant splice site. After 16 hours incubation, cells were washed twice in HKR buffer (119 mM NaCl, 2.5 mM KCl, 1 mM NaH2PO4, 2.5 mM CaCl2, 1.3 mM MgCl2, 20 mM HEPES, 11 mM dextrose, pH 7.4) and lysed in 40 μl HKR containing 0.1 % Triton X-100 for 15 minutes at room temperature. Luciferase activity was measured with a GLOMAXTM 96 Microplate Luminometer (Promega) using a Dual-Luciferase Reporter Assay (Promega) and normalized to the protein content. Results are displayed as the ratio between the protein-normalized luciferase signal obtained in TAT-PNA-treated cells and the signal obtained in control untreated cells.

### TAT-Cre recombinase production, purification and recombination assay

Raji cells were infected with a lentivirus encoding a Cre-reporter gene construct^53^. TAT-Cre recombinase was produced as described^54^. Briefly, E. coli BL21 transformed with the pTAT-Cre plasmid (#917, Addgene plasmid #35619) were grown for 16 hours in LB containing 100 µg/mL kanamycin. Protein production was induced at OD600 of 0.6 with 500 µM IPTG (isopropyl β-D-1-thiogalactopyranoside) for three hours. Bacteria were collected by centrifugation at 5000 x g and kept at -20°C. Purification was performed on Äkta prime (GE, Healthcare, USA) equipped with a 1 ml HisTrap FF column equilibrated with binding buffer (20 mM sodium phosphate, 500 mM NaCl, 5 mM imidazole pH 7.4). The day of the purification, bacterial pellet was resuspended in lysis buffer (binding buffer with protease inhibitors (Roche, ref. no. 4693132001; one tablet per 50 ml), 0.025 mg/ml DNase I (Roche, ref. no. 04716728001), and 2 mg/ml lysozyme (Roche, ref. no. 10 837 059 001) and sonicated six times for 30 seconds. After 20 minutes centrifugation at 5’000 x g, the supernatant was filtered through Steriflip 0.45µm and loaded on the column. Elution buffer (20 mM sodium phosphate, 500 mM NaCl, 500 mM imidazole pH 7.4) was used to detach His-tagged proteins from the column. Imidazole was removed from collected fractions by overnight dialysis using 10K MWCO cassette (Thermo Scientific, ref. no. 66807) in PBS. Raji cells encoding the Cre-reporter were treated for 48 hours with 20 µM TAT-Cre-recombinase. Fluorescence was imaged using a Nikon Eclipse TS100 microscope.

### Extracellular pH manipulation

Three hundred thousand HeLa cells were incubated in 1 mL RPMI media, supplemented with 10% FBS in 6-well plates, set at the indicated pH for 20 minutes at 37°C in 5% CO2, with or without 2 µg/ml gramicidin to induce depolarization. Changes to the extracellular pH were done by adding NaOH or HCl to the RMPI media. The pH was measured with a pH meter (Metrohm, 744 pH Meter) at the start and at the end of the experiment. TMR-TAT-RasGAP_317-326_ 40 µM was then added to cells for one hour. Cytosolic peptide internalization as well as TMR-TAT-RasGAP_317-326_-positive vesicles were quantitated based on confocal images. Cytosolic CPP fluorescence was assessed with ImageJ, by selecting a region within the cell cytoplasm devoid of endosomes. These experiments were performed in the presence of FBS because in its absence, cells are too sensitive to acidification or alkalization. In addition, TMR-labelled peptide was used to avoid FITC-quenching at acidic pH.

### Assessment of CPP binding to plasma membranes

Three hundred thousand cells were incubated for 60 seconds in 1 mL RPMI supplemented with 10% FBS and 10 mM HEPES in Eppendorf tubes at 37°C in the presence of increasing concentrations of FITC-TAT-RasGAP_317-326_. Half of the cells were then immediately placed on ice, pelleted at 4°C, and resuspended in one ml of ice-cold PBS and then split into two tubes, one of which receiving a final concentration of 0.2% (w:w) trypan blue to quench surface-associated FITC signals. The cells (still kept at 4°C) were then analyzed by flow cytometry. The surface associated peptide signal was calculated by subtracting total fluorescence measured in PBS and fluorescence measured after trypan blue quenching. The other half of the cells after the 60 second peptide incubation was incubated at 37°C for one hour at which time the cellular internalization of the labelled peptide was assessed by flow cytometry.

### Transient calcium phosphate transfection in HeLa cells

Calcium phosphate-based transfection of HeLa cells was performed as previously described^109^. Briefly, cells were plated overnight in DMEM (Invitrogen, ref. no. 61965) medium supplemented with 10 % heat-inactivated FBS (Invitrogen, ref. no. 10270-106), 2.5 ug of total plasmid DNA of interest was diluted in water, CaCl2 was added and the mixture was incubated in presence of HEPES 2x for 60 seconds before adding the total mixture drop by drop to the cells. Media was changed 10 hours after.

### Isothermal titration calorimetry (ITC)

ITC was performed using MicroCal ITC200 (Malvern Panalytical) at 37°C with 600 M FITC-R9 in the cell (total volume 300 µl) and consecutive injections (2.5 µl/injection, except for the first injection of 0.4 µl) of 6 mM PI from the syringe (total volume 40 µl) with 2 minutes delay between injections and 800 rotations/minute rotation speed. Differential power was set to 7, as we had no prior knowledge of the expected reaction thermodynamics. The results in Supplementary Fig. 12c are represented as a: thermogram (measurement of thermal power need to ensure that there is no temperature difference between reference and sample cells in the calorimeter as a function of time) and a binding isotherm (normalized heat per peak as a function of molar ratio).

### Colony formation assay

Three hundred thousand wild-type HeLa cells were plated overnight in RPMI with 10% FBS in 6-well plates. Cells were then treated for one hour in the presence of indicated concentrations of CPP, PI and membrane potential modulating agents (gramicidin or valinomycin) in 1 mL RPMI. As control, cells were either left untreated or incubated in presence of DMSO used as vehicle for gramicidin and valinomycin. Cells were then washed, trypsinized and plated on 10 cm dishes at a density of 300 cells per condition. Colonies were counted at day 14 after 100% ethanol fixation for 10 minutes and Giemsa staining. Washes were done with PBS.

### Genome-scale CRISPR/Cas9 Knockout screening

The human GeCKO v2 library (2 plasmid system) (Addgene plasmid #1000000049) was amplified by electroporation using a Bio-Rad Gene Pulser II electroporation apparatus (Bio-Rad #165-2105) and the Lucigen Endura bacteria (Lucigen ref. no. 60242). Cells were plated on LB Agar plates containing 100 µg/mL ampicillin. After 14 hours at 32°C, colonies were scrapped and plasmids recovered with the Plasmid Maxi kit (Qiagen, ref. no. 12162). To produced lentivirus library, 12 T-225 flasks were seeded with 12×106 HEK293T cells per flask in 40 ml DMEM, 10% FBS. The day after, 10 µg pMD2.G, 30 µg psPAX2 and 25 µg GeCKO plasmid library in 1.8 ml H2O were mixed with 0.2 ml 2.5 M CaCl2 (final calcium concentration: 250 mM). This solution was mixed (v/v) with 2x HEPES buffer (280 mM NaCl, 10 mM KCl, 1.5 mM Na_2_HPO_4_, 12 mM D-glucose, 50 mM HEPES), incubated for one minute at room temperature, added to the culture medium, and the cells placed back in a 37°C, 5% CO_2_ incubator for seven hours. The culture medium was then removed and replaced by DMEM supplemented with 10 % FBS containing 100 U/ml penicillin and 100 µg/ml streptomycin. Forty-eight hours later, the medium was collected and centrifuged 5 min at 2’000 g to pellet the cells. The remaining cell-free medium (12 x 40 ml) was then filtered through a 0.45 µm HV/PVDF (Millipore, ref. no. SE1M003M00) and concentrated ∼100 times by resuspending the viral pellet obtained by ultracentrifugation at 70’000 g for two hours at 4°C in ∼5 ml ice-cold PBS. The concentrated viruses were aliquoted in 500 µl samples and stored at -80°C.

To express the Cas9 endonuclease, cells (e.g. Raji or SKW6.4) were infected with Cas9 expressing viruses that were produced in HEK293T cells transfected with the lentiCas9-Blast (#849, Addgene plasmid #52962), pMD2.G, and psPAX2 plasmids as described in the main method under “Lentivirus production”. The infected cells were selected with 10 µg/mL blasticidin for a week. The multiplicity of infection (MOI) of the GeCKO virus library was determined as follow. Different volumes of the virus library were added to 3x10^6^ Cas9-expressing cells plated in 12-well plates. Twenty-four hours later, the cells were split in two wells of 12-well plates. One well per pair was treated with 10 µg/mL puromycin for 3 days (the other cultured in normal medium). Cell viability was determined by trypan blue exclusion and MOI was calculated as the number of cells in the well treated with puromycin divided by the number of cells in the control well. The virus volume yielding to a MOI ∼0.4 was chosen to perform large-scale infection of 12x107 cells that was carried out in 12-well plates with 3x106 cells per well. After 24 hours, the infected cells were collected and pooled in a T-225 flask and selected with 10 µg/mL puromycin for a week. Thirty million of these were frozen (control untreated cells) and 60 million others were treated with 40 µM TAT-RasGAP_317-326_ for 8 days (Raji) or for 17 days (SKW6.4) with a medium and peptide renewal every 2-3 days. Thirty million of the peptide-treated cells were then also frozen. Genomic DNA was extracted from the control and the peptide-treated frozen cells using the Blood & Cell Culture DNA Midi Kit according to manufacturer’s instructions (Qiagen, ref. no. 13343). A first PCR was performed to amplify the lentiCRISPR sgRNA region using the following primers:

F1: 5’-AATGGACTATCATATGCTTACCGTAACTTGAAAGTATTTCG-3’

R1: 5’-CTTTAGTTTGTATGTCTGTTGCTATTATGTCTACTATTCTTTCC-3

A second PCR (see Table S3 for the primers used) was performed on 5 µl of the first PCR reaction to attach Illumina adaptors with barcodes (nucleotides highlighted in green) and to increase library complexity (using the sequences highlighted in red) to prevent signal saturation when the sequencing is performed. The blue sequences are complementary to the extremities of the first PCR fragments.

Both PCRs were performed in 100 µl with the 2 µl of the Herculase II Fusion DNA Polymerase from Agilent (ref. no. 600675) according the manufacturer’s instructions. Amplicons were gel extracted, quantitated, mixed and sequenced with a MiSeq (Illumina). Raw FASTQ files were demultiplexed and processed to contain only unique sgRNA sequences. The number of reads of each sgRNA was normalized as described^125^. The MAGeCK algorithm^126^ was used to rank screening hits by the consistent enrichment among multiple sgRNAs targeting the same gene.

### CRISPR/Cas9-based genome editing

Single guide RNAs targeting the early exon of the protein of interest were chosen in the sgRNA library133 and are listed in table I below (Table S4). LentiCRISPR plasmids specific for a gene were created according to the provided instructions^127^. Briefly, oligos were designed as follow: Forward 5’-CACCGnnnnnnnnnnnnnnnnnnnn-3’; Reverse-3’-CnnnnnnnnnnnnnnnnnnnnCAA-5’, where nnnnnnnnnnnnnnnnnnnn in the forward oligo corresponds to the 20 bp sgRNA. Oligos were synthetized, then phosphorylated and annealed to form oligo complex. LentiCRISPR vector was BsmBI digested and dephosphorylated. Linearized vector was purified and gel extracted and ligated to oligo complex. The lentiCRISPR vector containing the sgRNA was then used for virus production. Recombinant lentiviruses were produced as described^93^ with the following modification: pMD.G and pCMVDR8.91 were replaced by pMD2.G and psPAX2 respectively. Cells were infected and selected with the appropriate dose of puromycin (2 µg/ml for HeLa). Clone isolation was performed by limiting dilution in 96 well-plate.

### TA cloning

TA cloning is a subcloning technique that allows integration of a PCR-amplified product of choice into a PCR2.1 vector based on complementarity of deoxyadenosine added onto the PCR fragment by Taq polymerase. This approach is useful to distinguish between several alleles and to determine whether the cells are heterozygous or homozygous at a given locus.

TA cloning kit (Life technologies, ref. no. K202020) was used according to manufacturer’s instructions to sequence DNA fragment containing the region targeted by a given sgRNA. Briefly, DNA was isolated and the fragment of interest was PCR-amplified using primers in Table S6, then ligated into PCR2.1 vector. *E. coli* competent cells were then transformed and at least 15 colonies were selected per condition for DNA isolation and sequencing.

### Plasmid constructs

The hKCNN4-V5.lti (#953) lentiviral plasmid encoding a V5-labeled version of the KCNN4 potassium channel was from DNASU (ref. n° HsCD00441560). The hKCNK5-FLAG.dn3 (#979) plasmid encoding the human KCNK5 potassium channel (NCBI reference sequence NM_003740.3), Flag-tagged at the C-terminus, was purchased from GenScript (ref. n° OHu13506). The Myc-mKCNJ2-T2A-IRES-tdTomato.lti (#978) lentiviral vector encoding the mouse Kir2.1 (KCNJ2) potassium channel and tdTomato (separated by an IRES) was generated by subcloning myc-mKCNJ2-T2A-Tomato.pCAG plasmid (#974, Addgene plasmid #60598) into a lentiviral backbone LeGo-iT2 (#809), a gift from Boris Fehse (Addgene plasmid #27343), through ligation of both plasmids after digestion with BamHI (NEB, reg. no. R313614). The pMD2.G plasmid (#554, Addgene plasmid #12259) encodes the envelope of lentivirus. The psPAX2 plasmid (#842, Addgene plasmid #12260) encodes the packaging system. Both pMD2.G and psPAX2 plasmids were used for lentiviral production. The Flag-hKCNQ5(G278S)-IRES-NeoR plasmid (#938) codes for the N-terminal Flag-tagged G278S human KCNQ5 inactive mutant and a neomycin resistant gene separated by an IRES sequence. It was generated by subcloning a BamHI/XmaI digested PCR fragment obtained by amplification of pShuttle-Flag-hKCNQ5(G278S)-IRES-hrGFP2 (#937, kind gift from Dr. Kenneth L. Byron) using forward primer #1397 (CAT CGG GAT CCG CTA TAC CGG CCA CCA TGG ATT ACA AGG A) and reverse primer #1398 (CAT CGC CCG GGG CTA TAC CGT ACC GTC GAC TGC AGA ATT C) into the lentiviral vector TRIP-PGK-IRES-Neo (#350) opened with the same enzyme. The Flag-hKCNQ5(SM,G278S)-IRES-Neo (#939) plasmid is identical to Flag-hKCNQ5(G278S)-IRES-NeoR except that the sequence targeted by the sgKCNQ5.1 sgRNA (Table S4) was mutated with the aim to decrease Cas9-mediated degradation. Silent mutations (SM), at the protein level, were introduced using the QuikChange II XL Site-Directed Mutagenesis Kit (ref. no. 200522) according to manufacturer’s instructions using forward primer #1460 (AAA TAA GAA CCA AAA ATC CTA TGT ACC ATG CCG TTA TCA GCT CCT TGC TGT GAG CAT AAA CCA CTG AAC CCA G) and reverse primer #1461 (CTG GGT TCA GTG GTT TAT GCT CAC AGC AAG GAG CTG ATA ACG GCA TGG TAC ATA GGA TTT TTG GTT CTT ATT T).

The Flag-hKCNQ5(SM)-IRES-NeoR (#940) lentiviral construct codes for a Flag-tagged wild-type version of human KCNQ5. It was made by reverting the G278S mutation in Flag-hKCNQ5(SM,G278S)-IRES-Neo (#939) using the QuikChange II XL Site-Directed Mutagenesis Kit with the #1462 forward primer (TTT TGT CTC CAT AGC CAA TAG TTG TCA ATG TAA TTG TGC CCC) and the #1463 reverse primer (GGG GCA CAA TTA CAT TGA CAA CTA TTG GCT ATG GAG ACA AAA). The pLUC705^50^ (#876, gift from Dr. Bing Yang) plasmid encodes a luciferase gene interrupted by a mutated human beta globin intron 2. This mutation creates a new aberrant splicing site at position 705 that when used produced an mRNA that encodes a truncated non-functional luciferase^50^. To introduce this construct into a lentiviral vector, the pLUC705 plasmid was digested with HindIII/XhoI, blunted with T4 DNA polymerase, and ligated into StuI-digested and dephosphorylated LeGO-iG2 (#807, Addgene plasmid #27341), resulting in plasmid pLUC705.LeGO-iG2 (#875). The pTAT-Cre (#917, Addgene plasmid #35619) bacterial plasmid encodes a histidine-tagged TAT-Cre recombinase. The Cre reporter lentiviral vector (#918, Addgene plasmid #62732) encodes a LOXP-RFP-STOP-LOXP-GFP gene construct. Cells expressing this plasmid appear red but once recombination has occurred when TAT-Cre is translocated into cells the RFP-STOP fragment will be excised, GFP but not RFP will now be produced, and cells will appear green.

### Peptides

TAT-RasGAP_317-326_ is a retro-inverso peptide (i.e. synthesized with D-amino-acids in the opposite direction compared to the natural sequence) labeled or not with FITC or TMR. The TAT moiety corresponds to amino-acids 48–57 of the HIV TAT protein (RRRQRRKKRG) and the RasGAP_317–326_ moiety corresponds to amino-acids from 317 to 326 of the human RasGAP protein (DTRLNTVWMW). These two moieties are separated by two glycine linker residues in the TAT-Ras-GAP317–326 peptide. FITC-bound peptides without cargo: TAT, MAP (KLALKLALKALKAALKLA), Penetratin (RQIKWFQNRRMKWKK), Transportan (GWTLNSAGYLLGKINLKALAALAKKIL), R9 (RRRRRRRRR), K9 (KKKKKKKKK) and (RE)9 (RERERERERERERERERE) were synthesized in D-amino-acid conformation. All peptides were synthesized in retro-inverso conformation (over the years different suppliers were used with routine checks for activity of TAT-RasGAP_317-326_ derived peptides, Biochemestry Department of University of Lausanne, SBS Genetech, China and Creative Peptides, USA) and resuspended to 1 mM in water.

### Statistical analysis

Statistical analysis was performed on non-normalized data, using GraphPad Prism 7. ANOVA multiple comparison analysis to wild-type condition was done using Dunnett’s correction (Fig. 2a, 2b and 2c (top panel) and PI internalization in Fig. 3f, as well as TAT-PNA internalization in Supplementary Fig. 8a. ANOVA multiple comparison analysis between several conditions was done using Tuckey’s correction (Supplementary Fig. 10a and 10e). Comparison between two conditions was done using two-tailed paired t-test for the CPP internalization experiments described in Fig. 2c (bottom panel), 4a-b and Supplementary Fig. 1d, 10c. All measurements were from biological replicates. Unless otherwise stated, the horizontal bars in the graph represent the median, the height of columns correspond to averages, and the dots in the Figures correspond to values derived from independent experiments.

### Data availability

DNA sequencing data from the CRISPR/Cas9-based screens are available through the following link: https://www.ncbi.nlm.nih.gov/sra/SRP161445. Note: this link is currently blocked for public access, but will be released upon publication.

---

**Supplementary Fig. 1.**
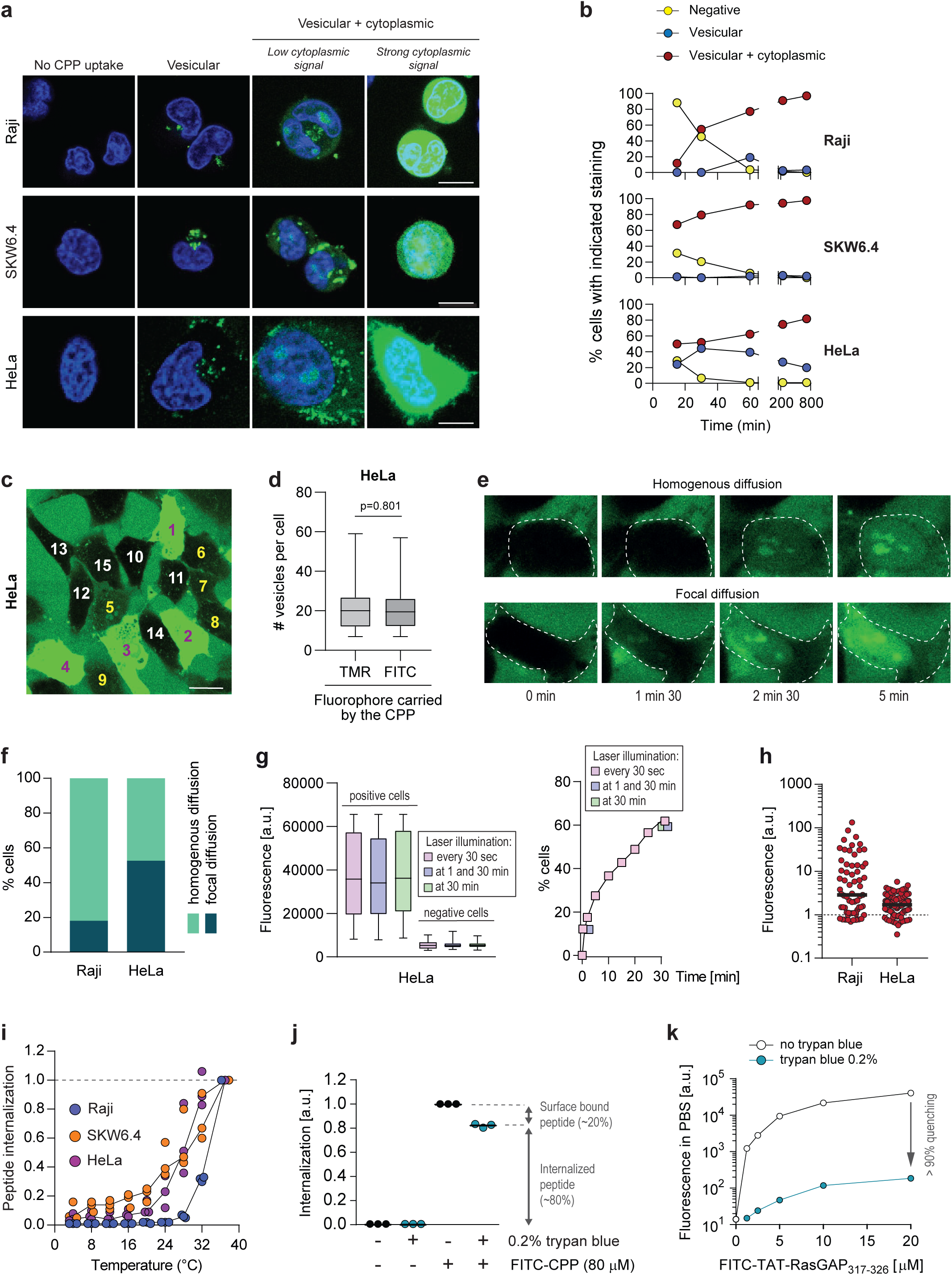

**Supplementary Fig. 2.**
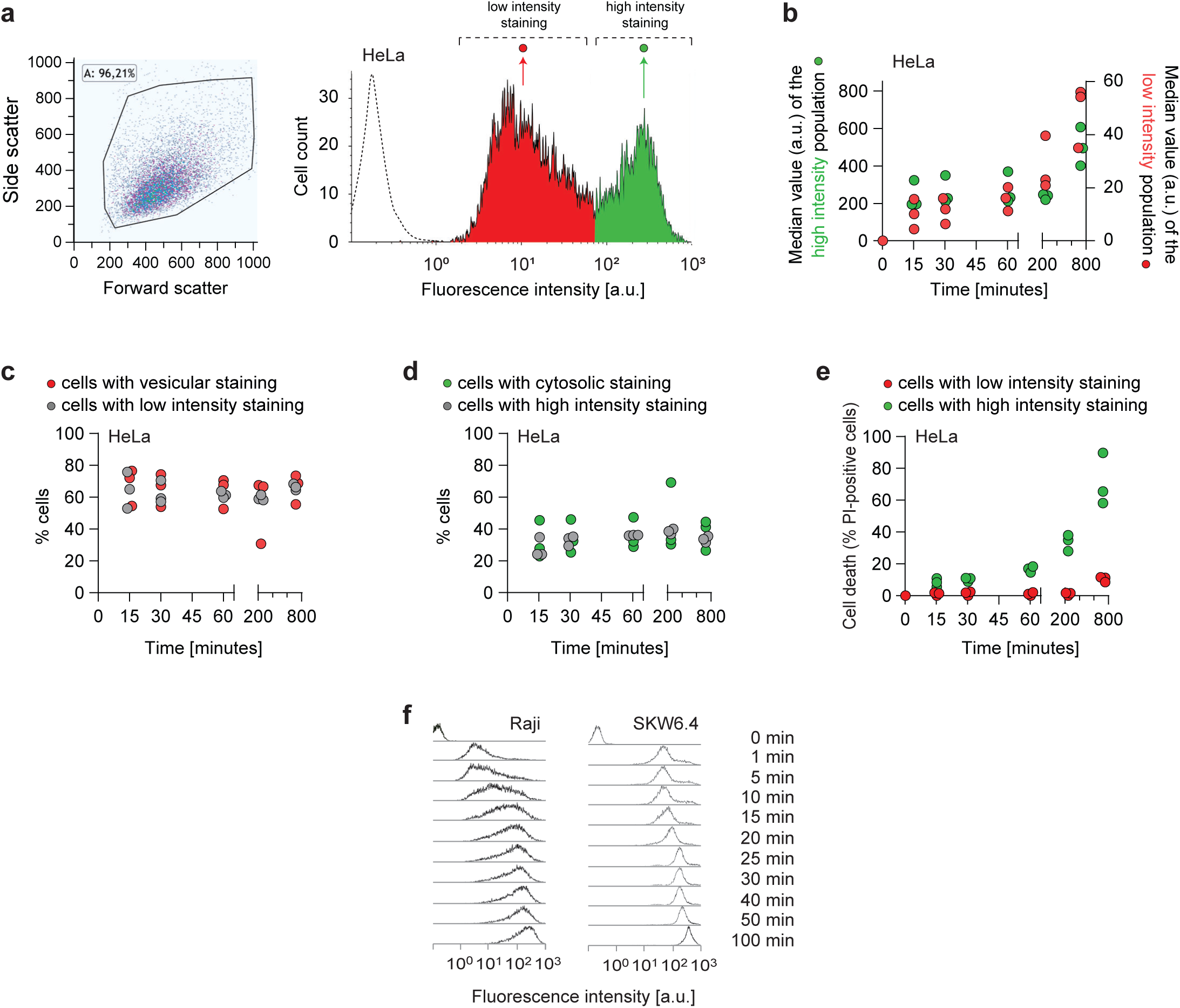

**Supplementary Fig. 3.**
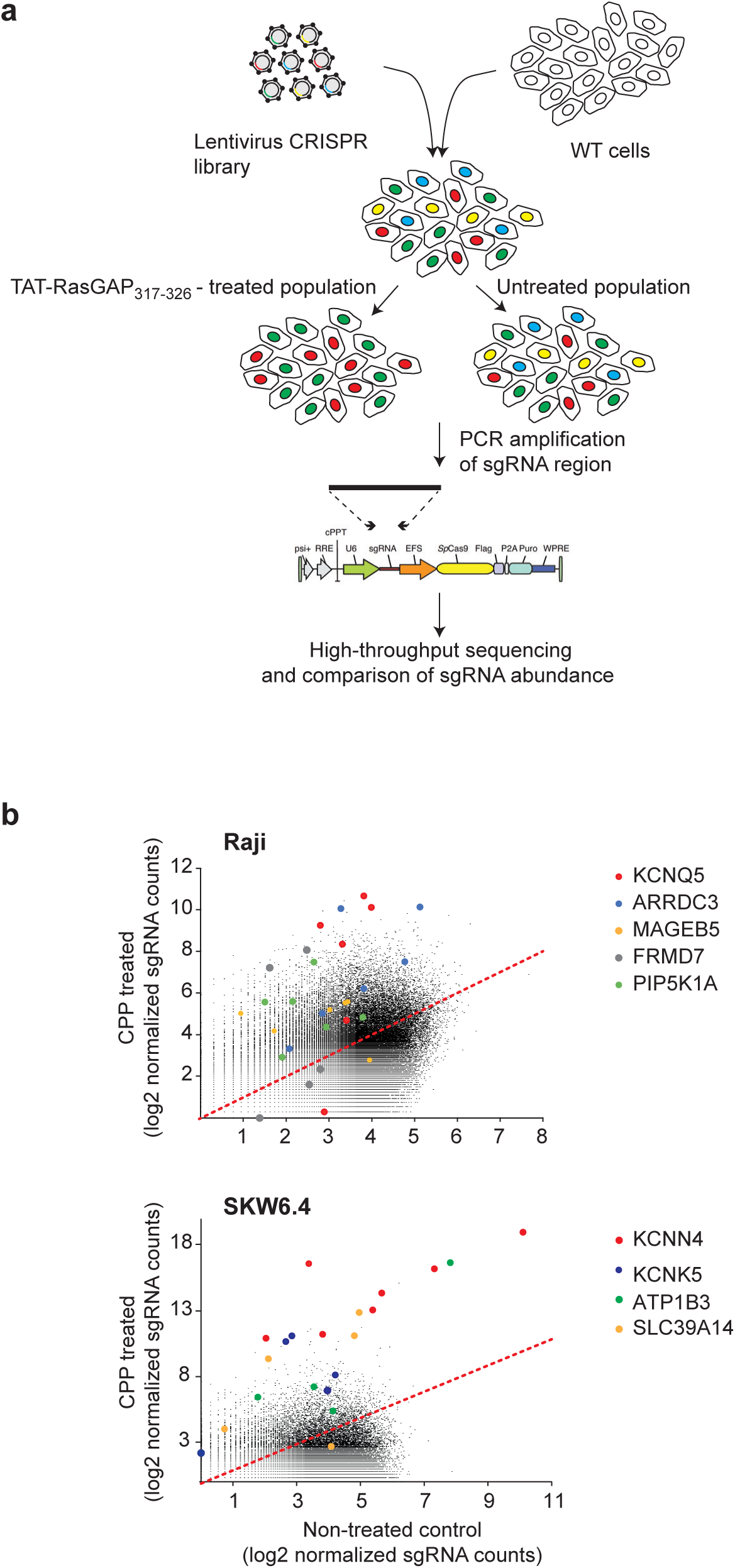

**Supplementary Fig. 4.**
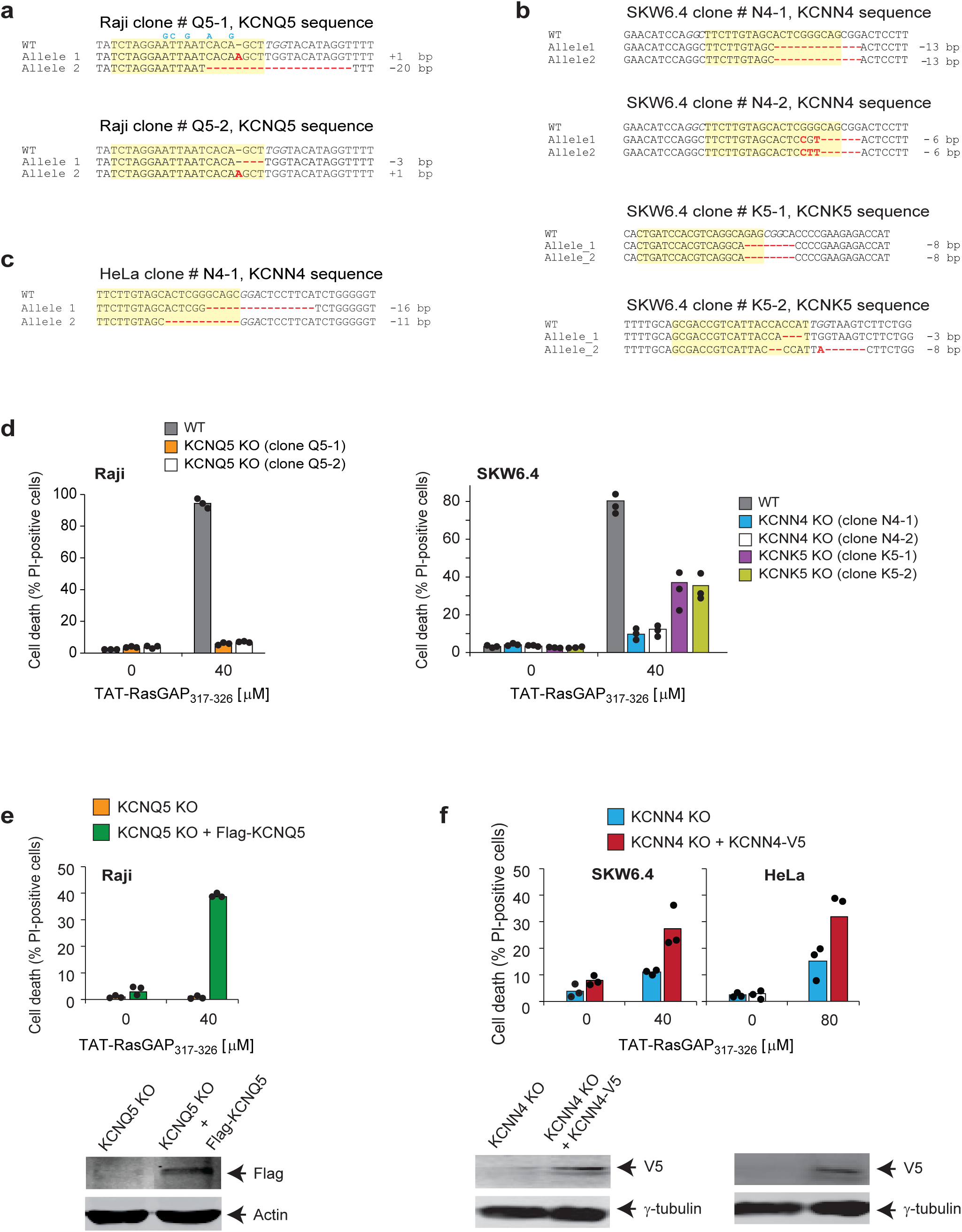

**Supplementary Fig. 5.**
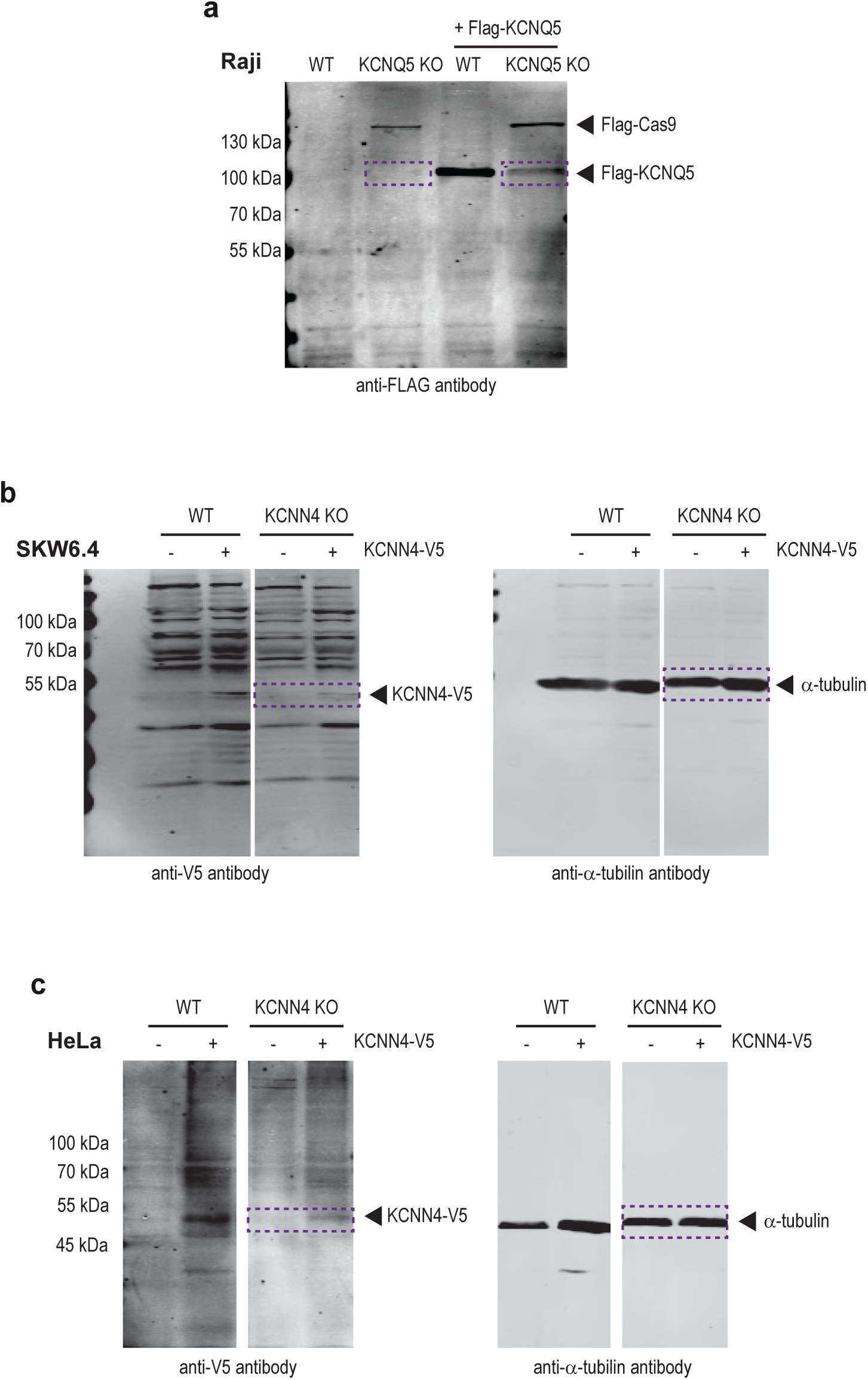

**Supplementary Fig. 6.**
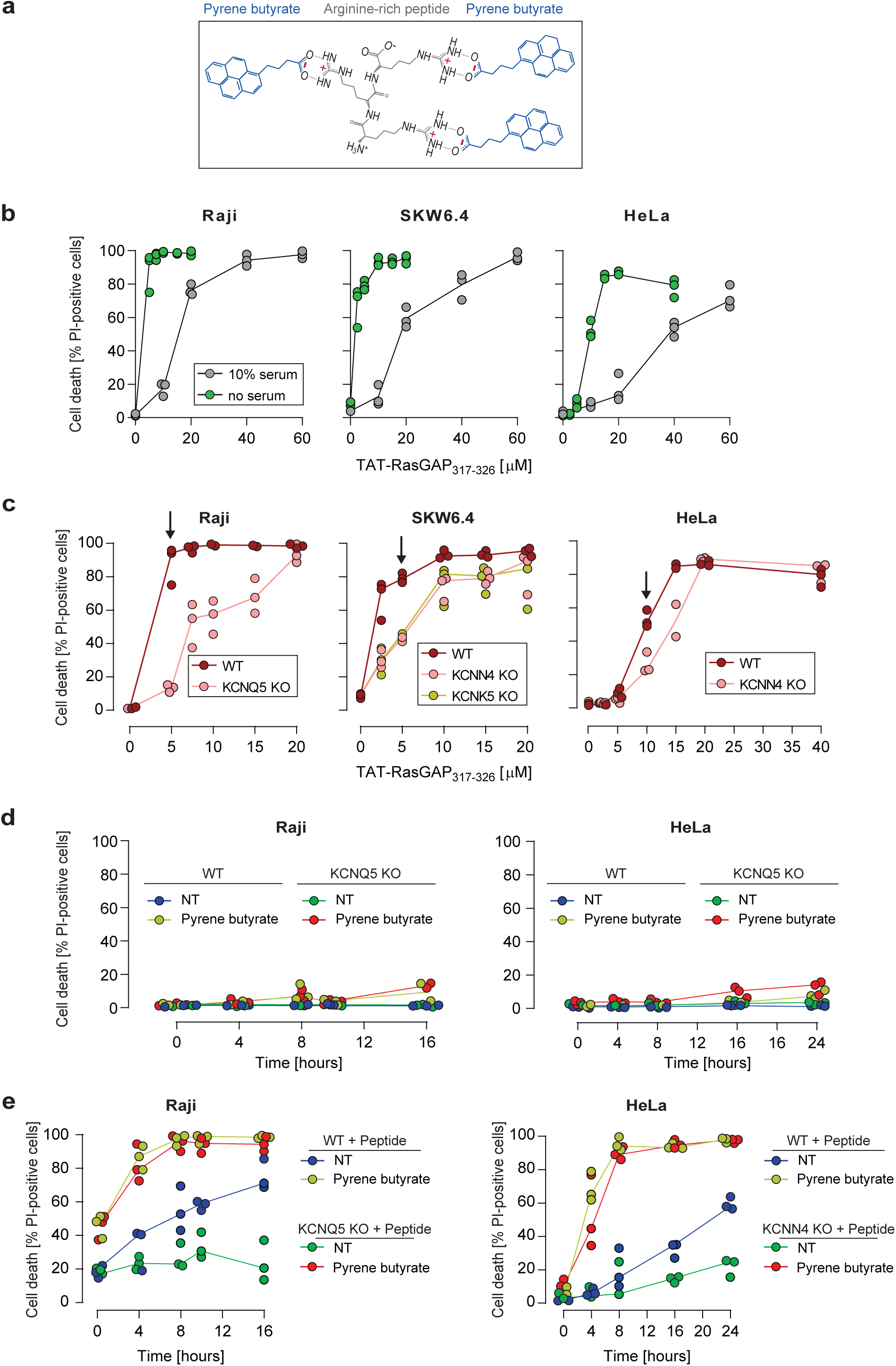

**Supplementary Fig. 7.**
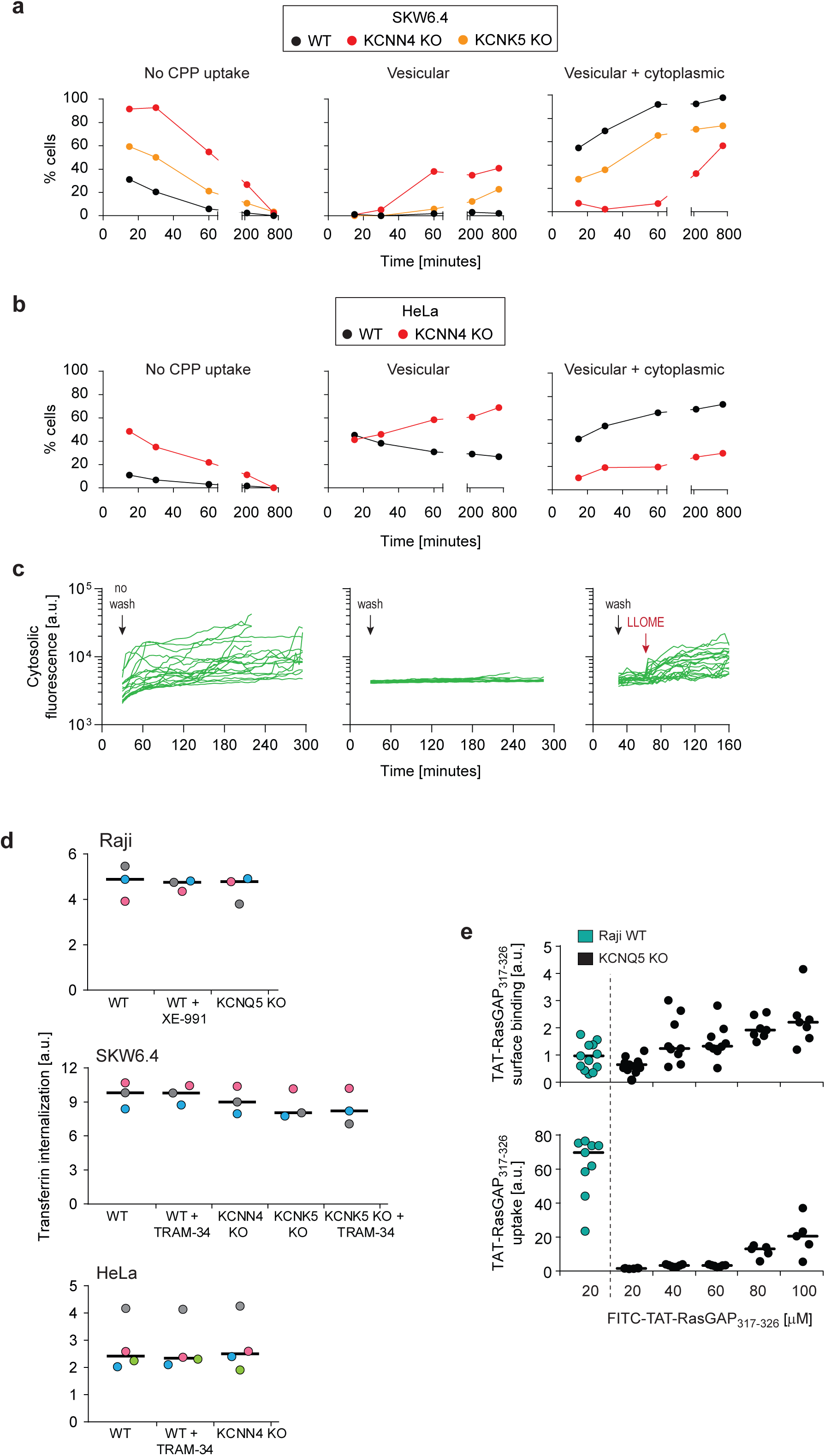

**Supplementary Fig. 8.**
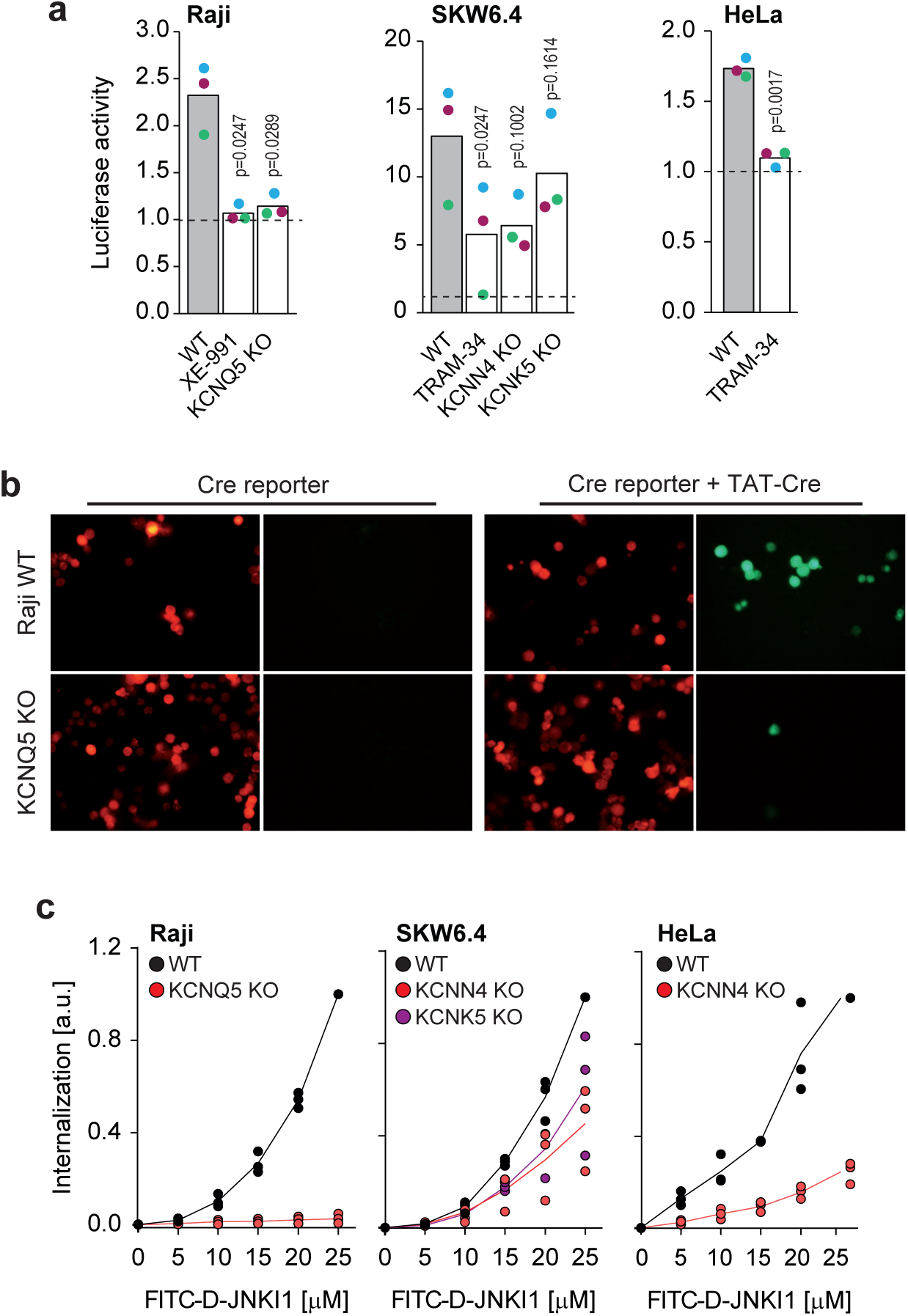

**Supplementary Fig. 9.**
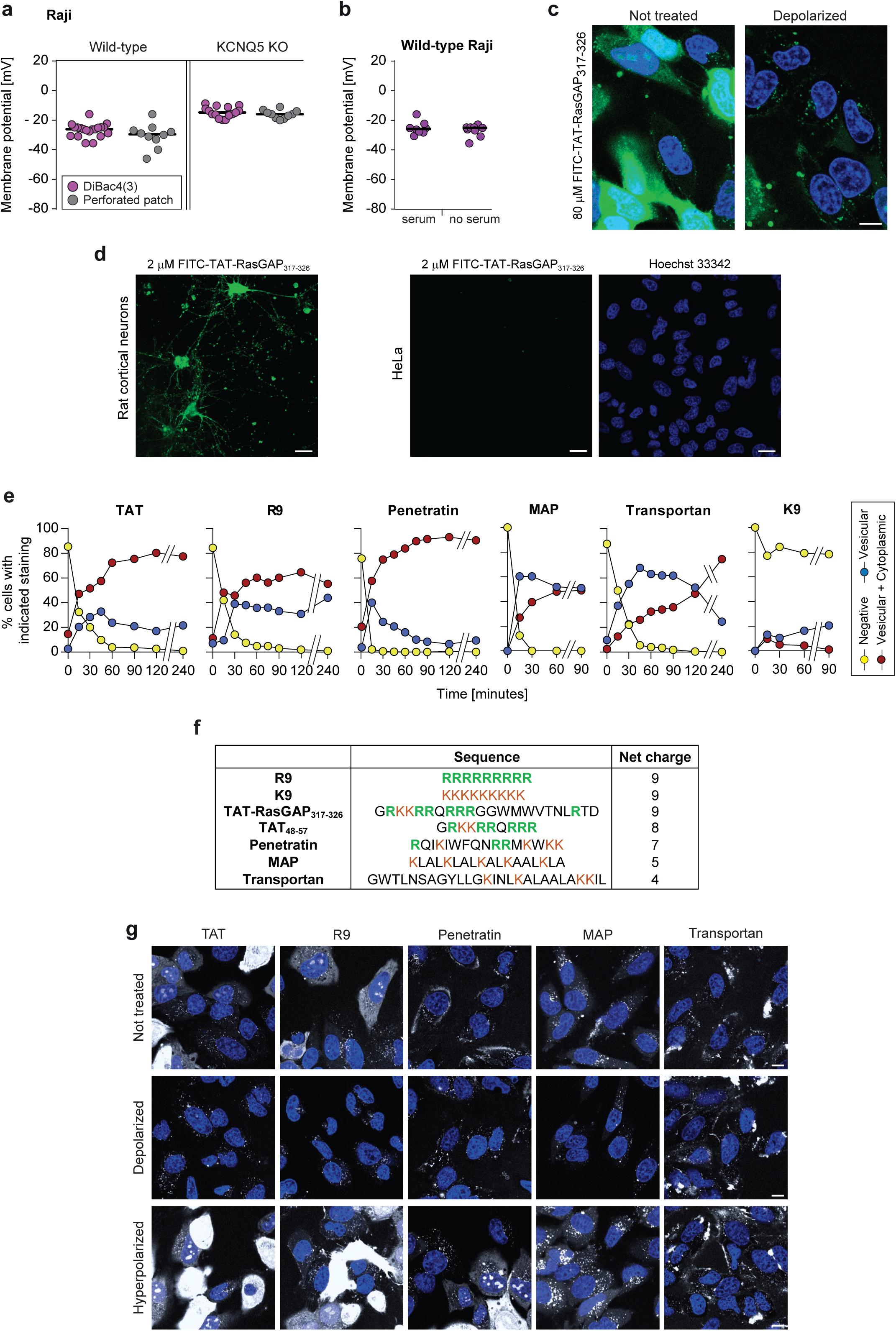

**Supplementary Fig. 10.**
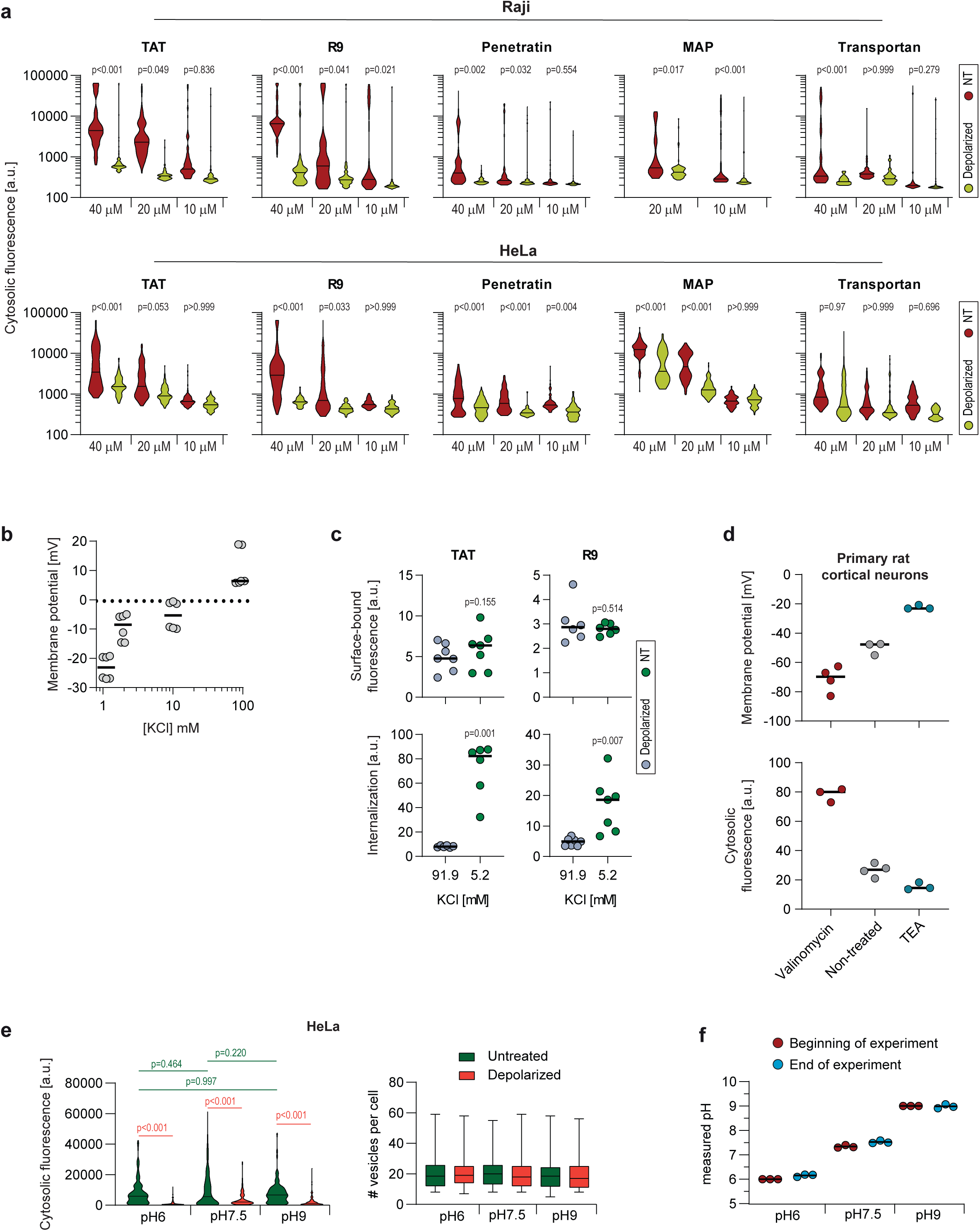

**Supplementary Fig. 11.**
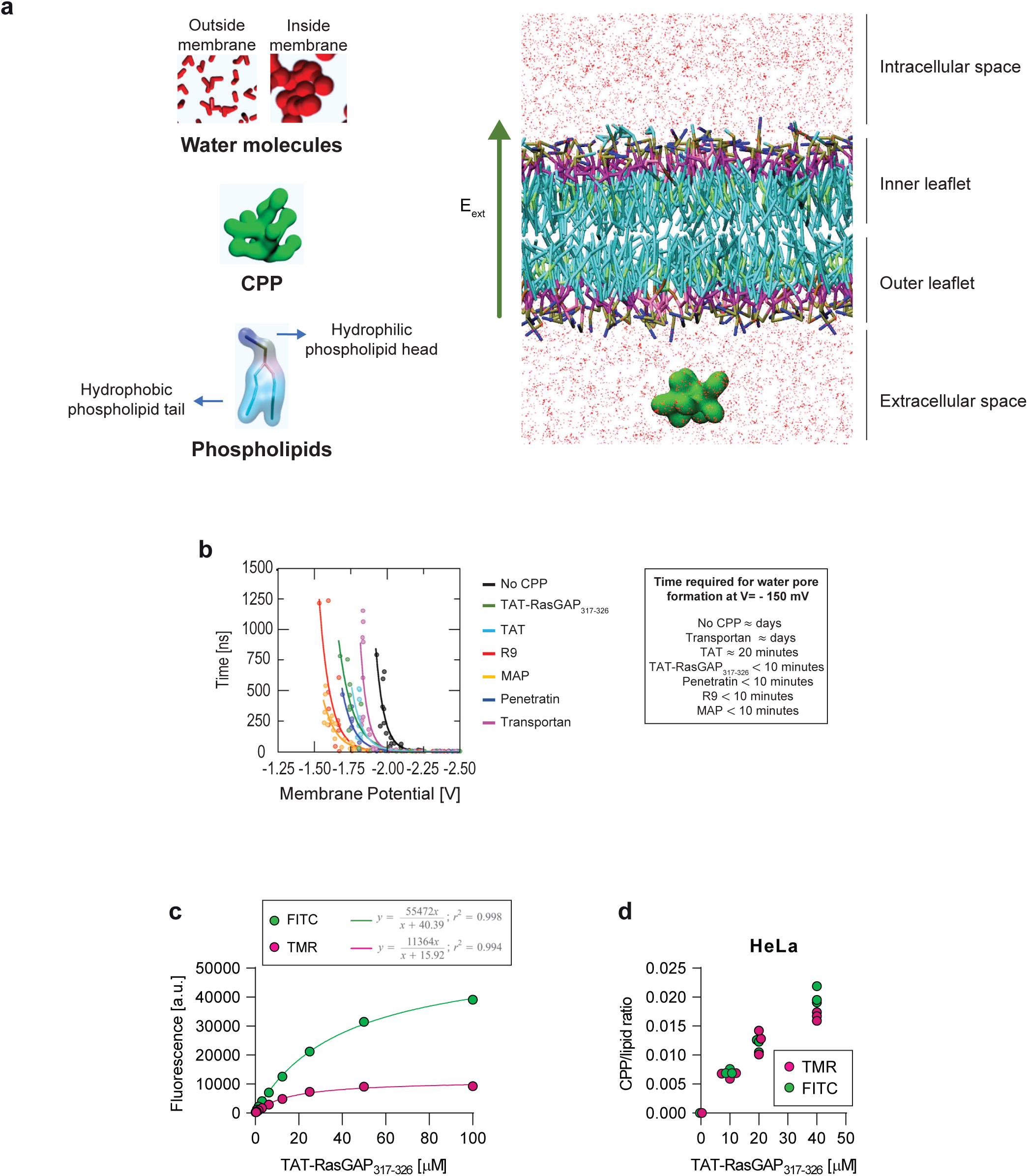

**Supplementary Fig. 12.**
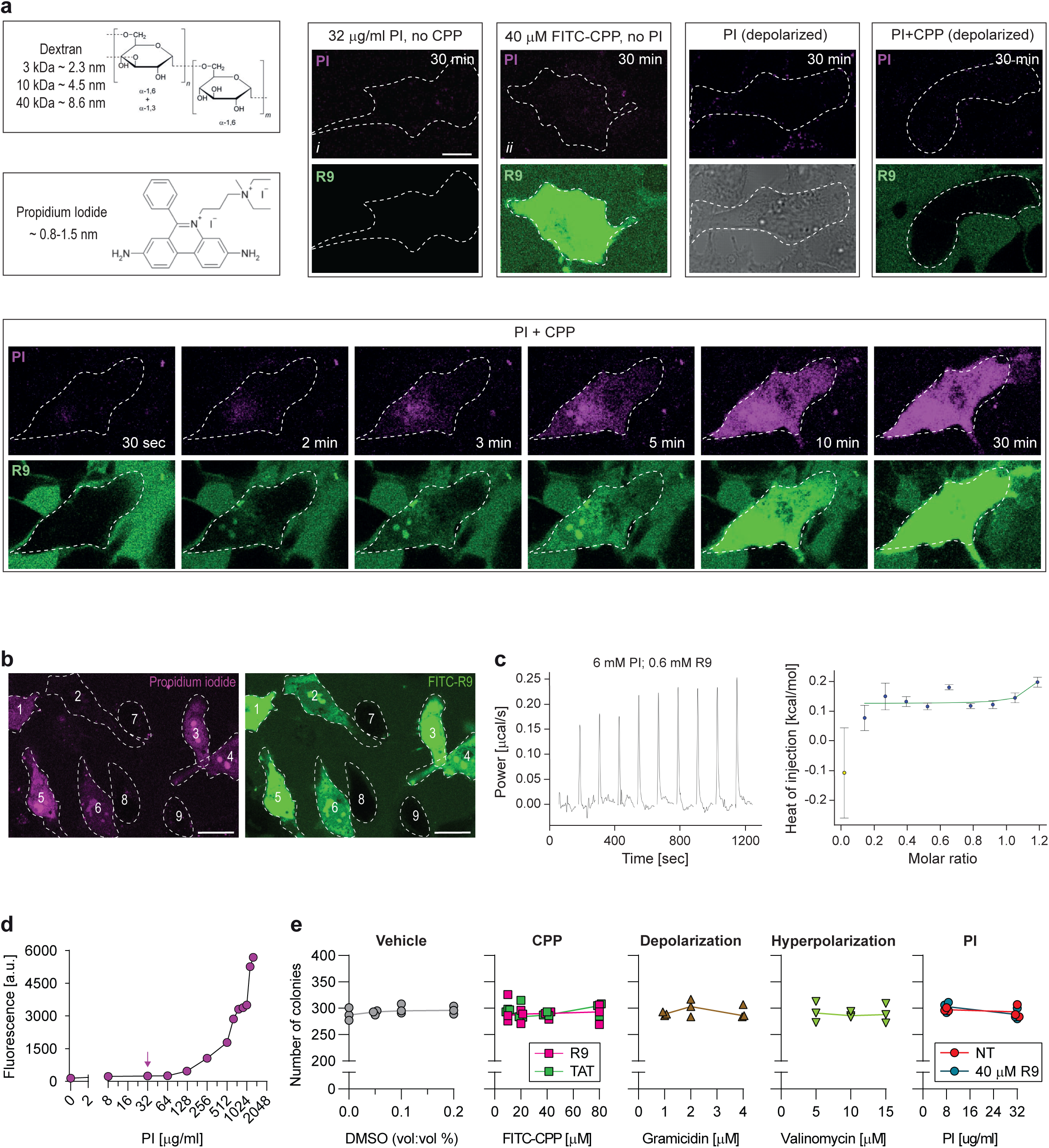

**Supplementary Fig. 13.**
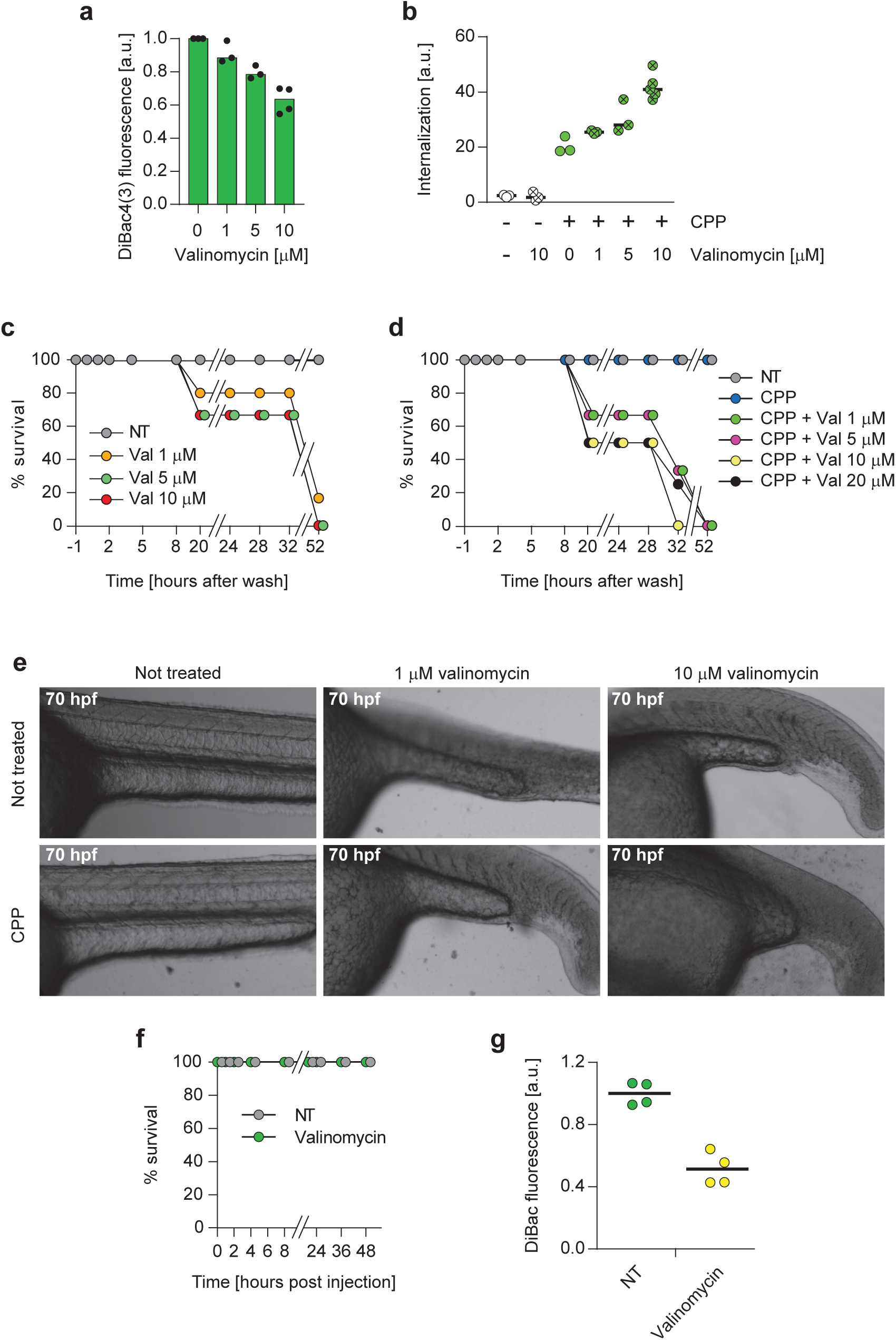

**Supplementary Fig. 14.**
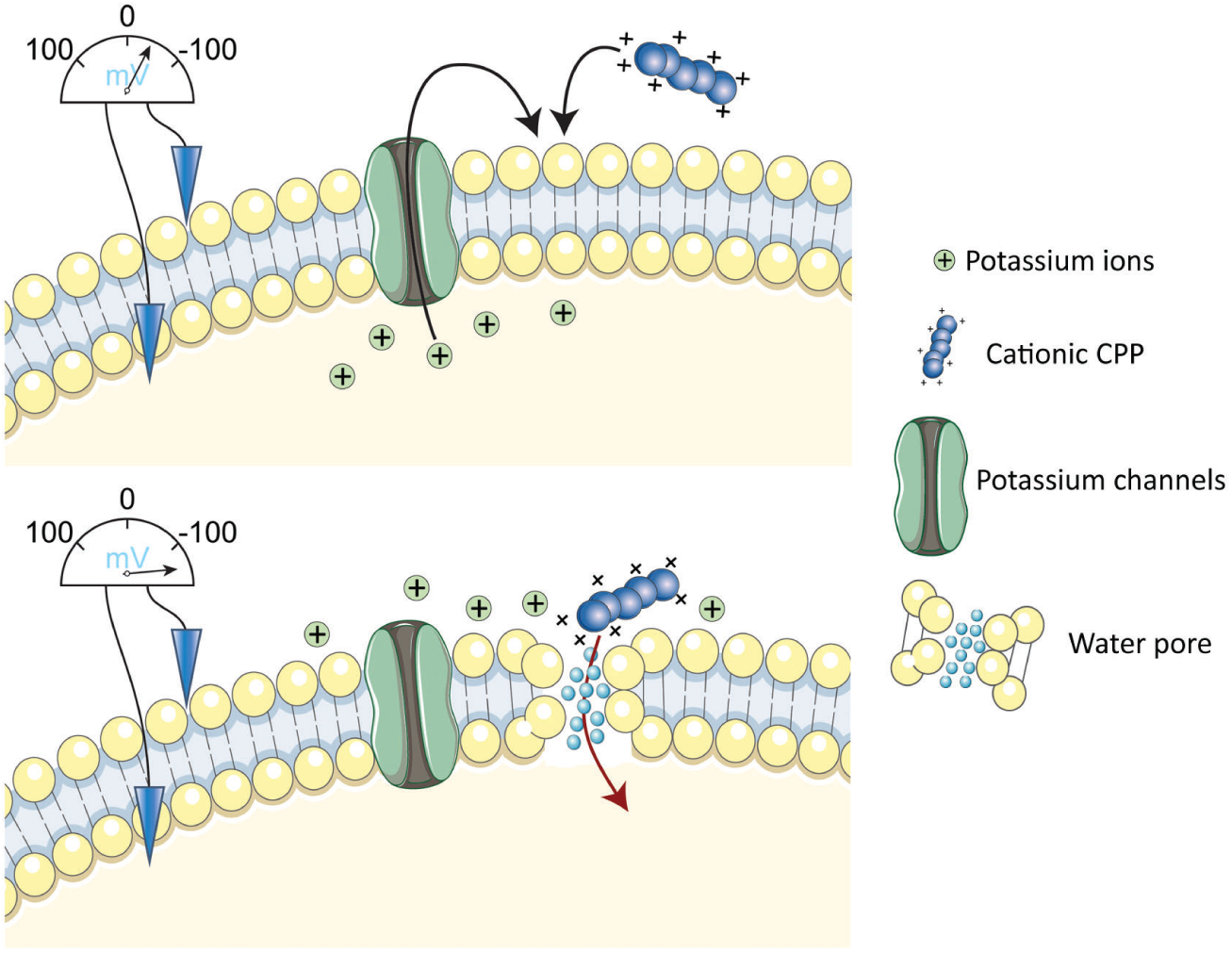

**Supplementary Fig. 15.**
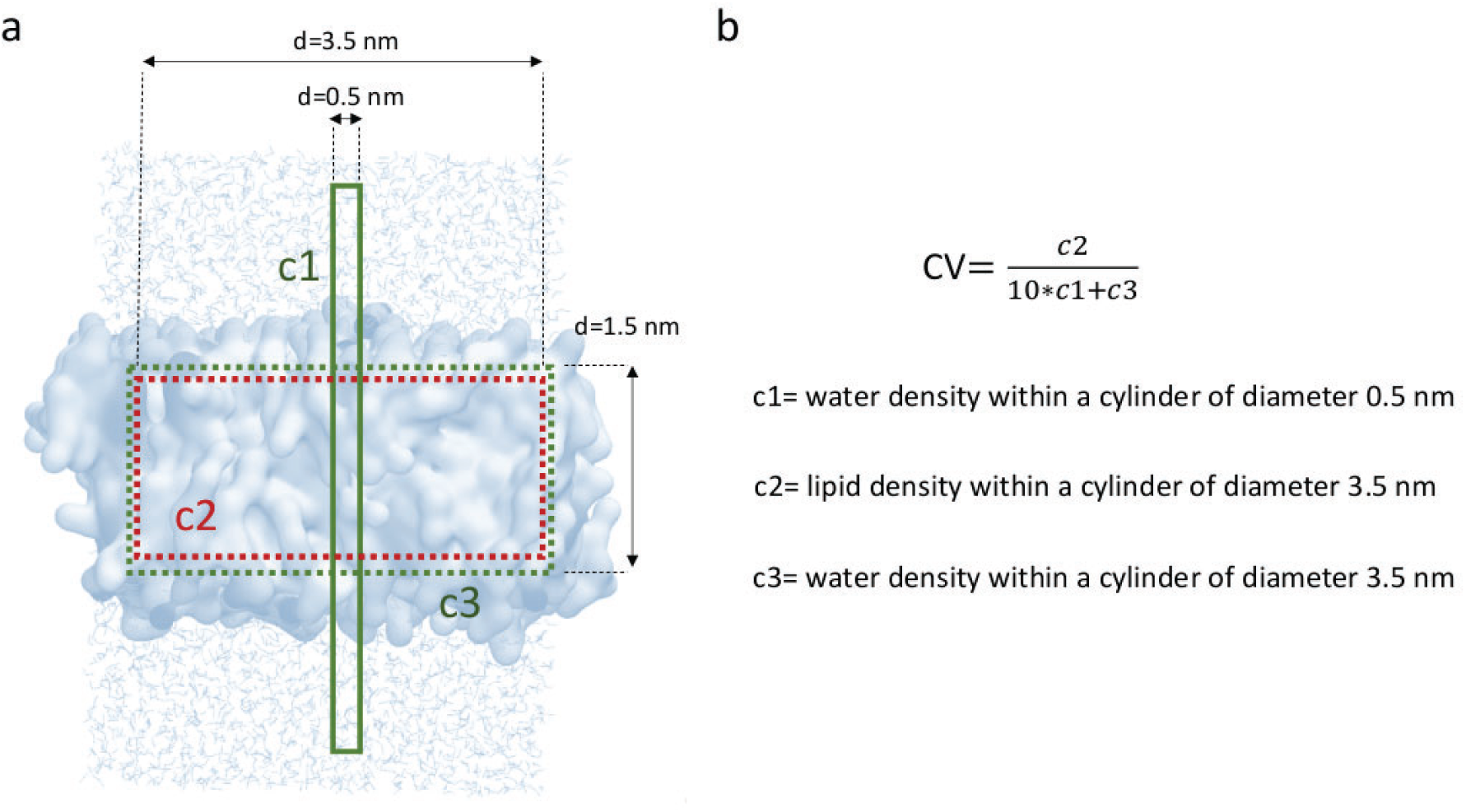

**Supplementary Fig. 16.**
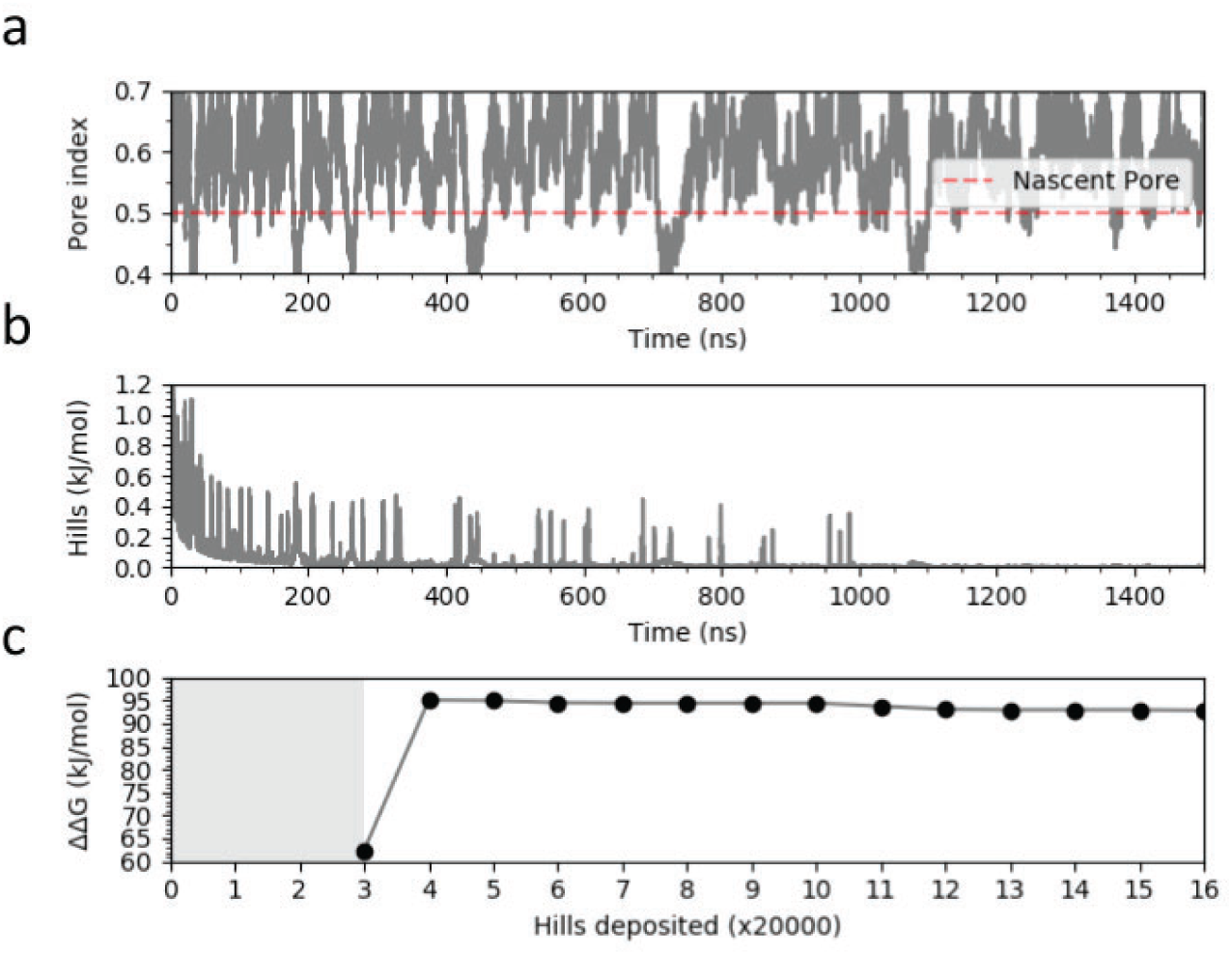

**Supplementary Fig. 17.**
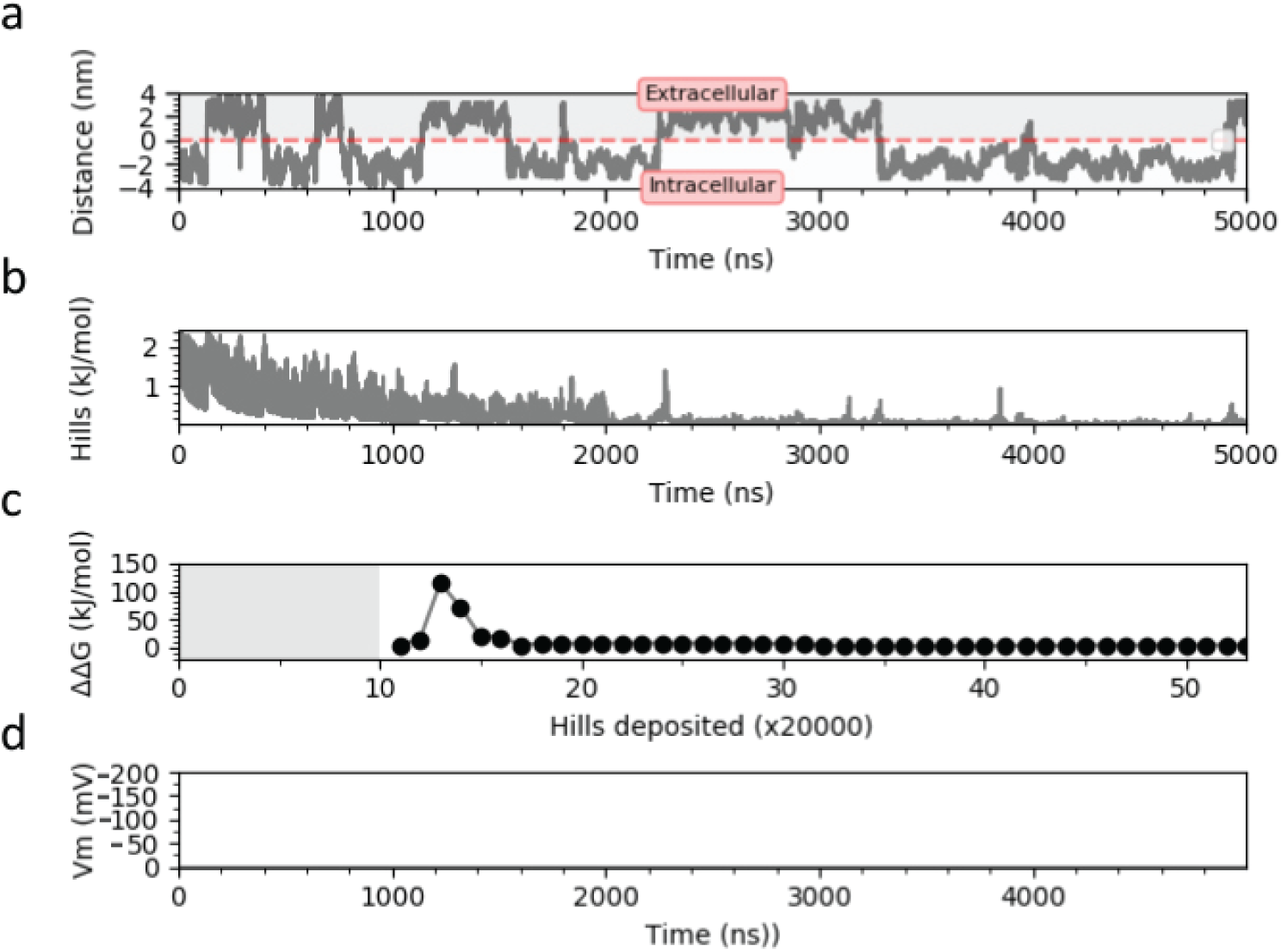

**Supplementary Fig. 18.**
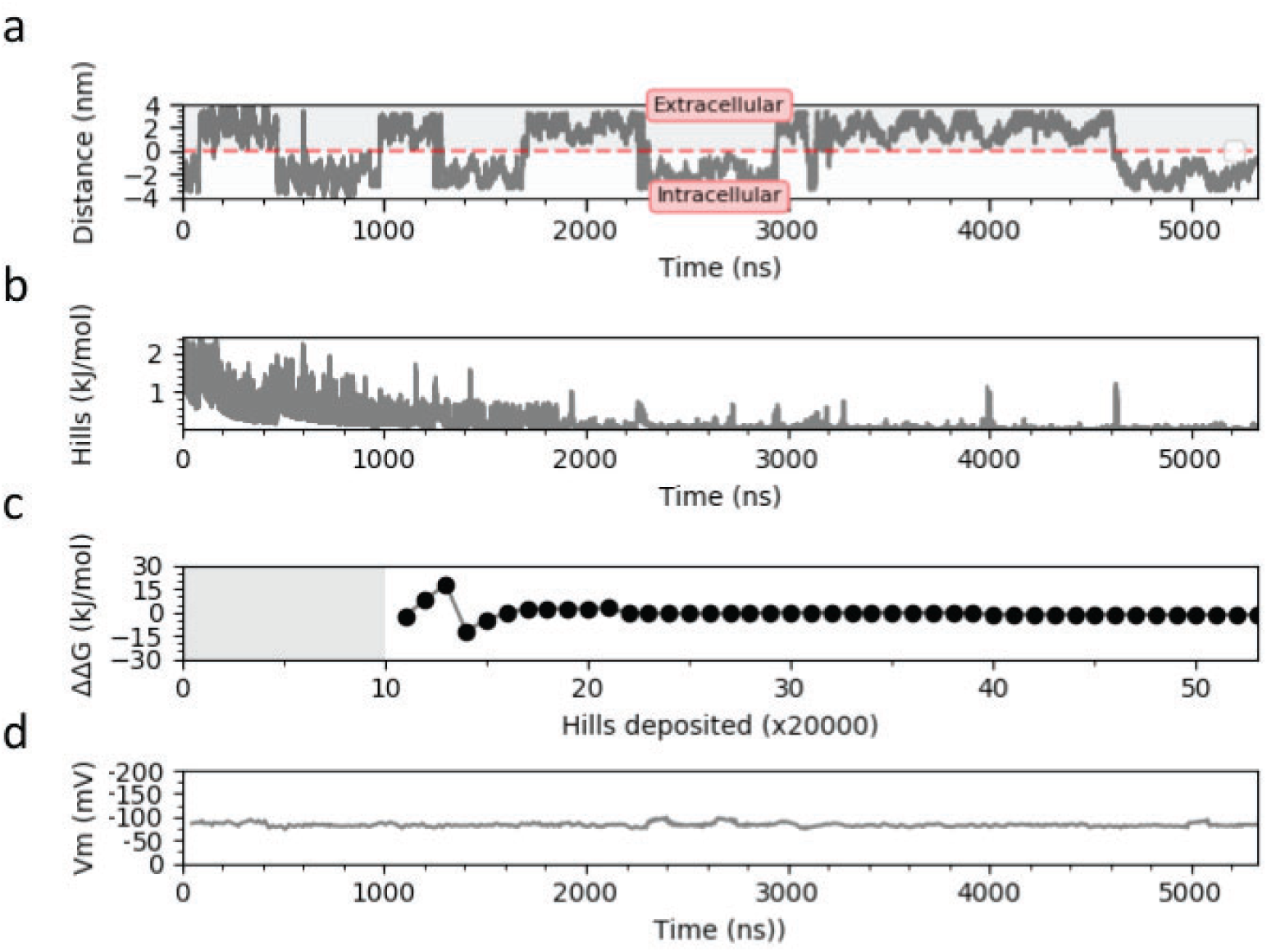

**Supplementary Fig. 19.**
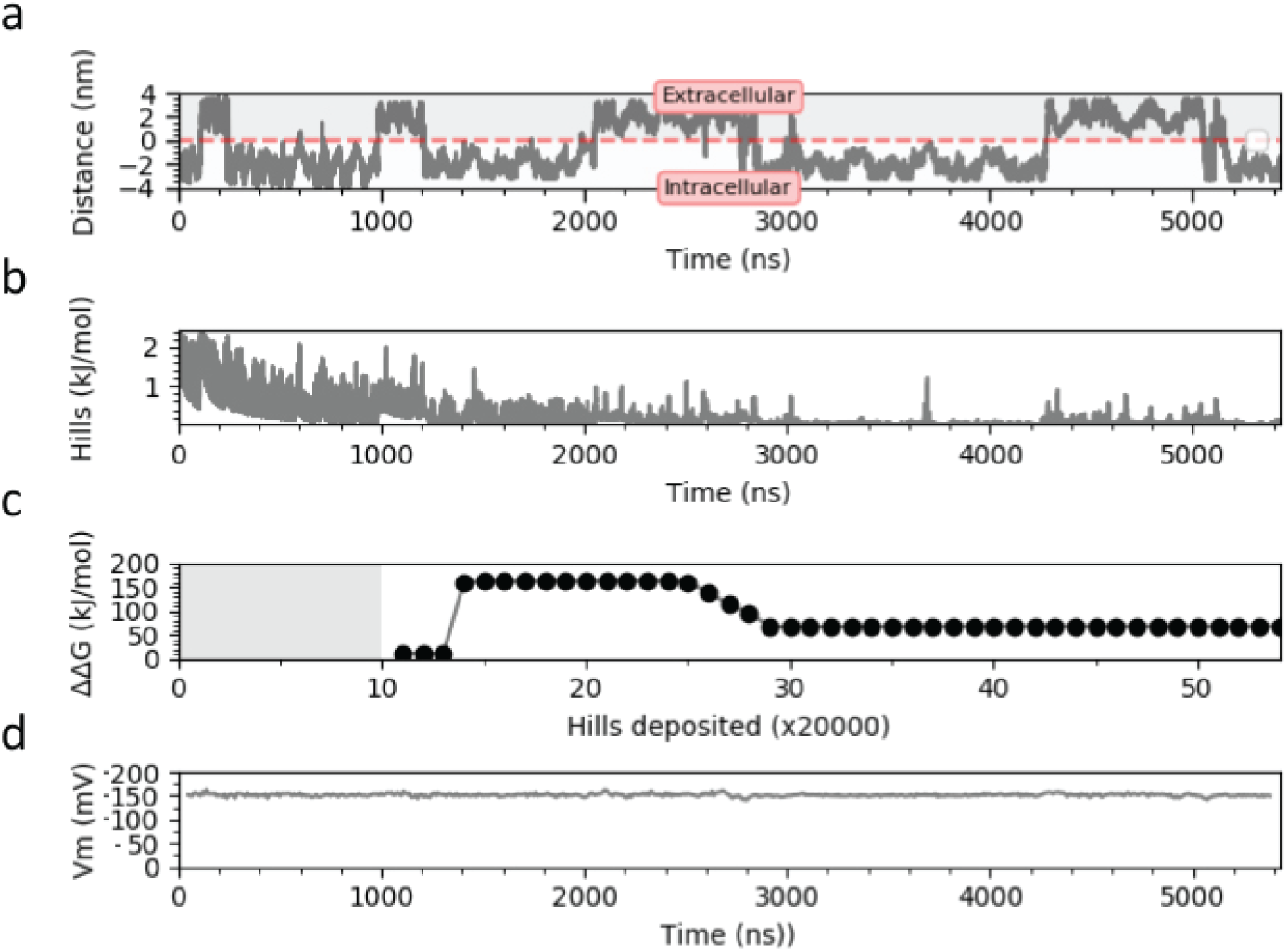

