## Supplementary Information for "Genetic, cellular and structural characterization of the membrane potential-dependent cell-penetrating peptide translocation pore"

#### Effect of serum on cell sensitivity to TAT-RasGAP$_{317\text{-}326}$ and CPP internalization

Serum removal sensitized cells to TAT-RasGAP$_{317\text{-}326}$-induced death (Supplementary Fig. 6a); therefore, the peptide concentrations had to be adapted to perform the experiments in the absence of serum to reach a similar biological activity (Supplementary Fig. 6b) and CPP uptake in cells. Treatment with pyrene butyrate (Supplementary Fig. 6) and valinomycin (Fig. 2) was used in the absence of serum as the latter is expected to interfere with the drug[1]. Serum-removal did not affect the cell membrane potential (Supplementary Fig. 9b) and the cytosolic internalization results were very similar between the experiments performed in the presence or in the absence of serum, once the CPP concentrations were adjusted (Fig. 2c and Supplementary Fig. 10a). Our results are in line with the previous observations that cationic CPPs interact with proteins present in serum[2,3], which results in lower CPP availability in the media.

#### Evaluation of water pore formation free energy though MARTINI coarse-grained models

The metadynamics protocol applied in this study has been validated by estimating the water pore formation free energy in a well-studied[4,5] symmetric DOPC membrane. Each layer contained 100 lipids. The membrane was solvated with 3’000 water molecules, obtaining a molecular system of 11’500 particles The MARTINI force field[6,7] was used to define phospholipids’ topology through a coarse-grained (CG) approach. The polarizable water model has been used to assess the water topology[8]. The molecular system has been minimized by a steepest descent protocol, and then equilibrated through five MD simulations of 1 ns each under the NPT ensemble.

Position restraints were applied during the first three molecular dynamics (MD) simulations and gradually removed, from 200 kJ/mol*nm$^2$ to 10 kJ/mol*nm$^2$. Velocity rescaling[9] temperature coupling algorithm and time constant of 1.0 ps were applied to keep the temperature at 310.00 K. Berendsen[10] semi-isotropic pressure coupling algorithm with reference pressure equal to 1 bar and time constant 5.0 ps was employed. Electrostatic interactions were calculated by applying the particle-mesh Ewald (PME)[11] method and van der Waals interactions were defined within a cut-off of 1.2 nm. Periodic boundary conditions were applied in all directions. A metadynamics[12,13] protocol was applied to estimate the free energy of water pore formation. The lipid/water density index has been selected as collective variable (Supplementary Fig. 15). The well-tempered metadynamics simulations were computed using GROMACS 2019.4 package and the PLUMED 2.5. open-source plug-in. Gaussian deposition rate of 1.2 kJ/mol every 5 ps was initially applied and gradually decreased on the basis of an adaptive scheme, with a bias factor of 50. Gaussian widths of 0.5 was applied following a well-established scheme[12,14-17]. In particular, the Gaussian width value was of the same order of magnitude as the standard deviation of the distance CV, calculated during unbiased simulations. Each system was simulated (with a 20 fs time step) until convergence was reached. The well-tempered metadynamics simulations were computed using GROMACS 2019.4 package[18] and the PLUMED 2.5 open-source plug-in[19] .The reconstruction of the free-energy surface was performed by the reweighting algorithm procedure[20], allowing the estimation of the free energy landscape. Further details about the convergence of the metadynamics simulation of a full DOPC membrane is reported in Supplementary Fig. 16. The comparison between the water pore formation free energy estimated by our MARTINI coarse-grained simulations and previous estimations available in literature is reported in Table S5.

#### In silico pore formation kinetics though Martini coarse-grained simulations

The water pore formation kinetics has been investigated in this study by applying a constant electrostatic potential to the molecular system[21-29]. An asymmetric multi-component membrane was constructed and solvated using CHARMM-GUI[30,31]. Each layer contained 100 lipids (Table S2), in a previously described composition[32]. The membrane was solvated with 3’000 water molecules, obtaining a molecular system of 11’500 particles. The MARTINI[6,7] force field was used to define phospholipids’ topology through a coarse-grained (CG) approach. The polarizable water model has been used to model the water topology[8]. The elastic network ELNEDYN[33] has been applied to reproduce the structural and dynamic properties of the CPPs. For each molecular system, one CPP was positioned 3 nm far from the membrane outer leaflet, in the water environment corresponding to the extracellular space. Then, the system was equilibrated through four MD simulations of 1 ns under the NPT ensemble. Position restraints were applied during the first three MD simulations ang gradually removed, from 200 kJ/mol*nm$^2$ to 10 kJ/mol*nm$^2$. Velocity rescaling[9] temperature coupling algorithm and time constant of 1.0 ps were applied to keep the temperature at 310.00 K. Berendsen[10] semi-isotropic pressure coupling algorithm with reference pressure equal to 1 bar and time constant 5.0 ps was employed. Then, all systems were simulated for the production run in the NPT ensemble with the time step of 20 fs. Electrostatic interactions were calculated by applying the particle-mesh Ewald (PME)[11] method and van der Waals interactions were defined within a cut-off of 1.2 nm. Periodic boundary conditions were applied in all directions. Trajectories were collected every 10 ps and the Visual Molecular Dynamics (VMD)[34] package was employed to visually inspect the simulated systems. GROMACS 2018 was used for simulations and data analysis[18]. The relatively small size of the molecular system and the application of Coarse-Grained Martini forcefield, allowed us to study the pore formation kinetics, requiring many simulations at varying field strengths. In detail, 25 simulations were performed for each molecular system at different external electric field strengths from 0.0055 V/nm to 0.090 V/nm. In the MD simulations, an external electric field $E_{ext}$ was applied parallel to the membrane normal *z*, i.e., perpendicular to the bilayer surface. This was achieved by including additional forces $F_i = q*E_{ext}$ acting on all charged particles *i.* All the MD simulations were performed until the water pore formation event was observed. In order to determine the effective electric field in simulations, we applied a computational procedure reported in literature[28]. The results are reported in Supplementary Figure 11.

### Supplementary Tables

##### Table S1: Media composition

| CAS Number | Components | Quantity in g/l |
|---|---|---|
| 13477-34-4 | Calcium Nitrate Tetrahydrate | 0.10000000 |
| 7487-88-9 | Magnesium Sulfate Anhydrous | 0.04884000 |
| 50-99-7 | D-Glucose Anhydrous | 2.00000000 |
| 56-40-6 | Glycine | 0.01000000 |
| 39537-23-0 | L-Alanyl-L-Glutamine | 0.44600000 |
| 74-79-3 | L-Arginine Free Base | 0.20000000 |
| 70-47-3 | L-Asparagine Anhydrous | 0.05000000 |
| 56-84-8 | L-Aspartic acid | 0.02000000 |
| 30925-07-6 | L-Cystine Dihydrochloride | 0.06520000 |
| 56-86-0 | L-Glutamic Acid | 0.02000000 |
| 71-00-1 | L-Histidine | 0.01500000 |
| 51-35-4 | L-Hydroxy-L-Proline | 0.02000000 |
| 73-32-5 | L-Isoleucine | 0.05000000 |
| 61-90-5 | L-Leucine | 0.05000000 |
| 657-27-2 | L-Lysine Monohydrochloride | 0.04000000 |
| 63-68-3 | L-Methionine | 0.01500000 |
| 63-91-2 | L-Phenylalanine | 0.01500000 |
| 147-85-3 | L-Proline | 0.02000000 |
| 56-45-1 | L-Serine | 0.03000000 |
| 72-19-5 | L-Threonine | 0.02000000 |
| 73-22-3 | L-Tryptophan | 0.00500000 |
| 69847-45-6 | L-Tyrosine Disodium Salt Dihydrate | 0.02883000 |
| 72-18-4 | L-Valine | 0.02000000 |
| 67-48-1 | Choline Chloride | 0.00300000 |
| 58-85-5 | D-Biotin | 0.00020000 |
| 137-08-6 | D-Ca Pantothenate | 0.00025000 |
| 59-30-3 | Folic Acid | 0.00100000 |
| 87-89-8 | Myo-Inositol | 0.03500000 |
| 98-92-0 | Nicotinamide (Nicotinic acid amide) | 0.00100000 |

| | | |
|---|---|---|
| 150-13-0 | P-Aminobenzoic Acid (PABA) | 0.00100000 |
| 58-56-0 | Pyridoxine Hydrochloride | 0.00100000 |
| 83-88-5 | Riboflavin | 0.00020000 |
| 67-03-8 | Thiamine Hydrochloride | 0.00100000 |
| 68-19-9 | Vitamine B12 | 0.00000500 |
| 70-18-8 | L-Glutathione Reduced | 0.00100000 |
| 34487-61-1 | Phenol Red Sodium Salt | 0.00530000 |
| WATER | | 996.66117500 |

#### Table S2. Phospholipid composition considered in the present study

| | Inner membrane | | Outer membrane | |
|---|---|---|---|---|
| type | Number of lipids | % | Number of lipids | % |
| POPC | 18 | 18 | 39 | 39 |
| POPE | 27 | 27 | 6 | 6 |
| PSM | 10 | 10 | 21 | 21 |
| POPS | 11 | 11 | - | - |
| POPI | 5 | 5 | - | - |
| CHOL | 29 | 29 | 34 | 34 |
| total | 100 | 100 | 100 | 100 |

#### Table S3. Primer sequences for the second PCR

| | |
|---|---|
| F2a | AATGATACGGCGACCACCGAGATCTACACTCTTTCCCTACACGACGCTCTTCCGATCTTCTTGTGGAAAGGACGAAACACCG |
| F2b | AATGATACGGCGACCACCGAGATCTACACTCTTTCCCTACACGACGCTCTTCCGATCTAGCTCTTGTGGAAAGGACGAAACACCG |
| F2c | AATGATACGGCGACCACCGAGATCTACACTCTTTCCCTACACGACGCTCTTCCGATCTCGAGCTCTTGTGGAAAGGACGAAACACCG |
| F2d | AATGATACGGCGACCACCGAGATCTACACTCTTTCCCTACACGACGCTCTTCCGATCTCATAACCTCTTGTGGAAAGGACGAAACACCG |
| F2e | AATGATACGGCGACCACCGAGATCTACACTCTTTCCCTACACGACGCTCTTCCGATCTGTGCTAACGTCTTGTGGAAAGGACGAAACACCG |
| R2_iC_26 | CAAGCAGAAGACGGCATACGAGAT**GCTCAT**GTGACTGGAGTTCAGACGTGTGCTCTTCCGATCTTCTACTATTCTTTCCCCTGCACTGT |
| R2_iA_12 | CAAGCAGAAGACGGCATACGAGAT**TACAAG**GTGACTGGAGTTCAGACGTGTGCTCTTCCGATCTTCTACTATTCTTTCCCCTGCACTGT |

#### Table S4. List of sgRNAs used to disrupt target genes

| Target gene | sgRNA name | sgRNA sequence |
|---|---|---|
| KCNN4 | sgKCNN4.1 | CTGCCCGAGTGCTACAAGAA |
| KCNN4 | sgKCNN4.2 | CATGGTGCCCGGCACCACGT |
| KCNK5 | sgKCNK5.1 | ATGGTGGTAATGACGGTCGC |
| KCNK5 | sgKCNK5.2 | CTCTGCCTGACGTGGATCAG |
| KCNQ5 | sgKCNQ5.1 | TCTAGGAATTAATTCACAGC |
| KCNQ5 | sgKCNQ5.2 | ACAGATCCTCCGCATGGTCG |

**Table S5. Comparison of the nascent water pore free energy estimation**

| Force field | Free energy difference (kJ/mol) | References |
|---|---|---|
| All-atom (CHARMM36) | 95.4 | Ting, C.L. et al.[5] |
| All-atom (CHARMM36) | 104.6 (by extrapolation) | Dixit, M. et al.[4] |
| CG-MARTINI | 93+/-1.5 | This study |

**Table S6. List of oligos used for sequencing after TA cloning**

| Target gene | Oligo name | Oligo sequence |
|---|---|---|
| KCNN4 | KCNN4.1.TA.F | GCAGAGAAGCACGTGCA |
| KCNN4 | KCNN4.1.TA.R | GGCAGCATGAGACTCCTTCC |
| KCNQ5 | KCNQ5.1.TA.F | GGGACATGATGTACAATGGA |
| KCNQ5 | KCNQ5.1.TA.F | CCAGAGAGCATCTGCATATG |

#### Supplementary Figure legends

**Supplementary Fig. 1: Characteristics of TAT-RasGAP$_{317-326}$ internalization**

**a**, Depiction of the different modes of CPP entry into cells. Confocal microscopy was performed on the indicated cell lines incubated for one hour with 40 μM FITC-TAT-RasGAP$_{317-326}$ in RPMI, 10% FBS. Cells were washed with PBS prior to visualization. Vesicular staining is indicative of CPP endocytosis while diffuse cytosolic staining is a consequence of CPP direct translocation into cells. Scale bar: 10 μm. **b**, Quantitation of the different modes of CPP entry as a function of time (FITC-TAT-RasGAP$_{317-326}$ continually present in the media) using the experimental conditions presented in panel a. Types of staining were visually quantitated as indicated in panel c (n>150 cells per condition). There was no indication of fluorescence quenching due to endosomal acidification in at least the first hour of CPP exposure (see panel d). Results correspond to the average of three independent experiments. **c**, Visual attribution of the various types of staining associated with FITC-CPP internalization in wild-type Hela cells. Cells 1-9 took up the CPP through both direct translocation and endocytosis (cells 1-4 and cells 5-9 displaying strong and weak cytosolic signal, respectively); cells 10-15 acquired FITC-TAT-RasGAP$_{317-326}$ in vesicles only, no cytosolic staining being detected. The experimental conditions are those of panel a. Scale bar: 20 μm. **d**, Potential effect of fluorophore quenching in CPP-containing endosome detection. Wild-type HeLa cells were incubated for one hour at 37°C in the presence of 40 μM TAT-RasGAP$_{317-326}$ labeled with either FITC (susceptible to quenching at low pH) or TMR (not quenched at low pH) fluorophores. The number of vesicles was visually determined based on confocal images. There were no fewer vesicles detected in cells when the peptide was labelled with FITC than when it was labelled with TMR. **e**, Representative confocal images of focal and homogenous cytosolic entry of 80 μM FITC-TAT-RasGAP$_{317-326}$ in HeLa cells over time. **f**, Quantitation in Raji and HeLa cells of the patterns described in panel e. An apparent homogenous type of entry could be focal if the entry point is outside of the confocal plane. Focal diffusion is likely therefore to be underestimated. The results correspond to three independent experiments. **g**, Cytosolic acquisition of fluorophore-labelled CPPs is not a consequence of laser-induced cellular damage. HeLa cells were incubated with 80 μM FITC-TAT-

RasGAP$_{317\text{-}326}$ for 30 minutes with the indicated frequencies of laser exposures (n>130 cells per condition). Quantitation of the cytosolic fluorescence intensity is shown on the left and the percentage of cells with cytosolic fluorescence is shown on the right. FITC-TAT-RasGAP$_{317\text{-}326}$ translocation into cells occurred similarly after 30 minutes whether the cells were illuminated once or 60 times over the 30 minutes period. Even if one argues that a single illumination is sufficient to cause the internalization seen after a 30 minute incubation period with the CPP, then the internalization should be the same whether the illumination is performed at 1 minute or at 30 minutes. This is not what is observed: the internalization of the FITC-labelled CPP increases overtime independently of the number of times the cells are illuminated. The results correspond to three independent experiments. **h**, Cells can concentrate CPPs in their cytosol. Wild-type Raji and HeLa cells were incubated with 40 µM FITC-TAT-RasGAP$_{317\text{-}326}$ for one hour at 37°C in RPMI, 10% FBS. Images were acquired with a confocal microscope. TAT-RasGAP$_{317\text{-}326}$ fluorescence quantitation was performed using ImageJ using region of interest within the cytosol of cells that had acquired the peptide through direct translocation. The dotted line represents the fluorescence in the cell culture media (i.e. outside of the cells). **i**, Quantitation of FITC-TAT-RasGAP$_{317\text{-}326}$ internalization at the indicated temperatures. Cells were incubated on a Thermoblock for one hour. Peptide internalization was measured by flow cytometry in the presence of 0.2% trypan blue to quench the surface-bound fluorescence. The results for each cell line correspond to three independent experiments. The data was normalized to the fluorescence at 37°C (dashed line). **j**, Assessment of the contribution of surface bound FITC-TAT-RasGAP$_{317\text{-}326}$ to overall cell-acquired fluorescence. HeLa cells were incubated with 80 µM FITC-TAT-RasGAP$_{317\text{-}326}$ for 60 minutes in 24 well plates in 0.5 ml RPMI, 10% FBS. Then, cells were washed once with PBS (1 ml), trypsinized, and resuspended in 0.5 ml PBS containing or not 0.2% trypan blue. Cell-associated fluorescence was recorded by flow cytometry. Data were normalized to the fluorescence values of cells incubated with FITC-TAT-RasGAP$_{317\text{-}326}$ in the absence of trypan blue. The results correspond to the average of three independent experiments. The efficiency of trypan blue-mediated FITC quenching is presented in the next panel. **k**, Quantitation, in the absence of cells, of the emitted FITC fluorescence at 525 nm (excitation at 488 nm) of FITC-TAT-RasGAP$_{317\text{-}326}$ at the indicated concentrations in PBS in the presence or in the absence of 0.2% trypan blue. Fluorescence was recorded in 96-well plates using a CYTATION3 apparatus.

**Supplementary Fig. 2: TAT-RasGAP$_{317-326}$ enters cells via endocytosis and direct translocation, but only direct translocation mediates its biological activity**

**a-e,** HeLa cells were incubated with 80 μM FITC-TAT-RasGAP317-326 in RPMI, 10% FBS for the indicated periods of time. Peptide internalization and cell death was assessed by flow cytometry. **a**, Left: representative flow cytometry dot plot showing the gating strategy used in flow cytometry experiments. Right: representative flow cytometry histogram of HeLa cells incubated with FITC-TAT-RasGAP$_{317-326}$ in RPMI, 10% FBS for 60 minutes. The dotted line represents cellular auto-fluorescence. **b**, Variation over time of the low and high intensity peak values. Peptide internalization was assessed by flow cytometry. The results correspond to three independent experiments. Note the different scales used to plot the median values of the low and high intensity populations. **c**, Quantitation of cells with low intensity peptide fluorescence and vesicular staining. Total peptide internalization was assessed by flow cytometry. The percentage of cells with low CPP cytosolic fluorescence was visually quantitated on confocal images (n>150 cells per condition). The results correspond to three independent experiments. **d**, Quantitation of the pattern of appearance of cells with high intensity peptide fluorescence and strong diffuse cytosolic staining. Data assessment as in panel c. The results correspond to three independent experiments. **e**, Kinetics of FITC-TAT-RasGAP$_{317-326}$-induced death in the two populations described in panel (b). Cell death was assessed by propidium iodide (PI) staining. The results correspond to three independent experiments. **f**, Representative flow cytometry histograms of Raji (left) or SKW6.4 (right) cells incubated with 40 μM FITC-TAT-RasGAP$_{317-326}$ for different period of time. The results correspond to one of three independent experiments. Note: concentrations above 40 μM of TAT-RasGAP$_{317-326}$ led to too extensive Raji and SKW6.4 cell death, preventing analysis of peptide internalization at these concentrations.

**Supplementary Fig. 3: CRISPR/Cas9 screen and target gene identification**

**a**, Scheme of the CRISPR/Cas9 screening strategy to identify genes involved in TAT-RasGAP$_{317-326}$ cellular entry. Raji and SKW6.4 cells were infected with a single guide RNA (sgRNA) CRISPR/Cas9 lentiviral library in conditions favoring the expression of only one specific sgRNA per cell (which is indicated by differentially colored cells). The infected cells were then incubated in the presence or in the absence of 40 μM TAT-

RasGAP$_{317-326}$ for 8 days (Raji) or 17 days (SKW6.4). The expression of sgRNAs in both populations was assessed by massive parallel sequencing to determine the enrichment or depletion in specific sgRNAs in the peptide-treated population.

**b**, Scatter-plot depicting the changes in sgRNA expression between the control and treated cell populations. The most significantly modulated sgRNA sets are color-coded.

**Supplementary Fig. 4: Validation of genes involved in the TAT-RasGAP$_{317-326}$ internalization identified through CRISPR/Cas9 screening**

**a-c,** Sequencing of the regions targeted by the sgRNAs disrupting KCNQ5, KCNK5 and KCNN4 potassium channels in Raji (panel a), SKW6.4 (panel b) or HeLa cells (panel c). Mutations, insertions and deletions induced by the CRISPR/Cas9 system are shown in red. Except the cases mentioned below, these mutations induce a frame shift and early termination of the open reading frame. The sequences targeted by the sgRNAs are highlighted in yellow. The sgRNA-induced mutations in the first allele of clone Q5-2, in the first allele of SKW6.4 cells clone N4-2, and in the first allele of clone K5-2 do not induce a frame-shift but are located in critical domains necessary for potassium channel activity, such as the pore-forming region of KCNK5 (uniprot accession number:O95279), region adjacent to the pore of KCNQ5 (uniprot accession number: Q9NR82) or calcium recognition domain of KCNN4 (uniprot accession number: O15554). Hence, these alleles presumably encode non-functional channels. Clones Q5-1 in Raji, N4-1 and K5-1 in SKW6.4, and N4-1 in HeLa cells were those that were used in subsequent experiments. The blue nucleotides correspond to silent changes, aimed at reducing sgRNA-targeting, introduced in the KCNQ5-encoding lentiviral vector used in panel e. PAM (protospacer adjacent motif) sequences are italicized. **d**, The indicated wild-type (WT) and knock-out (KO) cells were tested for their ability to be killed by TAT-RasGAP$_{317-326}$. Cell death was assessed after 16 hours.
**e**, Wild-type or KCNQ5 knock-out Raji cells were infected or not with a sgRNA-resistant FLAG-KCNQ5-encoding lentivirus and treated with TAT-RasGAP$_{317-326}$ for 16 hours. Expression of FLAG-KCNQ5 construct was detected by western blot using an anti-FLAG antibody. **f**, KCNN4 knock-out (KO) SKW6.4 and HeLa cells were infected or not with a KCNN4-expressing lentivirus and then treated with TAT-RasGAP$_{317-326}$ for 16 hours (SKW6.4) or 24 hours (HeLa). Cell death was assessed by flow cytometry of PI-stained cells. Expression of KCNN4-V5 construct was detected by western blot using an anti-V5 antibody. Note that the Raji, SKW6.4 and HeLa knock-out cell lines

still express Cas9 and the sgRNAs targeting the potassium channels. Hence, the ectopically expressed wild-type channel-encoding cDNAs can be targeted by the CRISPR/Cas9 system but this nevertheless allows for a detectable expression of the FLAG- or V5-tagged constructs although to much lower levels than in similarly infected wild-type cells lacking Cas9 and sgRNAs (see Supplementary Fig. 5). In Raji cells, the re-expressed KCNQ5 channel contained 5 silent changes in the sgRNA recognition site (see panel a) but this proved insufficient to allow for a strong ectopic expression (see Supplementary Fig. 5a, last two lanes). In panels d to f, the results correspond to the average of three independent experiments.

**Supplementary Fig. 5: Uncropped western blots assessing the expression of tagged potassium channels through lentiviral expression**

**a**, Wild-type and KCNQ5 knock-out Raji cells infected or not with FLAG-tagged KCNQ5. **b**, Wild-type and KCNN4 knock-out SKW6.4 cells infected or not with V5-tagged KCNN4. **c**, Wild-type and KCNN4 knock-out HeLa cells infected or not with V5-tagged KCNN4. The bands encompassed by dashed purple rectangles are those shown in Supplementary Fig. 4e-f.

**Supplementary Fig. 6: Potassium channels modulate TAT-RasGAP$_{317-326}$ entry, but not the peptide-induced death**

**a**, Quantitation of concentration-dependent TAT-RasGAP$_{317-326}$ induced death in Raji, SKW6.4 and HeLa cells in the presence or in the absence of serum. The results correspond to the median of three independent experiments. **b**, Quantitation of wild-type (WT) and knock-out (KO) cell death after incubation for 16 hours (Raji and SKW6.4) or 24 hours (HeLa) in serum-free RPMI media in the presence of increasing concentrations of TAT-RasGAP$_{317-326}$. Arrows indicate the chosen peptide concentrations for experiments performed in the absence of serum described in panels c and d. The results correspond to the median of three independent experiments. **c**, Cell death assessment in wild-type (WT) and knock-out (KO) Raji and HeLa cells incubated in RPMI without serum, treated (+ pyrene butyrate) or not (NT) with 50 μM pyrene butyrate for the indicated periods of time. DMSO (0.25% vol:vol) was used as a vehicle control for the condition in the absence of pyrene butyrate. The results correspond to the median of three independent experiments. The scheme on the right depicts the structure of pyrene butyrate and how the molecule interacts with arginine residues found in CPPs.

**d**, Assessment of peptide-induced death at different time points in pyrene butyrate-treated cells lacking or not specific potassium channel. Cells were incubated with 5μM (Raji) or 10 μM (HeLa) of TAT-RasGAP$_{317-326}$, pre-incubated or not with 50 μM pyrene butyrate for 30 minutes (in RPMI without serum). The results correspond to the median of three independent experiments.

**Supplementary Fig. 7: Potassium channels modulate direct CPP translocation, but not endocytosis**

**a**, Same as Fig. 1c, but for wild-type, KCNN4 and KCNK5 SKW6.4 knock-out cells. The results correspond to the average of three independent experiments. **b**, As panel a, but for wild-type and KCNN4 knock-out HeLa cells. The results correspond to the average of three independent experiments. **c**, Quantitation of TAT-RasGAP$_{317-326}$ cytosolic access resulting from direct translocation (left, n=19) and endosomal escape (middle, n=16 and right. n=22) in the presence or in the absence of 1 mM LLOME, a endosome/lysosome disruptor[35]. In direct translocation condition, the peptide was continuously present in the media, while it was washed out after 30 minutes to assess endosomal escape. LLOME was used to show that our experimental setup allows for the detection of cytosolic CPPs if they are released from endocytic vesicles. The results correspond to three independent experiments. **d**, Quantitation by flow cytometry of 20 μg/ml AlexaFluor488-transferrin internalization in the indicated wild-type (WT) cell lines and their corresponding knock-out (KO) versions, pretreated or not for 30 minutes with the XE-991 (10 μM) or TRAM-34 (10 μM) potassium channel inhibitors. Transferrin internalization was allowed to proceed for 60 minutes (still in the presence of inhibitors when these were used in the 30 minute pre-incubation period). To quench membrane bound transferrin fluorescence, cells were incubated with 0.2% trypan blue prior to flow cytometry analysis. The independent experiment replicates are color-coded. **e**, Assessment of FITC-TAT-RasGAP$_{317-326}$ cell surface binding on wild-type and KCNQ5 knock-out Raji cells after 60 seconds of incubation (top), as well as associated peptide internalization after one hour of treatment (bottom). The results correspond to at least five independent experiments.

**Supplementary Fig. 8: Potassium channels regulate the cellular internalization of various TAT-bound cargos**

**a**, TAT-PNA-induced luciferase activity in the indicated cell lines pretreated or not with potassium channel inhibitors (XE-991 or TRAM-34) or genetically invalidated for specific potassium channels. Results are normalized to non-stimulated cells (dashed lines). The independent experiment replicates are color-coded. The p-values correspond to the assessment of the significance of the differences with the control wild-type condition using ANOVA multiple comparison analysis with Dunnett's correction. **b**, Representative microscopy images of wild-type and KCNQ5 knock-out Raji cells expressing loxP-RFP-STOP-loxP-GFP and treated or not with 20 μM TAT-Cre for 48 hours. The results correspond to one of three independent experiments. **c**, Internalization of FITC-D-JNKI1 in the indicated cell lines genetically invalidated (KO) or not (WT) for specific potassium channels. The results correspond to the median of three independent experiments.

**Supplementary Fig. 9: Intracellular internalization of various CPPs**

**a**, Membrane potential measurement validation. Membrane potential of wild-type and KCNQ5 knockout (KO) Raji cells was measured using DiBac4(3), a fluorescent membrane potential sensor, or by performing perforated patch electrophysiology recordings. Each dot in the figures reporting DiBac4(3) measurements corresponds to the median of 10'000 cell recording. For perforated patch, each dot in the figures corresponds to the membrane potential of one cell. **b.** Membrane potential measurement in wild-type Raji cells performed in the presence or in the absence of 10% serum. Membrane potential was assessed using DiBac4(3). **c**, Representative confocal images of wild-type HeLa cells incubated with 80 μM FITC-TAT-RasGAP$_{317\text{-}326}$ for one hour in the presence (depolarized) or in the absence (not treated) of 2 μg/ml gramicidin. Scale bar: 10 μm. **d**, Representative confocal images of primary rat cortical neurons (left) and wild-type HeLa cells (right) incubated for one hour at 37°C with FITC-TAT-RasGAP$_{317\text{-}326}$ in the absence of serum. To highlight the differential capacity of these cells to take up the CPP in cytosol, a low concentration of TAT-RasGAP$_{317\text{-}326}$ (2 μM) was used here. At higher concentrations, this CPP readily enters HeLa cells (see panel c). Post incubation cells were washed and imaged with a confocal microscope using the same laser settings for both cell types. Nuclei of HeLa cells were labelled with Hoechst (blue staining). Scale bar: 20 μm. **e**, Confocal microscopy quantitation of HeLa cells displaying the indicated types of CPP internalization. The CPPs (40 μM, except for MAP 20 μM) were continuously present in the media during the course of the experiment. The results correspond to the average of three independent experiments. **f**, Sequences of the indicated CPPs and their net charge. Positively charged amino acids (arginine and lysine) are color-coded. **g,** Representative confocal images of wild-type HeLa cells incubated with 10 μM of the indicated CPP in the absence of serum in physiological, depolarized (2 μg/ml gramicidin) or hyperpolarized (10 μM valinomycin) conditions. Refer to Fig. 2c for the corresponding quantitation of cytosolic fluorescence and number of CPP-positive endocytic vesicles per cell. Scale bar: 10 μm.

**Supplementary Fig. 10: Effect of membrane potential modulation on CPP internalization.**

**a**, Cytosolic CPP levels in normal or depolarized (2 μg/ml gramicidin) conditions in Raji (top) and HeLa (bottom) cells (n>150) incubated for one hour at 37°C with the indicated CPPs in RPMI with 10% FBS. Cytosolic fluorescence was quantitated using ImageJ based on confocal images by selecting a region within a cell devoid of CPP-containing endosomes. Statistical analysis to compare normal and depolarized conditions at given concentrations was performed using ANOVA test with Tuckey correction for multiple comparisons. **b**, Setting membrane potential by varying extracellular potassium concentrations. Assessment of membrane potential changes in Raji cells incubated in RPMI medium containing the indicated concentrations of potassium chloride (isotonicity was maintained by adapting the sodium chloride concentrations; see methods). Membrane potential was measured with DiBac4(3). The results correspond to the median of 4-6 independent experiments. **c,** Assessment of FITC-CPP binding to cellular membrane of wild-type Raji cells in normal or depolarized conditions after 60 seconds of incubation (top), as well as associated peptide internalization after one hour of treatment (bottom). Cells were preincubated for 30 minutes in the presence of RPMI-media containing 5.2 or 91.9 mM potassium chloride and then treated with 40 μM of the indicated CPPs. The results correspond to at least six independent experiments. Comparison between non-treated or depolarized conditions was done using two-tailed paired t-test. **d**, Assessment of membrane potential (top) and FITC-TAT-RasGAP$_{317-326}$

cytosolic uptake (bottom) in control, depolarized (TEA-treated), and hyperpolarized (valinomycin-treated) primary rat cortical neurons. The results correspond to the median of at least three independent experiments based on confocal images (n>100 cells). **e**, Quantitation of cytosolic TMR-labelled TAT-RasGAP$_{317-326}$ internalization (left) and number of CPP-positive vesicles (right) in HeLa cells (n>150) at the indicated pH in depolarizing and control conditions. These experiments were performed in the presence of 10% FBS as acidification or alkalization were toxic for the cells in the absence of serum. Cells were preincubated in media at the indicated pH in presence or in the absence of 2 μg/ml gramicidin (depolarization), prior to addition of 40 μM TMR-TAT-RasGAP$_{317-326}$ for one hour. The quantitation was performed based on confocal microscopy images using ImageJ. Statistical analysis was performed using ANOVA test with Tuckey correction for multiple comparisons. **g.** The pH values remained stable throughout the duration of the experiment in panel f.

**Supplementary Fig. 11: In silico modeling of CPP direct translocation**

**a**, Schematic depiction of the molecular system used estimate water pore formation kinetics. The static electric field ($E_{ext}$) has been highlighted with a green arrow. Water molecules are shown as red structures (small when outside membranes and large when found within membranes). **b**, Assessment of the time necessary for water pore formation in the presence or in the absence of the indicated CPPs at 37°C based on *in silico* pore formation kinetics experiments. A fitting function [$t=A_0*e^{(A1*V)}$] was used, where t is the simulation time of water pore formation, V is the transmembrane potential, $A_0$ and $A_1$ the fitting coefficients. The Pearson correlation coefficients of the fitting curves are $R_{R9}$=0.85, $R_{Penetratin}$=0.95, $R_{MAP}$=0.92, $R_{TATRasGAP}$=0.91, $R_{Transportant}$=0.80, and $R_{NoCPP}$=0.87. **c**, Fluorescence of FITC- or TMR-labeled TAT-RasGAP$_{317-326}$ as a function of the peptide concentration in 200 μl PBS in 96-well plates using CYTATIOn3 apparatus. The curves were fitted with Michaelis-Menten-like equations). **d**, Fluorescence-based CPP/lipid ratio calculations in HeLa cells (the lipids considered here are those of the plasma membrane). Cells (75'000) were incubated with the indicated concentrations of FITC- or TMR-labeled TAT-RasGAP$_{317-326}$ for one minute at 37°C in 1 ml RPMI, 10% FBS (in this condition, no or only marginal CPP uptake into cells occurs; hence here the CPP signal associated with cells corresponds to cell surface-associated peptides). Cells were then washed on ice twice with ice-cold PBS, lysed in RIPA buffer and scraped. Fluorescence was measured using a

CYTATION3 apparatus. This fluorescence was converted in moles of CPPs using the standard curve presented in panel c. Knowing the cell number, the cell surface area[36] (≈ 1600 $\mu m^2$) and the number of lipid molecules per $\mu m^2$ of plasma membrane (≈$5*10^6$)[37], the moles of lipids in the membrane of the 75'000 HeLa cells used in this experiment could be calculated. The graph reports the CPP to lipid ratio as a function of the CPP concentrations.

**Supplementary Fig. 12: Evidence for low molecular weight pore formation in living cells during CPP direct translocation**

**a**, Representative images related to Fig. 3f. HeLa cells were incubated with 32 μg/ml propidium iodide (PI) in the presence or in the absence of 40 μM FITC-R9, or left untreated for 30 minutes at 37°C in RPMI, 10% FBS. Depolarization was induced with 2 μg/ml gramicidin. Images were obtained by confocal microscopy. As shown below in panel (d) the fluorescence produced by 32 μg/ml of PI is under the threshold of detection. The observed fluorescence dots and signal at this concentration correspond therefore to cell autofluorescence and not PI fluorescence (compare images *i* and *ii*). Hence, when PI fluorescence is detected in cells (see the example in panel a), the corresponding PI concentration is higher than 32 μg/ml (see panel d). In other words, cellular PI fluorescence is detected when cells are able to concentrate this cationic dye in their cytoplasm, as they are able to do with cationic CPP (see Supplementary Fig. 1h). **b**, Criteria used for the visual scoring performed for Fig. 3f, based on images in panel (a) of this figure in HeLa cells. The percentage of cells positive for PI and cells that have acquired FITC-R9 through direct translocation, have been assessed based on the following criteria. Cells 1, 3-6 are considered as positive for PI (left) and cells 1-6 are counted as those where direct translocation has occurred (right). The rest of the cells were considered as negative. Scale bar: 20 μm. **c**, PI does not interact with R9. The potential binding between PI (6 mM) and R9 (0.6 mM) at 37°C was assessed by isothermal titration calorimetry. Representative power and heat of injection are shown. This experiment was repeated three times with similar results. **d**, PI fluorescence detection in media in the absence of cells. Wells containing the indicated PI concentrations in RPMI were illuminated using the same settings as in Fig. 3f and 3g (left panel). Fluorescence intensity within the full region of the obtained images was quantitated using ImageJ. The arrow indicates the PI concentration used for water pore-related experiments (32 μg/ml). **e**, Assessment of colony formation potential after transient CPP and membrane potential treatments. DMSO was used as a vehicle for gramicidin and valinomycin. HeLa cells were incubated for one hour with the indicated treatment, washed and plated on 10 cm dishes. Number of colonies were counted after 14 days in culture after Giemsa staining.

**Supplementary Fig. 13: Zebrafish and mouse membrane potential modulation.**
**a**, Eighteen hours post fertilization zebrafish embryos were incubated for 40 minutes with the indicated concentrations of valinomycin and 950 nM DiBac4(3). DiBac4(3) fluorescence was then recordedand normalized to the non-treated control. The decrease in DiBac4(3) fluorescence indicates membrane hyperpolarization. Membrane potential values could not be calculated, as a standard curve would have to be performed in zebrafish. DiBac4(3) internalization was assessed from confocal images of the fish tail region. **b**, Eighteen hours post fertilization zebrafish embryos were incubated with or without 3.12 μM TAT-RasGAP$_{317-326}$ (W317A), a mutant version that is not toxic to cells, in the presence of the indicated concentrations of valinomycin. Peptide internalization was assessed from confocal images of the fish tail region. **c-d**, Eighteen hours post fertilization zebrafish embryos were incubated 1 hour with the indicated concentrations of valinomycin, in the absence (panel c) or in the presence (panel d) of 3.12 μM TAT-RasGAP$_{317-326}$ (W317A), then washed, and incubated in Egg water. The viability of each fish was assessed over 52 hours at the indicated time points. **e**, Representative images of zebrafish treated as described in panel (c) and (d), washed, and further incubated in Egg water. Images were taken with a CYTATION3 apparatus at a 4x magnification at 70 hours post fertilization (hpf). **f**, Survival of 48 hours post fertilization zebrafish embryos following intramuscular injection of 3.12 μM TAT-RasGAP$_{317-326}$ (W317A) peptide in the presence or in the absence of 10 μM valinomycin. Survival was visually assessed under a binocular microscope by taking into consideration the embryo transparency (as dead embryos appear opaque), development characteristics and motility. **g**, Mice were intradermally injected with DiBac4(3) in the presence or in the absence of 10 μM valinomycin. DiBac4(3) fluorescence was then recorded and normalized to the mean of non-treated (NT) control. DiBac4(3) fluorescence was assessed as in panel (a).

**Supplementary Fig. 14: Model of CPP direct translocation through water pores.**
Cationic CPP translocation across cellular membranes is favored by the opening of potassium channels or by hyperpolarizing drugs, such as valinomycin. This sets a sufficiently low membrane potential permissive for CPP direct translocation. When cationic CPPs bind to these already polarized membranes, they induce megapolarization (i.e. a membrane potential estimated to be -150 mV or lower). This leads to the formation of water pores that are then used by CPPs to enter cells.

**Supplementary Fig. 15: Design of the lipid/water density index collective variables**
**a**, Visual description of the collective variables (c1, c2, and c3) used to compute the lipid/water density index. **b**, Equation used to compute the lipid/water density index for each metadynamics simulation.

**Supplementary Fig. 16: Convergence of the metadynamics simulation of water pore formation in a full DOPC membrane model**
The convergence of the metadynamics simulation was demonstrated by following a well-established computational procedure. **A**, Time evolution of the lipid/water density index. **b**, Gaussian height added to the system. **c**, The free energy difference between low energy states at different times along the simulation was calculated to assess the convergence. The estimated free energy profile is reasonably stable in the last 500 ns of the simulation. The uncertainty, calculated as the SD from the asymptotic value of the free energy obtained from the last part of the simulation, is 1.5 kJ/mol. It is worth mentioning, for this and the next supplementary figures, that the uncertainty does not consider the force-field inaccuracy. The diffusive behavior of the lipid/water density index CV is deduced by checking the ability of the system to overcome the free energy barriers when the gaussian height is closed to zero.

**Supplementary Fig. 17: Convergence of the metadynamics simulation of CPP translocation with a Vm of 0mV**
The convergence of the metadynamics simulation was demonstrated by following a well-established computational procedure. **a**, Time evolution of the CPP-membrane

distance. **b**, Gaussian height added to the system. **c**, The free energy difference between low energy states at different times along the simulation was calculated to assess the convergence. The estimated free energy profile is reasonably stable in the last 500 ns of the simulation. The uncertainty, calculated as the SD from the asymptotic value of the free energy obtained from the last part of the simulation is 1.9 kJ/mol. For this and the next supplementary figures, the diffusive behavior of the CPP-membrane distance CV was deduced by checking the ability of the system to overcome the free energy barriers when the gaussian height is close to zero. **d**, Transmembrane potential computed along the simulation every 10 ns and averaged over 10 windows.

**Supplementary Fig. 18: Convergence of the metadynamics simulation of CPP translocation with a Vm of -80mV**

The convergence of the metadynamics simulation was demonstrated by following a well-established computational procedure. **a**, Time evolution fo the CPP-membrane distance. **b**, Gaussian height added to the system. **c**, The free energy difference between low energy states at different times along the simulation was calculated to assess the convergence. The estimated free energy profile is reasonably stable in the last 500 ns of the simulation. The uncertainty, calculated as the SD from the asymptotic value of the free energy obtained from the last part of the simulation is 3.6 kJ/mol. **d**, Transmembrane potential computed along the simulation every 10 ns and averaged over 10 windows.

**Supplementary Fig. 19: Convergence of the metadynamics simulation of CPP translocation with a Vm of -150mV**

The convergence of the metadynamics simulation was demonstrated by following a well-established computational procedure. **a**, Time evolution of the CPP-membrane distance. **b**, Gaussian height added to the system. **c**, The free energy difference between low energy states at different times along the simulation was calculated to assess the convergence. The estimated free energy profile is reasonably stable in the last 500 ns of the simulation. The uncertainty, calculated as the SD from the asymptotic value of the free energy obtained from the last part of the simulation is 1.9 kJ/mol. **d**, Transmembrane potential computed along the simulation every 10 ns and averaged over 10 windows.

#### Supplementary movies

**Supplementary Movie 1: TAT-RasGAP$_{317\text{-}326}$ internalization in Raji cells over a 16-hour period.**

Representative confocal time-lapse recording of wild-type Raji cells incubated with 5 μM TAT-RasGAP$_{317\text{-}326}$ for 16 hours in RPMI in the absence of serum. For the first 30 minutes of the recording, images were taken every 30 seconds, then until the end of the recording, images were taken every five minutes. Peptide was present in the media throughout the recording. Yellow and pink arrows indicate cells taking up the peptide by direct translocation and by endocytosis, respectively. Cyan arrows point towards labelled endosomes and green asterisks to dead cells. Scale bar: 20 μM. Time is displayed in hours:minutes.

**Supplementary Movie 2: Early peptide entry in wild-type Raji cells.**

Time-lapse recording of Raji cells incubated with 40 μM TAT-RasGAP$_{317\text{-}326}$ for 30 minutes in RPMI, 10% FBS. Peptide was present in the media throughout the recording and images were taken for 30 minutes at 10-second intervals. Scale bar: 10 μM. Time is displayed in minutes:seconds.

**Supplementary Movie 3: Early peptide entry in wild-type HeLa cells**.

Time-lapse recording of HeLa cells incubated with 80 μM FITC-TAT-RasGAP$_{317\text{-}326}$ in RPMI, 10% FBS. Yellow and pink arrows indicate cells experiencing direct translocation and endocytosis, respectively. Images were taken for 30 minutes at 10 second intervals. Scale bar: 20 μM. Time is displayed in minutes:seconds.

**Supplementary Movie 4: Distinction between endosomal escape and direct translocation.**

Wild-type HeLa cells were preincubated with 80 μM FITC-TAT-RasGAP$_{317\text{-}326}$ for 30 minutes in RPMI, 10% FBS and then imaged every 5 minutes for 4 hours at 37°C, 5% $CO_2$. Movie on the left was recorded in the continuous presence of the peptide. Movie on the right was recorded after the peptide was washed three times with RPMI, 10% FBS. Scale bar: 10 μM. Time is displayed in hours:minutes.

**Supplementary Movie 5: *In silico* visualization of water pore formation in the presence of the indicated CPPs across a polarized membrane bilayer.**

This movie shows the translocation of the indicated CPPs across a plasma membrane in the presence of a membrane potential of -2.2 V. This simulation was performed by molecular dynamics MARTINI coarse-grained approach using an asymmetric multi-component bilayer in the presence of ion-imbalance to polarize the membrane.

**Supplementary Movie 6: *In silico* visualization of water pore formation in the presence of the indicated CPPs across a non-polarized membrane bilayer.**

This movie shows the lack of translocation of the indicated CPPs across a plasma membrane in the absence of a membrane potential (0 V). This simulation was performed by molecular dynamics MARTINI coarse-grained approach using an asymmetric multi-component bilayer in the absence of ion-imbalance.

**Supplementary Movie 7: *In silico* visualization of water pore formation in the presence of constant electrical field across a membrane bilayer.**

This movie shows water pore formation in an asymmetric multi-component bilayer in the presence of a membrane potential (no CPP added). This *in silico* simulation was performed with externally applied electric field using molecular dynamics MARTINI coarse-grained approach. In the absence of the $V_m$, interfacial water dipoles predominantly pointed to the hydrophobic core of the membrane partially shielding the phospholipid headgroup dipoles. The application of a ~2V $V_m$ resulted in an enhanced alignment of water dipoles at the membrane outer leaflet and a weakened orientation at the inner side. Also, the lipid headgroups slightly tilted in the direction of the electrostatic field lines. These molecular events increased the probability of the formation of a “nascent pore”, a tiny water wire across the membrane. The insertion of water molecules into the membrane hydrophobic core decreased the free energy barrier for lipid protrusions towards the membrane center and finally resulted in the formation of a metastable hydrophilic membrane pore[29].
